# Early Pastoralism in Central European Forests: Insights from Ancient Environmental Genomics

**DOI:** 10.1101/2023.12.01.569562

**Authors:** Giulia Zampirolo, Luke E. Holman, Rikai Sawafuji, Michaela Ptáková, Lenka Kovačiková, Petr Šída, Petr Pokorný, Mikkel Winther Pedersen, Matthew Walls

## Abstract

Central European forests have been shaped by complex human interactions throughout the Holocene, with significant changes following the introduction of domesticated animals in the Neolithic (∼7.5 – 6.0 kyr BP). However, understanding early pastoral practices and their impact on forests is limited by methods for detecting animal movement across past landscapes. Here we examine ancient sedimentary DNA (*seda*DNA) preserved at the Velký Mamuťák rock shelter, in northern Bohemia (Czech Republic), which has been a forested enclave since the early Holocene. We find that domesticated animals, their associated microbiomes, and plants potentially gathered for fodder, have clear representation by the Late Neolithic, around 6.0 kyr BP, and persist throughout the Bronze Age into recent times. We identify a change in dominant grazing species from sheep to pigs in the Bronze Age (∼4.1 – 3.0 kyr BP) and interpret the impact this had in the mid-Holocene retrogressions that still define the structure of Central European forests today. This study highlights the ability of ancient metagenomics to bridge archaeological and paleoecological methods and provide an enhanced perspective on the roots of the Anthropocene.

## Main text

The emergence of agriculture marks a radical juncture in the entanglement of humans and Earth systems. In Central Europe, the first agricultural sites are associated with the Linearbandkeramik (LBK) peoples, who rapidly occupied fertile lowlands after 7.5 kyr BP, introducing a suite of domesticated plants and animals from the Near East (*1, 2*). Initially, domesticated animals were kept in proximity to agricultural settlements, typically located on loess soils conducive to cultivation. However, the Late Neolithic (sometimes referred to as Eneolithic ∼6.4 – 4.2 kyr BP) witnessed the advent of forest grazing, facilitating a full expansion of pastoralism across varied Central European environments (*3–7*). By the Bronze Age (∼4.1 kyr BP), rich and diverse broadleaf mosaics transitioned into the comparatively species-poor structure that continues to define many Central European forests today (*8–10*). Human influence likely played a significant role in this profound transformation as pastoral activities can impact forest succession (*11*). However, detecting domesticated animal movements across past landscapes is challenging, and the coevolution of forest structure and human agency remains poorly understood.

In this context, the analysis of ancient environmental DNA from sediments offers an opportunity to refine insights into paleoecological changes through time (*12*, *13*) by complementing traditional fossil evidence (*14–16*). Prior attempts to extract ancient DNA from archaeological deposits have utilised target capture techniques to detect hominin and mammal DNA in cave sediments (*17–19*) and rock shelters (*20*, *21*). However, the potential for shotgun sequencing of bulk ancient environmental DNA in a broader range of archaeological settings, such as semi-open to open-air sites, has not yet been fully realised. With this in mind, we identified the Velký Mamuťák (VM) rock shelter as an ideal site for investigation. VM is situated in the Český Ráj region of northern Bohemia, which is a forested area enclosed by sandstone outcrops that create dramatic relief (Fig. 1a). VM is remarkable for its exceptional organic preservation, and deep occupation layers spanning all significant periods from the Mesolithic to the present (Fig 1. b-c). Furthermore, VM benefits from a comprehensive multi-proxy account of environmental practices, derived from prior archaeological and paleoecological examinations (*22*, *23*). At VM, we collected a sequence of sediment samples from stratigraphic layers to examine metagenomic changes spanning the early Holocene to recent times.

**Figure 1.**
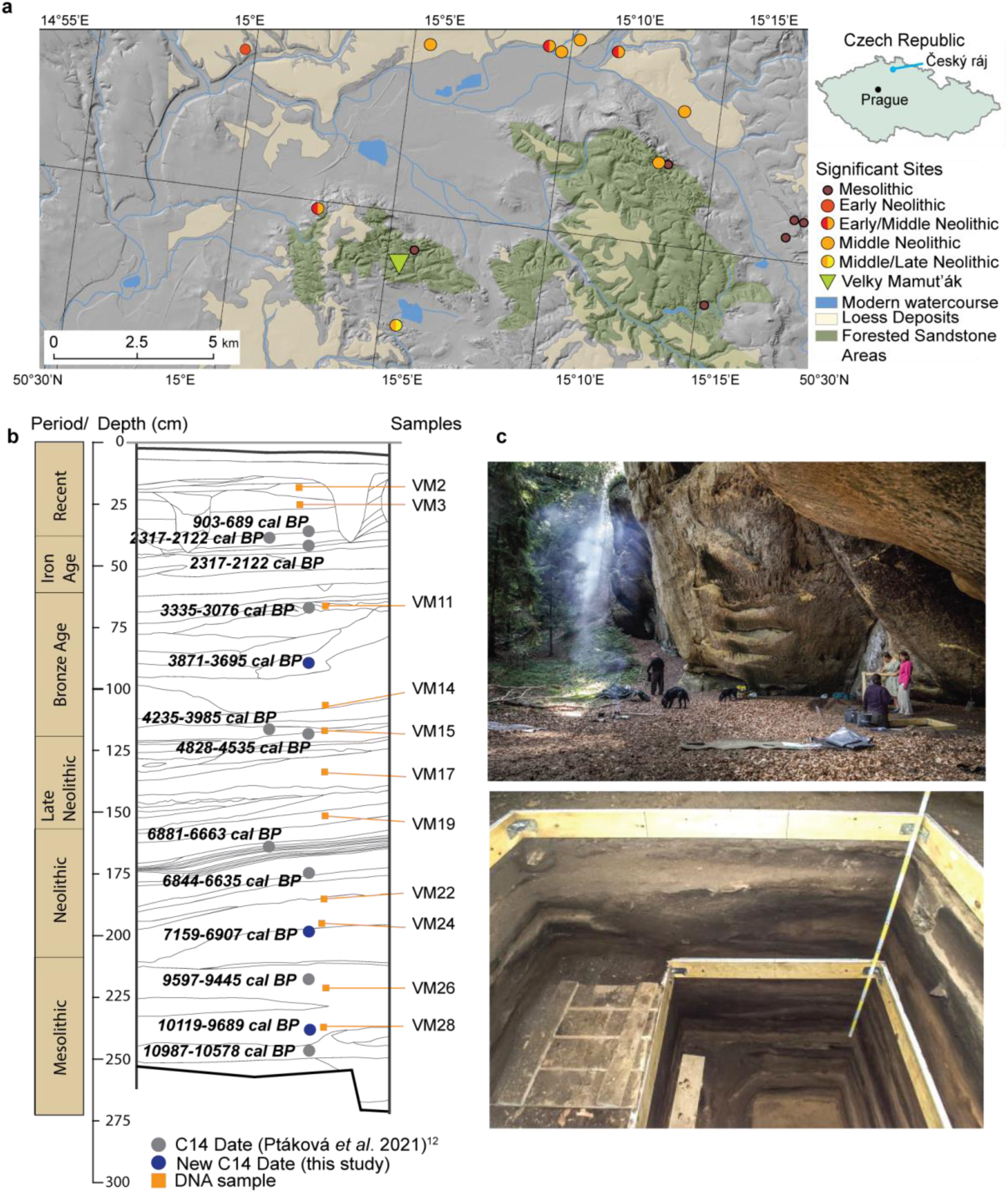
Archaeological setting and age profile. **a**. Location of VM and nearby archaeological sites from Mesolithic to Late Neolithic (Based on geological map 1: 50 000. Adapted by Václav Vondrovský. In: Geological map 1: 50,000 [online]. Praha: Czech Geological Survey [cit. 2022-10-01]. **b**. Sediment profile with radiocarbon ages from a previous study(*22*) (grey) and new dates from this study (blue) together with samples processed for eDNA (orange). **c**. Rock shelter and trench where sediment was sampled.

We identified shifts in domesticated animal DNA at the site from the onset of the Late Neolithic (∼6.0 kyr BP) through the Bronze Age (∼3.0 kyr BP) and show, using phylogenetic placement, that key taxa are ancestral lineages to modern species. In addition, microbial source analysis (*24*, *25*) confirms the presence of bovine species, sheep, and pigs within the same sediment layers, demonstrating the stratigraphic integrity of DNA within the site. We even find evidence of human-associated microbes in a Bronze Age (∼3.0 kyr BP) layer. Furthermore, we track changes in plant DNA which likely corresponds with both grazing practices and wider understandings of environmental retrogressions in this period. Our results at VM support the understanding that the full expansion of herding to forested ecoregions did not take place until the Late Neolithic, with initial management practices focused on sheep (*Ovis*). Importantly, we identify a gradual change in dominant species from sheep to pigs (*Sus*) by the Late Bronze Age. This correlates with the mid-Holocene transformation of forest structure and can potentially be the consequence of this shift in forest succession patterns, nutrient depletion, and habitat connectivity. Overall, our results demonstrate the possibilities for *seda*DNA to explore past human-environment systems and emphasise the value of archaeological deposits as genetic archives for historical ecology.

## Results

### Site, samples, and age-depth model

VM (50°31.10945′ N, 15°4.26233′ E) is located in the Český Ráj region of northern Bohemia (Fig.1a), with a strong history of research on paleoenvironments and prehistoric human settlements in the immediate vicinity (5 km)(*23*, *26–28*). VM is a 450m^2^ area, sheltered by 10 m overhanging cretaceous sandstone and is enclosed in a small canyon with limited drainage (Fig. 1c). Such conditions create a cooled and stable microclimate where sediment accumulated throughout the Holocene. The stratified deposit consists of cultural layers with organic remains producing an extensive assemblage of artefacts and ecofacts. Analyses of charcoal, faunal remains, insect remains, plant macro-remains, malacofauna, pollen, phytoliths, and microcharcoal are reported in prior publications (*22*, *23*). VM yielded abundant well-preserved coprolites from domesticated animals in layers as early as the Late Neolithic (Fig 2b). The presence of stabled animals’ enriched pollen and phytolith contributions to sediments from subsequent periods have been interpreted to outline changes in both the local environment and grazing practices (*23*). However, representation of domesticated animals among recovered archaeozoological remains was generally sparse, likely due to the temporary nature of sheltering events that took place at the site. Given this limitation, *seda*DNA emerged as a promising technique to enhance the fossil record with further insight into species’ presence.

**Figure 2.**
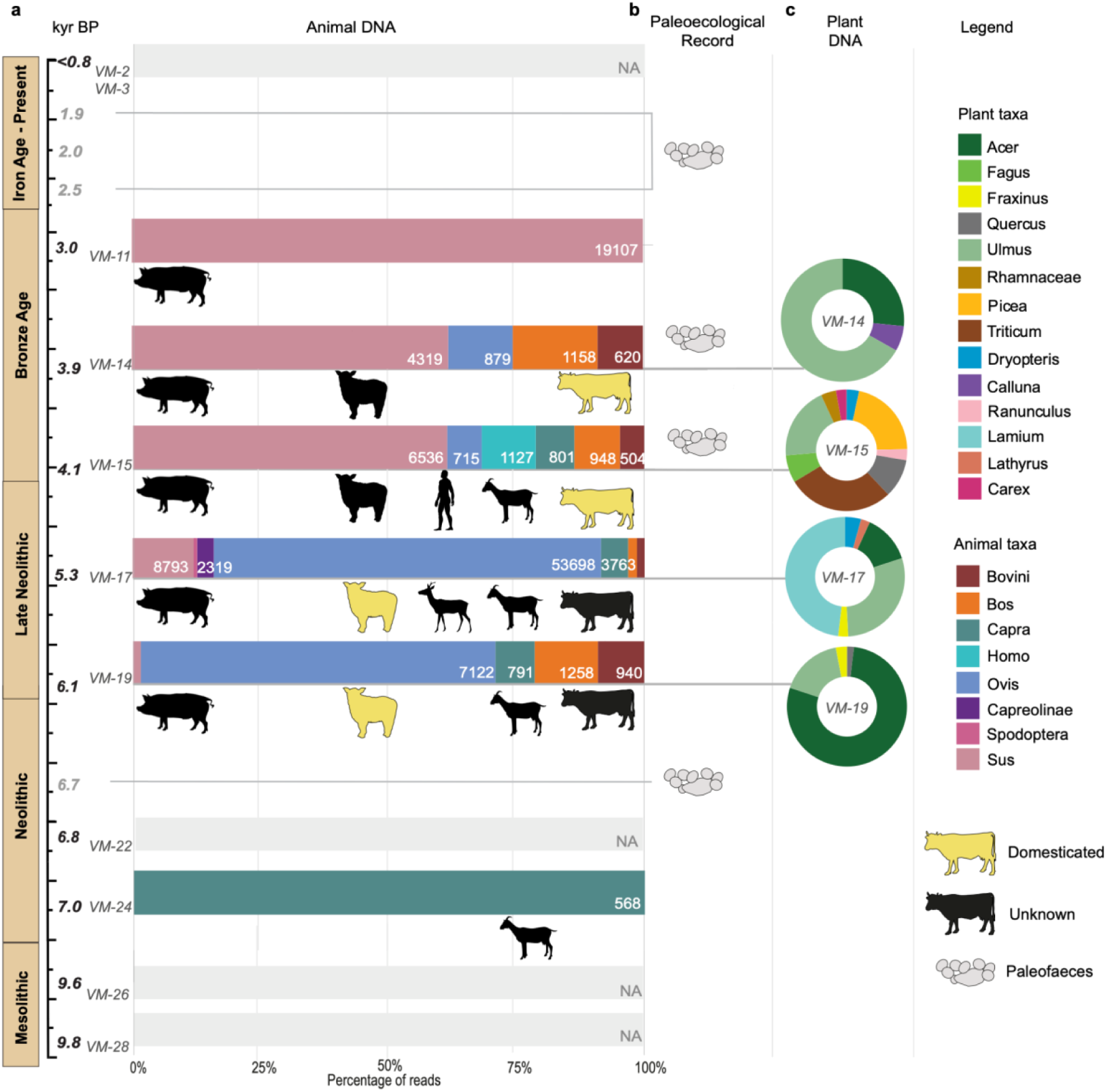
Plant and animal DNA taxonomic profiles. **a.** The relative proportion of DNA reads assigned to animals with total unique reads for each taxon. **b.** The palaeoecological data was reproduced from previously published studies (*22*, *23*). **c**. The relative proportion of DNA reads assigned to plants at the genus level.

Our sampling strategy was therefore built on this prior knowledge assisted by additional ^14^C dating. We collected 28 sediment samples from different occupation layers spanning from the Mesolithic to recent. Of the 28 samples, 22 were extracted for DNA analyses and 11 of these were successfully converted to dual-indexed Illumina libraries (Dataset S1). Challenges in DNA extraction are addressed in the discussion below. Radiocarbon dating of macro botanical remains was used to establish a depositional age-depth model for the entire profile (Supplementary Information S1 and Fig. 1b) with OxCal (v 4.4.4)(*29*) calibrated years BP using the IntCal20(*30*). Overall, the resulting model demonstrated a relatively steady rate of deposition that was beneficial for understanding age ranges represented by the sediment samples (Supplementary Information S1, Fig. S1, Table S1).

### Metagenomic analyses

Libraries were sequenced on an Illumina HiSeq 4000 (80bp single-end) and Illumina Novaseq 6000 (100bp paired-end) platforms obtaining a total of 409,863,279 reads, which were parsed for quality control and removal of low complexity reads and duplicates (see Methods, see Dataset S1). A total of 171,216,656 reads were then parsed through the Holi pipeline for competitive mapping(*13*). Taxonomic profiling and post-mortem DNA damage was estimated using metaDMG(*31*). We used a data-driven filtering approach to investigate and set a minimum threshold for the DNA damage (for details see Methods, Supplementary Information S3), which was applied to obtain the final taxonomic profiles (Fig. 2). In summary, we find post-mortem DNA damage to vary between 5-16% for animals and 5-14% for plants (SI Figs. S7-S9), increasing by depth and age. The five layers of the taxonomic profile, reported as *NA* (VM 2, 3, 22, 26, 28, Fig. 2), fell below the threshold values for authenticated DNA damage, primarily driven by a low significance in the fit to the beta-binomial model used by metaDMG (Zfit < 2) and/or low degree of damage detected (Dfit < 5) (see Methods; SI Figs. S4, S7; Dataset S2). To determine whether the animals found were domestic or wild of origin we extracted all mitochondrial reads at the family level within each of the animal species. For taxa with sufficient reads (> 30), we first performed a phylogenetic placement using BEAST(*32*) and hereafter placed the unique SNPs (single nucleotide polymorphisms) using pathPhynder (*33*) (SI Figs. S10-S12).

### Animal DNA

We find a clear shift and appearance of DNA from domestic animals during the progression of the Neolithic (∼7.0-6.0 kyr BP, see Fig.2). Prior to this, no animal DNA was found in the earliest layers (∼9.9-9.6 kyr BP), and goat or possibly ibex (*Capra sp.*) appears as the only taxon in the Early Neolithic layer (∼7.0 kyr BP). We were unable to classify *Capra* to species level in this layer due to the low number of mitochondrial reads. In the Late Neolithic period (∼6.1-5.3 kyr BP), we record the first appearance of a wider range of animals including cattle or aurochs (*Bos* sp.), pig or wild boar (*Sus* sp.), goat (*Capra hircus*) and sheep (*Ovis aries*). We also find reads classified as wisent (*Bison*) and buffalo (*Bubalus*), however, both species and their geographical distribution in this area remains debated (*34–37*) (*38–40*), and reads could likely be missing or represent past genetic diversity from the taurine cattle found in the same layers. We therefore extracted all reads classified to each genus and blasted these against the nucleotide database (NCBI, RefSeq databases) using blastn (*41*) (see Methods). We find all reads to have the highest E-value across all *Bos* sp., and we therefore report these reads as Bovini. In the same layer, we also detected reads assigned to reindeer (*Rangifer tarandus)*, which we also consider to be an incorrect match, likely due to the absence of a reference genome for the abundant roe deer (*Capreolus capreolus*). We therefore re-mapped the raw sequences to the roe deer reference genome (GCA_000751575.1) to calculate the median edit distance. This reported no difference in the calculation (median = 1), thus the reads classified as reindeer and roe deer are reported as Capreolinae.

During the Bronze Age (∼4.1. kyr BP), we find a composition of animals similar to the Late Neolithic period, with the difference that pig/wild boar increased in abundance. Furthermore, we find ancient human DNA (*Homo sapiens*), which show ancient DNA damage characteristics similar to the other taxa. However, the low coverage and limited number of mitochondrial reads (12 reads, Dataset S3) prevented a further in-depth analysis of haplogroup determination or population placement.

In contrast to the animal diversity in the *seda*DNA record from the earlier periods, the Late Bronze Age (∼3.0 kyr BP) is less diverse with only domestic pig/wild boar present. The uppermost layers are from much more recent periods (Iron Age and later). Here we find domestic pig/wild boar and fly (*Rhagoletis*) but with limited evidence for DNA damage (SI Fig. S4a).

We next phylogenetically placed each consensus mitochondrial genome, where sufficient coverage allowed, onto their respective phylogenetic tree, in order to determine whether the animals were falling closer to domesticates or wild living relatives and counted supporting and conflicting SNPs along each of the branches (see Methods; SI Figs. S10-S12).

We found supporting evidence of domestic alleles for a minimum of one taxon in all layers between 6.1-3.0 kyr BP. In the earliest layer from the Late Neolithic (∼6.1 kyr BP), *Ovis* reads cluster basal to currently living *Ovis aries* haplogroup B and European mouflon (*Ovis aries musimon)*, both domesticated, but with only one supporting SNP (SI Figs. S10a-S11b; Dataset S5). In the younger layer of the late Neolithic (∼5.3 kyr BP) we find strong support (23 SNPs) for a placement within the domestic sheep (*Ovis aries*) branch, with only one conflicting SNP, and the lowest placement within haplogroup B (5 supporting SNPs and 2 conflicting) (SI Figs. S10a-S11a; Dataset S5). Little support (posterior probability = 0.63) was found for the basal placement of the cattle mitochondrial DNA in this layer (SI Fig. S10b), however pathPhynder analysis placed *Bos* reads ancestral to both domestic cattle and aurochs (*5* supporting and 1 conflicting SNPs) (SI Fig. S12c).

The mitochondrial reads of the genus *Bos* from the Early Bronze Age periods (∼4.1-3.9 kyr BP) fall basal to the cattle haplogroups Q and T, but with a low posterior probability between 0.58 and 0.65 (SI Fig. S10b). However, despite this, the SNPs support the Bronze Age (∼4.1 kyr BP) cattle placement as ancestral to the *Bos taurus* haplogroups Q and T (3 supporting SNPs and 2 conflicting ones). It is important to note that the *Bos primigenius* lineage (aurochs) shows a total of 6 conflicting SNPs, indicating that this placement unlikely represents wild species (SI Fig. S12b). Similarly, in the upper layer of the Early Bronze Age (∼3.9 kyr BP), we support the placement of *Bos* reads at the basal node (2 unique SNPs) of domesticated species carrying haplogroup Q and T (SI Fig. S12a). In order to test that the placements were not affected by transitions due to DNA damage, we restricted the analysis to transversions only, for both Early Bronze Age layers, however, this did not change the placement although fewer SNPs were found (SI Fig. S12d).

For several of the other taxa, the available mitochondrial reads were insufficient (<30) to confidently proceed with the phylogenetic investigation (indicated by the label “Unknown” in Fig. 2; see also Dataset S3).

### Plant DNA

We find a relatively low diversity of plants and only 4 samples to yield taxa in the period between the Late Neolithic (∼6.1 kyr BP) and Early Bronze Age (∼3.9 kyr BP). The Late Neolithic was initially (∼6.1 kyr BP) dominated by elm (*Ulmus*) and maple (*Acer*), with a low abundance of ash *(Fraxinus*) and oak (*Quercus*), which likely indicates densely forested conditions. This finding is in agreement with previously published results of pollen and charcoal analyses (*22*, *23*). As the Late Neolithic progressed (∼5.3 kyr BP), we find evidence for the continuous presence of elm and ash trees, including nettles (*Lamium*) as the most abundant taxon. Nettles grow naturally in the forest and are typically available from spring to early summer.

The richest plant diversity we find in the Early Bronze Age (∼4.1-3.9 kyr BP), where elm (*Ulmus*), spruce (*Picea*), and maple (*Acer*) are the dominant tree species, accompanied by oak (*Quercus*) and beech (*Fagus*); leaves of these species also have high nutritional value and historically were favoured fodder(*42*). In this same context, we find a high abundance of wheat (*Triticum*), sedges *(Carex*) that most often is found in wetlands, and heather (*Calluna*) which typically grows on well-drained soils.

### Microbial sources

Lastly, we taxonomically profiled the microbial composition in our samples and compared these to potential source microbiomes to estimate the contributions of each source to each sample. We selected metagenomic sources from different environments including forest, wetland, river sediment, and grassland metagenomes as well as selected mammalian gut and faecal metagenomes. The reads from all samples were mapped, classified and their post-mortem DNA damage estimated (see Methods). We then calculated the proportion of sources using Sourcetracker2(*24*, *43*) both with and without the DNA damage filtering criteria, which were also applied to animal and plant data, and found identical trends. However, filtered data also meant removing the majority of the diversity, most likely due to a low number of reads, hence the proportion without filtering is shown in Fig. 3 (see more detailed discussion in Supplementary Information S5, SI Fig. S15, Dataset S6 and S7).

**Figure 3.**
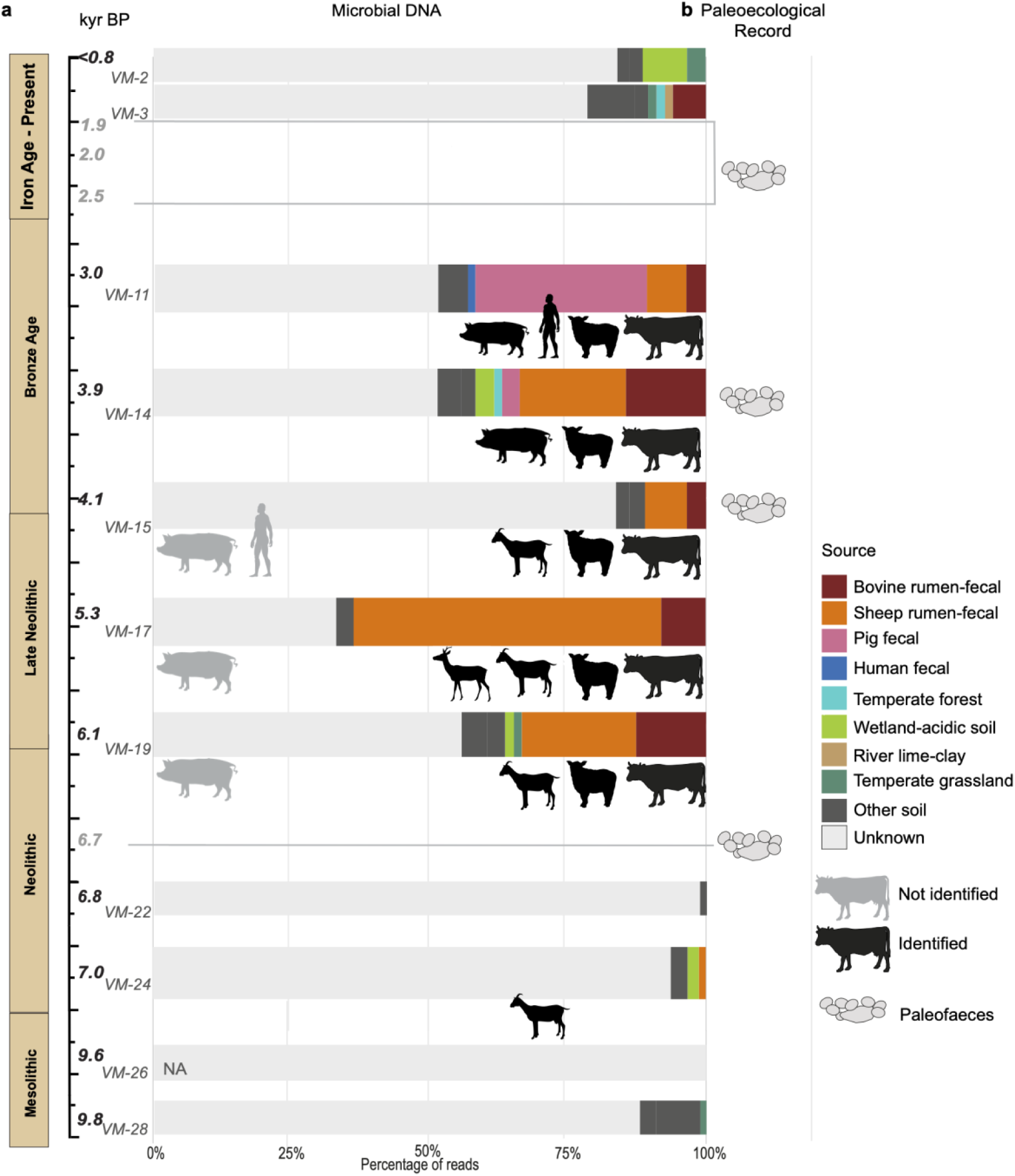
Microbial DNA taxonomic profiles. **a.** The relative proportion of DNA assigned to different sources. **b.** The palaeoecological data was reproduced from a previously published study (*22*). Black outlines represent identified gut metagenomes, while grey outlines represent not identified sources compared to the animal DNA record.

We find the presence of pig, human, sheep, and bovine faecal microbiomes, and that their presence systematically follows the layers in which the animal DNA is also found (Fig. 3). During the pre-and Early Neolithic period the sources are classified primarily as ‘unknown’, which likely reflects that the source soil and sediment microbiomes are not similar to those inhabiting the soil in the rock shelter. A small proportion of bovine rumen-faecal microbiome (1%) are found in the Early Neolithic sample at ∼7.0 kyr BP, which likely derives from the goat/ibex (*Capra* sp.) as found by the animal DNA in this period. A major change occurs during the Late Neolithic (∼6.1-5.3 kyr BP) where a large proportion (77 %) of the microbiome comes from bovine and sheep rumen-faecal matter, a trend that continues up until the Early Bronze Age periods (∼4.1-3.9 kyr BP) where it comprises of 33%. By the late Bronze Age ∼3.0 kyr BP, a change in the metagenome occurs as the pig faecal becomes the dominant microbiome with more than 30% of the metagenome, though we still find a small proportion (10%) of bovine and sheep rumen-faecal microbiota remaining. In the same layer, we also find a low abundance of human faecal microbiome (∼1.3%), which is not the layer where ancient human DNA is found. Representations of the mammalian gut and faecal metagenomes clearly follow the same patterns identified in animal DNA.

Overall, we find only a small proportion of the soil and sedimentary microbial communities to be of wetland-acidic soil, temperate forest, and temperate grassland (1-3%), which is most likely explained by reference source metagenomes being dissimilar to the sediment microbiome.

### DNA taphonomy in the archaeological context

The challenges we encountered during the extraction of the DNA and the conversion to sequencing libraries may be related to several factors. Only 11 of our 22 samples were successfully converted into Illumina libraries and sequenced (Dataset S1). In addition, we found six of the libraries that required a higher number of PCR amplification cycles (> 18 cycles) to exhibit less complexity in plant and animal taxa, which ultimately resulted in high levels of duplication and is likely explained by too few template DNA molecules (Supplementary Information S2, Fig. S2). The lack of template DNA molecules can be connected to several factors such as the amount of deposited material with DNA in the sediments, faster degradation, or localised factors leading to poor preservation in these specific samples. On the other hand, given the organic-rich composition of the sediments (*22*, *23*), inhibiting substances such as humic acids co-extracted during the DNA extractions, despite efforts of removal, may also explain the challenges in both DNA isolation (*44–46*) and eventual downstream molecule preparation (*47*, *48*).

Several lines of evidence demonstrate the DNA we recovered stayed in respective deposited layers through time. If both upward and downward leaching had occurred, the proportion of DNA in the source layer should be larger than in the layers to which it leaches. This does not seem to be the case in this deposit. For example, the youngest layer in the Late Neolithic shows less cattle DNA than the layers both above and below. Early Neolithic caprine DNA was recovered in a single layer without traces above or below. We also find layers beneath the Neolithic period to show the absence of animal DNA, and furthermore, no domesticated animal DNA was detected beneath the Neolithic boundary. Furthermore, using the microbial source tracking, we could authenticate the DNA record, showing that the layers in which we recover animals also show the presence of their respective ruminant and faecal microbiomes in varying abundances which are likely related to the intensity of the presence at the rock shelter.

Accounting for post-depositional taphonomy of DNA molecules is important for the interpretation and hence application of *seda*DNA in archaeological contexts. We expected that bioturbation and/or changes in groundwater level would facilitate some post-depositional movement of metagenomic DNA between some of the stratigraphic layers. Indeed, several studies from other sedimentary deposits have shown that DNA can leach between sediment layers in both caves (*49*) and open-air settings (*50*). We interpret the apparent stratigraphic integrity of our results as an indication that DNA molecules recovered likely were those bound to mineral particles (*51*) or contained within organic structures that do not leach such as micro and macrofossils. We also find cytosine to thymine mis-incorporations due to DNA damage to increase with depth and age (see SI paragraph 3, SI Figs. S8, S9), which we would not expect with the implied mixing of metagenomic material which would take place with post-depositional movement.

## Discussion

Our results complement prior paleoecological analyses from VM and its surroundings by demonstrating a patterned progression in animal, microbial, and plant DNA. While this unique dataset is representative of a single site, it provides valuable insights into broader connections between human practices and environmental outcomes in Central European forests through time. This includes an apparent shift in forest grazing practices during the Bronze Age, which we interpret as a potential factor in the retrogressions of the mid-Holocene (*8*, *9*, *52*). As such, VM underlines the potential of *seda*DNA to open new source deposits for understanding human agency in the postglacial and Holocene succession of Central European forests.

The earliest indication of forest grazing at VM may be the caprine DNA (*Capra* sp.) in association with bovine rumen-faecal microbiome from the Early Neolithic layer (∼7.0 kyr BP). However, further DNA data is required to distinguish domestic goat (*Capra hircus*) from wild species such as ibex (*Capra ibex*), whose occurrence in Český Ráj cannot be fully excluded, despite being undocumented in the fossil record for this period. Clearer genetic representation of domesticated species at VM begins during the Late Neolithic (∼6.1 kyr BP), long after their initial introduction to Central Europe. This supports the perspective of slow environmental neolithization, where the full impact of agriculture across varied ecological zones depended on population expansions associated with Late Neolithic innovations in cultivation and land tenure (*5*, *10*). In a peripheral environment like Český Ráj, it is interesting that DNA from the full range of domesticates - cattle, pigs, goats, and sheep are initially represented; with the exception of a few bone fragments of sheep or goat, these are absent in the archaeozoological assemblage from this period at VM (*22*, *23*). However, despite this range, both animal and microbial DNA demonstrate that sheep were the dominant species in Late Neolithic layers. The phylogenetic analyses allowed us to place cattle (*Bos taurus*) carrying modern haplogroup T and Q, and domestic sheep (*Ovis aries*) carrying haplogroup B, with the species brought to Europe from the Near East by early farmers (*53–59*). However, since cross-breeding between European aurochs and introduced Near Eastern cattle was practised during the Neolithic in Central Europe (*55*, *60–66*), we cannot exclude the presence of wild aurochs at the site. Our identification of Capreolinae is consistent with the faunal records from Late Neolithic archaeological sites elsewhere in Bohemia, where roe deer (*Capreolus capreolus*) was also found (*67*). This continued presence of less abundant wild species suggests ongoing hunting activities in forested environments to supplement grazing.

Český Ráj appears to remain a broadleaf forest mosaic during this stage of Late Neolithic pastoralism (*8*). Sheep are selective grazers and prefer feeding on saplings, shrubs, herbs, and grasses in forest environments (*68*). However, plant DNA from Late Neolithic layers is initially dominated by maple (*Acer*) and elm (*Ulmus*), the leaves of which are high-nutrition fodder, particularly maple (*69*, *70*). This could be a possible indication of seasonality, where animals were brought to the area to graze on select leaf fodder available in the autumn or winter. The later presence of *Lamium* (∼5.3 kyr BP), typically an early seasonal succession weed, is interesting as deadnettle (*Lamium purpureum*) can be palatable but closely related henbit (*Lamium amplexicale*) is toxic for sheep (*42*, *71*). Further investigation of DNA from the Late Neolithic may indicate strong control in grazing combined with adjustment and broadening of seasonality through time. The absence of domesticated crops in both the fossil and genetic records throughout the Late Neolithic could indicate that animals were not crossing cultivated areas while grazing in the vicinity of VM. Overall, this snapshot of Late Neolithic pastoralism is suggestive of a pattern of transhumance, where animals were herded to forested hinterlands during brief seasonal episodes. While further analysis may demonstrate the beginnings of biotic and abiotic impact on forest structure, Late Neolithic pastoralism does not appear to have changed the ecological matrix.

The apparent shift in dominance from sheep to pigs through the Bronze Age (4.1 – 3.0 kyr BP) indicates significant changes in pastoral practices. The co-presence of human DNA and human microbiome, along with the suitability of VM as a shelter, suggests that these pigs were extensively managed. We were unable to phylogenetically place the pig DNA found, due to insufficient mtDNA coverage, however, it would likely not be possible to genetically discern these differences due to the complex lineages of Near Eastern domesticates and local domestications from *Sus scrofa* by around 6.0 kyr BP (*63*, *72*). In any case, pigs are less selective than sheep in forest grazing patterns, and both feral domestic pigs and wild boars remain a key issue in contemporary forest conservation in Europe (*73*). Pigs have the capacity to disrupt normal patterns of succession and pedogenesis through indiscriminate consumption of saplings, herbs, shrubs, grasses, and forest litter. Pigs also practise rooting, which can lead to erosion of soil and nutrient depletion. Indeed, rooting by pigs can significantly impact forest resilience by damaging mycorrhizal networks by replacing symbiotic with pathogenic fungi (*74*).

A corresponding change in woody plant DNA begins in the Early Bronze Age (∼4.1 kyr BP), consisting of taxa such as oak (*Quercus*), beech (*Fagus*), and spruce (*Picea*). These species are less favourable as animal feed due to their low nutritive values (*69*). Wheat (*Triticum*) DNA also appears at the site during this period, together with the presence of other taxa such as sedges (*Carex*) and heather (*Calluna*). Here again, DNA provides extended insight because while wheat is represented in the fossil record at this point by charred grains, sedges, and heather do not appear in the regional pollen or macrofossil record until the start of the Iron Age (∼3.0-1.9 kyr BP) (*9*, *23*). Representation of these species, if deposited with pig faeces, suggests they were crossing cultivated fields and pastures in the vicinity of VM. We interpret this shift as an indication that Český Ráj had by this point become the isolated vestige of forest that it is today, bounded by the sandstone formations that make the area unsuitable for cultivation.

More intensive sampling of *seda*DNA across a range of Neolithic and Bronze Age sites could reveal the timing and sequence of habitat segmentations between Bohemian forests due to expansions in cultivation. These alterations in Bronze Age pastoral practices coincide with the wider mid-Holocene retrogression of Bohemian forest ecosystems, characterised by a shift to comparatively species-poor conditions, as evidenced by pollen records and nutrient depletion in paleosols (*8*, *9*, *75*). While global-scale Quaternary climatic cycles are known to have influenced local factors such as precipitation, impacts of pastoralism may have affected the resilience and capacity of Central European forests to absorb these changes. In areas adjacent to Český Ráj, rapid agricultural expansion through slash-and-burn land conversion is observed in charcoal records, and our results from VM support the interpretation that this resulted in habitat segmentation of forests by the Bronze Age (*11*). No longer used in transhumance, forested islands became a sort of commons predominantly used for grazing pigs; we argue this could be a prime factor in the observed state shift. Expanded studies of *seda*DNA from archaeological and palaeoecological contexts could help confirm a wider range of ecological damage such as the transformation of mycorrhizal networks.

Shotgun sequencing sedimentary ancient DNA demonstrates here its power to enhance archaeological and paleoecological understandings of coupled human-environment systems. At VM, *seda*DNA successfully complements the fossil record to provide a more nuanced representation of species presence through which complex interactions between environmental practice and impact can be assessed. In the context of Central Europe, this can help to further unravel the coevolution of forests and pastoral practices, providing a critical point of reference for understanding forest resilience in the present. Archaeological deposits must be understood as valuable genetic archives for understanding the deep roots of the Anthropocene (*76*).

## Methods

### Sampling

In 2019, we collected a total of 28 sediment samples (∼10 grams), taken directly in the stratigraphic profile, of the trench in sectors B and D (Fig. 1c). We avoided contamination by removing any exposed layer, prior to transferring material to sterile 15-mL centrifuge tubes using sterile disposable scalpels while wearing face masks and nitrile gloves. All samples were hereafter transported to the ancient DNA dedicated laboratories at Globe Institute, University of Copenhagen, Denmark, and stored at -20℃, until subsequent DNA extraction and preparation.

### Radiocarbon dating

Three additional macrofossils were radiocarbon dated, at depths of 89, 198, and 239 cm (SI Table S1), to strengthen the previously published chronology(*22*), now totaling 14 AMS dates. AMS dates were processed at Poznan Radiocarbon Laboratory, Center for Applied Isotope Studies at the University of Georgia, and Beta Analytic (for laboratory codes, and ages see SI Table S1). All dates were calibrated using IntCal20 and used to create an age-depth model in Oxcal (v 4.4.4) (*29*).

### Extraction, library preparation and sequencing

We extracted DNA from a total of 22 sediment samples by subsampling ∼0.25 cm^3^ of sediment into 15 ml sterile spin tubes and adding 5ml of pH 8.0 sodium phosphate buffer (MP Biomedical, Irvine, CA, USA) together with 80ul (18mg/ml) of Proteinase K (Roche Diagnostics, Mannheim, Germany). The reactions were then homogenised using a FastPrep® (MP Biomedicals, Santa Ana, CA, USA) for 2 times 40 seconds at 4.5m/sec and subsequently incubated overnight at room temperature. All samples were hereafter processed by following the Zymo-spin-based protocol (*77*) and 20ul of the extracted DNA (including two negative controls) were then converted to double-stranded Illumina libraries following the standard protocol (*78*). The final reaction was purified with magnetic beads (MagBio HighPrep PCR, MagBio Genomics Inc., USA) at a 1:1.8 ratio. All libraries were equimolarly pooled and sequenced on a HiSeq 4000 80bp single-end or Illumina NovaSeq 6000 100bp paired-end platform at the GeoGenetics Sequencing Core, University of Copenhagen.

### Metagenome analyses

Basecalled and demultiplexed reads were trimmed using AdapterRemoval (v2.3.0)(*79*) and parsed through the ‘Holi’ pipeline (*14*) for quality control, low complexity filtering (≤ 25 bp) and dereplication. The filtered reads were then mapped competitively using Bowtie2 v. 2.3.2 (with options *--k 5000 --no-unal*), against a set of publicly available databases including the non-redundant nt-database (NCBI) as well as the RefSeq database (downloaded 20 December 2020). The alignments were then parsed through metaDMG (version 0.17.5)(*31*) for taxonomic profiling (allowing a similarity between 95%-100% similarity), and DNA damage estimation using the Bayesian fitting model. We explored the output damage statistics and used a data-driven approach to filter for ancient taxa (see Supplementary Information S3, Dataset S2) by requiring each genus to have a minimum level of damage ≥ 5% (*D*_fit_) and a significance ≥ 2 (Z_fit_). We also filtered the concentration for the beta-binomial distribution (*phi*) ≥ 100 and the standard deviation of the damage (*D*_fit_ std) ≤ 0.10. Lastly, key statistics and taxonomic profiles were plotted using R (Supplementary Information S3, SI Figs. S3-S9) and edited with Adobe Illustrator (v. 24.0.3, https://adobe.com/products/illustrator). We particularly tested the reads classified as wisent (*Bison*) and buffalo (*Bubalus*) by extracting their mapped reads at genus level and performing a nucleotide sequence alignment using blastn (v. 2.13.0, last accessed: November 2022) (*41*). Scripts and workflow are available for download here 10.5281/zenodo.7628960.

### Phylogenetic placement

Phylogenetic placement of mitochondrial genomes for relevant animal genuses was performed to distinguish domestic taxa from their wild relatives. First, we performed a preliminary alignment to the NCBI reference mitochondrial genome of sheep (*Ovis* sp.), cattle (*Bos taurus*), aurochs (*Bos primigenius*), domestic pig (*Sus domesticus*), wild boar (*Sus scrofa),* goat/ibex (*Capra* sp.) to verify the abundance of total aligned reads (Dataset S3). As regards the assignment of reads to Capreolinae, we had to remap the sequences to both reindeer (*Rangifer tarandus*, GCA_004026565.1) and roe deer (*Capreolus capreolus*, GCA_000751575.1) separately since the latter was absent in the database used, and we calculated the median of the resulting edit distance in order to determine the best alignment. After this step, we excluded from further phylogenetic analyses all the reads assigned to *Sus* and *Capra.* because of low abundance (Dataset S3). We then downloaded a set of mitochondrial complete genome fasta sequences of sheep and cattle from NCBI’s GenBank and performed a multiple genome alignment with MAFFT (v7.427) reproducing Maximum Likelihood phylogenetic trees (RAXML-NG, v.1.1) from previously published studies (*80*, *81*). We, selected a total of 56 mitogenomes of modern sheep from various breeds of *O. aries, O. ammon, O. vignei, O. orientalis ophion,* and *O. aries musimon*, using a modern *Capra hircus* as outgroup(*80*) (Dataset S4). As for reads assigned to *Bos*, we used a total of 15 mitogenomes from *Bos primigenius, Bos taurus* haplogroups T, Q, and R, using a modern *Bos indicus* as an outgroup (*81*) (Dataset S4). We generated a consensus read for each haplogroup using Geneious (v. 2021.2.2) with the criterion of the majority rules. We extracted the readIDs classified within each genus from metaDMG-lca files following the method described in Kjær et al. (2022)(*13*) and aligned the damaged filtered reads against each consensus sequence separately using Bowtie2 (v. 2.3.2), with a minimum mapping quality of 30. We produced a consensus of the sequences from the bam files with ANGSD (v. 0.928) and an alignment file using MAFFT (v7.427). We performed phylogenetic placement using BEAST (v.1.10.4) with 20,000,000 replicates and a Bayesian inference analysis with the MCMC sampling method, applying the HKY model (SI Fig. S10). Trees were visualised with FIGTREE (v1.4.4). In addition to that, to avoid mapping bias, we re-mapped all the extracted fast reads to the consensus sequence following the method described in Kjær et al. (2022)(*13*) and we retrieved a total of 124 and 343 reads for *Ovis* and *Bos* respectively. Finally, we ran pathPhynder (v. 1.a)(*33*) to identify the unique markers carried by the different haplogroups of sheep and cattle. This algorithm allowed us to place the above-mentioned sequences to branches of a tree based on ancestral and derived SNPs (Single Nucleotide Polymorphisms). We performed both transitions and transversions analysis (*pathPhynder -s all -t 100*) as well as only transversion analysis (*pathPhynder -s all -t 100 -m transversions*), but the latter was not successful in one of our *Bos* alignments (VM-17) and in both *Ovis* alignments (VM-17, VM-19).

### Microbial source tracking

All reads were additionally aligned against the bacterial database GTDB(*82*) release 202 using bowtie2 default end-to-end alignment. DNA damage estimation for each taxonomic level was evaluated using metaDMG with similar settings as above. Source data was downloaded from published microbiome datasets (Dataset S6) and analysed identically with the exception that taxonomic profiles were parsed at the species level to Sourcetracker2(*24*, *43*). With the option --sink_rarefaction_depth 100 to estimate the proportion of each source in sample datasets. Similarly, we parsed the taxonomic profiles with DNA damage (*D*_fit_ ≥ 0.05, Z_fit_ ≥ 2, see SI Fig. S15 and Supplementary Information S5). Lastly, we plotted the proportion of source categories (Dataset S7) per sample and excluded categories with less than 1% proportion.

## Data & code availability

The raw Illumina sequencing data are available from the European Nucleotide Archive under study accession number PRJEB59830. R scripts, metadata, and additional files are permanently archived with the DOI 10.5281/zenodo.7628960.

## Acknowledgments

We acknowledge the support from the GeoGenetics Sequencing Core (University of Copenhagen). All authors also thank Antonio Fernandez-Guerra for help and advice on the microbial analysis.

## Contributions

M.W.P., P.P., and M.W., conceived and planned the idea and experiments. G.Z., and L.E.H., carried out the laboratory work. G.Z., M.W.P., R.S., L.E.H., A.G.F., P.P., and M.W., planned and carried out the data analysis. P.S., P.P., M.P., and L.K., contributed to sample preparation. All authors contributed to the interpretation of the results. G.Z., M.W.P., and M.W., took the lead in writing the manuscript. All authors provided feedback and helped shape the research, analysis and manuscript.

## Ethical declarations

The authors declare no competing interests.

## List of links for silhouettes used in Fig. 2 and Fig. 3

Sus: http://phylopic.org/image/008d6d88-d1be-470a-8c70-73625c3fb4fb/

Bos: http://phylopic.org/image/415714b4-859c-4d1c-9ce0-9e1081613df7/

Ovis: http://www.freepik.com ; Designed by ilonitta / Freepik

Capra: http://phylopic.org/image/c39a1d31-eb36-4b62-ba4d-32e29e0e5a8d/

Capreolinae: http://phylopic.org/image/b36a215a-adb3-445d-b364-1e63dddd6950/

Homo: http://phylopic.org/image/c089caae-43ef-4e4e-bf26-973dd4cb65c5/

## References

1. J. Jakucs, E. Bánffy, K. Oross, V. Voicsek, C. Bronk Ramsey, E. Dunbar, B. Kromer, A. Bayliss, D. Hofmann, P. Marshall, A. Whittle, Between the Vinča and Linearbandkeramik Worlds: The Diversity of Practices and Identities in the 54th-53rd Centuries cal BC in Southwest Hungary and Beyond. J World Prehist. 29, 267–336 (2016).

2. S. Shennan, The First Farmers of Europe: An Evolutionary Perspective (Cambridge University Press, 2018).

3. R. Berthon, L. Kovačiková, A. Tresset, M. Balasse, Integration of Linearbandkeramik cattle husbandry in the forested landscape of the mid-Holocene climate optimum: Seasonal-scale investigations in Bohemia. Journal of Anthropological Archaeology. 51, 16–27 (2018).

4. A. Bogaard, Questioning the relevance of shifting cultivation to Neolithic farming in the loess belt of Europe: evidence from the Hambach Forest experiment. Vegetation history and archaeobotany. 11, 155–168 (2002).

5. J. Kolář, M. Macek, P. Tkáč, D. Novák, V. Abraham, Long-term demographic trends and spatio-temporal distribution of past human activity in Central Europe: Comparison of archaeological and palaeoecological proxies. Quaternary science reviews. 297, 107834 (2022).

6. M. Saqalli, A. Salavert, S. Bréhard, R. Bendrey, J.-D. Vigne, A. Tresset, Revisiting and modelling the woodland farming system of the early Neolithic Linear Pottery Culture (LBK), 5600–4900 b.c. Vegetation history and archaeobotany. 23, 37–50 (2014).

7. R. Gillis, I. Kendall, M. Roffet-Salque, M. Zanon, A. Anders, R.-M. Arbogast, P. Bogucki, V. Brychova, E. Casanova, E. Claßen, P. Csengeri, L. Czerniak, L. Domboróczki, D. Fiorillo, D. Gronenborn, L. Hachem, J. Jakucs, M. Ilet, K. Lyublyanovics, E. Lenneis, A. Marciniak, T. Marton, K. Oross, J. Pavúk, J. Pechtl, J. Pyzel, P. Stadler, H. Stäuble, I. Vostrovská, I. van Wijk, Vigne, M. Balasse, R. Evershed, Forest ecosystems and evolution of cattle husbandry practices of the earliest central European farming societies. Research Square (2022), doi:10.21203/rs.3.rs-1419935/v1.

8. L. Juřičková, P. Šída, J. Horáčková, V. Ložek, P. Pokorný, The lost paradise of snails: Transformation of the middle-Holocene forest ecosystems in Bohemia, Czech Republic, as revealed by declining land snail diversity. Holocene. 30, 1254–1265 (2020).

9. J. Novák, V. Abraham, P. Šída, P. Pokorný, Holocene forest transformations in sandstone landscapes of the Czech Republic: Stand-scale comparison of charcoal and pollen records. The Holocene. 29, 1468–1479 (2019).

10. M. Ptáková, P. Šída, V. Vondrovský, P. Pokorný, Islands of Difference: An Ecologically Explicit Model of Central European Neolithisation. Environ. Archaeol., 1–9 (2021).

11. P. Bobek, H. S. Svobodová, B. Werchan, M. G. Švarcová, P. Kuneš, Human-induced changes in fire regime and subsequent alteration of the sandstone landscape of Northern Bohemia (Czech Republic). The Holocene. 28, 427–443 (2018).

12. L. Parducci, I. G. Alsos, P. Unneberg, M. W. Pedersen, L. Han, Y. Lammers, J. S. Salonen, M. M. Väliranta, T. Slotte, B. Wohlfarth, Shotgun Environmental DNA, Pollen, and Macrofossil Analysis of Lateglacial Lake Sediments From Southern Sweden. Frontiers in Ecology and Evolution. 7 (2019), doi:10.3389/fevo.2019.00189.

13. K. H. Kjær, M. Winther Pedersen, B. De Sanctis, B. De Cahsan, T. S. Korneliussen, C. S. Michelsen, K. K. Sand, S. Jelavić, A. H. Ruter, A. M. A. Schmidt, K. K. Kjeldsen, A. S. Tesakov, I. Snowball, J. C. Gosse, I. G. Alsos, Y. Wang, C. Dockter, M. Rasmussen, M. E. Jørgensen, B. Skadhauge, A. Prohaska, J. Å. Kristensen, M. Bjerager, M. E. Allentoft, E. Coissac, PhyloNorway Consortium, A. Rouillard, A. Simakova, A. Fernandez-Guerra, C. Bowler, M. Macias-Fauria, L. Vinner, J. J. Welch, A. J. Hidy, M. Sikora, M. J. Collins, R. Durbin, N. K. Larsen, E. Willerslev, A 2-million-year-old ecosystem in Greenland uncovered by environmental DNA. Nature. 612, 283–291 (2022).

14. M. W. Pedersen, B. De Sanctis, N. F. Saremi, M. Sikora, E. E. Puckett, Z. Gu, K. L. Moon, J. D. Kapp, L. Vinner, Z. Vardanyan, C. F. Ardelean, J. Arroyo-Cabrales, J. A. Cahill, P. D. Heintzman, G. Zazula, R. D. E. MacPhee, B. Shapiro, R. Durbin, E. Willerslev, Environmental genomics of Late Pleistocene black bears and giant short-faced bears. Curr. Biol. 31, 2728–2736.e8 (2021).

15. F. V. Seersholm, M. W. Pedersen, M. J. Søe, H. Shokry, S. S. T. Mak, A. Ruter, M. Raghavan, W. Fitzhugh, K. H. Kjær, E. Willerslev, M. Meldgaard, C. M. O. Kapel, A. J. Hansen, DNA evidence of bowhead whale exploitation by Greenlandic Paleo-Inuit 4,000 years ago. Nat. Commun. 7, 13389 (2016).

16. C. F. Ardelean, L. Becerra-Valdivia, M. W. Pedersen, J.-L. Schwenninger, C. G. Oviatt, J. I. Macías-Quintero, J. Arroyo-Cabrales, M. Sikora, Y. Z. E. Ocampo-Díaz, I. I. Rubio-Cisneros, J. G. Watling, V. B. de Medeiros, P. E. De Oliveira, L. Barba-Pingarón, A. Ortiz-Butrón, J. Blancas-Vázquez, I. Rivera-González, C. Solís-Rosales, M. Rodríguez-Ceja, D. A. Gandy, Z. Navarro-Gutierrez, J. J. De La Rosa-Díaz, V. Huerta-Arellano, M. B. Marroquín-Fernández, L. M. Martínez-Riojas, A. López-Jiménez, T. Higham, E. Willerslev, Evidence of human occupation in Mexico around the Last Glacial Maximum. Nature. 584, 87–92 (2020).

17. D. Massilani, M. W. Morley, S. M. Mentzer, V. Aldeias, B. Vernot, C. Miller, M. Stahlschmidt, M. B. Kozlikin, M. V. Shunkov, A. P. Derevianko, N. J. Conard, S. Wurz, C. S. Henshilwood, J. Vasquez, E. Essel, S. Nagel, J. Richter, B. Nickel, R. G. Roberts, S. Pääbo, V. Slon, P. Goldberg, M. Meyer, Microstratigraphic preservation of ancient faunal and hominin DNA in Pleistocene cave sediments. Proc. Natl. Acad. Sci. U. S. A. 119 (2022), doi:10.1073/pnas.2113666118.

18. B. Vernot, E. I. Zavala, A. Gómez-Olivencia, Z. Jacobs, V. Slon, F. Mafessoni, F. Romagné, A. Pearson, M. Petr, N. Sala, A. Pablos, A. Aranburu, J. M. B. de Castro, E. Carbonell, B. Li, M. T. Krajcarz, A. I. Krivoshapkin, K. A. Kolobova, M. B. Kozlikin, M. V. Shunkov, A. P. Derevianko, B. Viola, S. Grote, E. Essel, D. L. Herráez, S. Nagel, B. Nickel, J. Richter, A. Schmidt, B. Peter, J. Kelso, R. G. Roberts, J.-L. Arsuaga, M. Meyer, Unearthing Neanderthal population history using nuclear and mitochondrial DNA from cave sediments. Science. 372 (2021), doi:10.1126/science.abf1667.

19. V. Slon, C. Hopfe, C. L. Weiß, F. Mafessoni, M. de la Rasilla, C. Lalueza-Fox, A. Rosas, M. Soressi, M. V. Knul, R. Miller, J. R. Stewart, A. P. Derevianko, Z. Jacobs, B. Li, R. G. Roberts, M. V. Shunkov, H. de Lumley, C. Perrenoud, I. Gušić, Ž. Kućan, P. Rudan, A. Aximu-Petri, E. Essel, S. Nagel, B. Nickel, A. Schmidt, K. Prüfer, J. Kelso, H. A. Burbano, S. Pääbo, M. Meyer, Neandertal and Denisovan DNA from Pleistocene sediments. Science. 356, 605–608 (2017).

20. S. Silvestrini, M. Romandini, G. Marciani, S. Arrighi, L. Carrera, A. Fiorini, J. M. López-García, F. Lugli, F. Ranaldo, V. Slon, L. Tassoni, O. A. Higgins, E. Bortolini, A. Curci, M. Meyer, M. C. Meyer, G. Oxilia, A. Zerboni, S. Benazzi, E. E. Spinapolice, Integrated multidisciplinary ecological analysis from the Uluzzian settlement at the Uluzzo C Rock Shelter, south-eastern Italy. J. Quat. Sci. 37, 235–256 (2022).

21. F. Braadbaart, F. H. Reidsma, W. Roebroeks, L. Chiotti, V. Slon, M. Meyer, I. Théry-Parisot, A. van Hoesel, K. G. J. Nierop, J. Kaal, B. van Os, L. Marquer, Heating histories and taphonomy of ancient fireplaces: A multi-proxy case study from the Upper Palaeolithic sequence of Abri Pataud (Les Eyzies-de-Tayac, France). Journal of Archaeological Science: Reports. 33, 102468 (2020).

22. M. Ptáková, P. Pokorný, P. Šída, J. Novák, I. Horáček, L. Juřičková, P. Meduna, A. Bezděk, E. Myšková, M. Walls, P. Poschlod, From Mesolithic hunters to Iron Age herders: a unique record of woodland use from eastern central Europe (Czech Republic). Vegetation history and archaeobotany. 30, 269–286 (2021).

23. P. Šída, P. Pokorný, Mezolit Severních Čech III: Vývoj Pravěké Krajiny Českého Ráje: Vegetace, Fauna, Lidé (Archeologický ústav AV ČR, 2020).

24. D. Knights, J. Kuczynski, E. S. Charlson, J. Zaneveld, M. C. Mozer, R. G. Collman, F. D. Bushman, R. Knight, S. T. Kelley, Bayesian community-wide culture-independent microbial source tracking. Nat. Methods. 8, 761–763 (2011).

25. M. C. Wibowo, Z. Yang, M. Borry, A. Hübner, K. D. Huang, B. T. Tierney, S. Zimmerman, F. Barajas-Olmos, C. Contreras-Cubas, H. García-Ortiz, A. Martínez-Hernández, J. M. Luber, P. Kirstahler, T. Blohm, F. E. Smiley, R. Arnold, S. A. Ballal, S. J. Pamp, J. Russ, F. Maixner, O. Rota-Stabelli, N. Segata, K. Reinhard, L. Orozco, C. Warinner, M. Snow, S. LeBlanc, A. D. Kostic, Reconstruction of ancient microbial genomes from the human gut. Nature. 594, 234–239 (2021).

26. 26. J. Svoboda, P. Pokorný, I. Horáček, S. Sázelová, V. Abraham, M. Divišová, M. Ivanov, R. Kozáková, J. Novák, M. Novák, P. Šída, A. Perri, Late Glacial and Holocene sequences in rockshelters and adjacent wetlands of Northern Bohemia, Czech Republic: Correlation of environmental and archaeological records. Quat. Int. 465, 234–250 (2018).

27. J. Novák, J. Svoboda, P. Šída, J. Prostředník, P. Pokorný, A charcoal record of Holocene woodland succession from sandstone rock shelters of North Bohemia (Czech Republic). Quat. Int. 366, 25–36 (2015).

28. J. Svoboda, Mezolit Severnich Cech II: Komplexní výzkum skalních převisů na Českolipsku a Děčínsku, 2003–2015 (Vol. v.v.i.) (Archeologický ústav AV CR, Brno, 2017).

29. C. B. Ramsey, Bayesian Analysis of Radiocarbon Dates. Radiocarbon. 51, 337–360 (2009).

30. P. J. Reimer, W. E. N. Austin, E. Bard, A. Bayliss, P. G. Blackwell, C. B. Ramsey, M. Butzin, H. Cheng, R. Lawrence Edwards, M. Friedrich, P. M. Grootes, T. P. Guilderson, I. Hajdas, T. J. Heaton, A. G. Hogg, K. A. Hughen, B. Kromer, S. W. Manning, R. Muscheler, J. G. Palmer, C. Pearson, J. van der Plicht, R. W. Reimer, D. A. Richards, E. Marian Scott, J. R. Southon, C. S. M. Turney, L. Wacker, F. Adolphi, U. Büntgen, M. Capano, S. M. Fahrni, A. Fogtmann-Schulz, R. Friedrich, P. Köhler, S. Kudsk, F. Miyake, J. Olsen, F. Reinig, M. Sakamoto, A. Sookdeo, S. Talamo, The IntCal20 Northern Hemisphere Radiocarbon Age Calibration Curve (0–55 cal kBP). Radiocarbon. 62, 725–757 (2020).

31. C. Michelsen, M. W. Pedersen, A. Fernandez-Guerra, L. Zhao, T. C. Petersen, T. S. Korneliussen, metaDMG – A Fast and Accurate Ancient DNA Damage Toolkit for Metagenomic Data. https://www.biorxiv.org/content/10.1101/2022.12.06.519264v1 (2022), doi:10.1101/2022.12.06.519264.

32. A. J. Drummond, A. Rambaut, BEAST: Bayesian evolutionary analysis by sampling trees. BMC Evol. Biol. 7, 214 (2007).

33. R. Martiniano, B. De Sanctis, P. Hallast, R. Durbin, Placing Ancient DNA Sequences into Reference Phylogenies. Molecular biology and evolution. 39, msac017 (2022).

34. J. A. Lenstra, J. Liu, The Year of the Wisent. BMC Biol. 14 (2016), p. 100.

35. D. Massilani, S. Guimaraes, J.-P. Brugal, E. A. Bennett, M. Tokarska, R.-M. Arbogast, G. Baryshnikov, G. Boeskorov, J.-C. Castel, S. Davydov, S. Madelaine, O. Putelat, N. N. Spasskaya, H.-P. Uerpmann, T. Grange, E.-M. Geigl, Past climate changes, population dynamics and the origin of Bison in Europe. BMC Biol. 14, 93 (2016).

36. J. Soubrier, G. Gower, K. Chen, S. M. Richards, B. Llamas, K. J. Mitchell, S. Y. W. Ho, P. Kosintsev, M. S. Y. Lee, G. Baryshnikov, R. Bollongino, P. Bover, J. Burger, D. Chivall, E. Crégut-Bonnoure, J. E. Decker, V. B. Doronichev, K. Douka, D. A. Fordham, F. Fontana, C. Fritz, J. Glimmerveen, L. V. Golovanova, C. Groves, A. Guerreschi, W. Haak, T. Higham, E. Hofman-Kamińska, A. Immel, M.-A. Julien, J. Krause, O. Krotova, F. Langbein, G. Larson, A. Rohrlach, A. Scheu, R. D. Schnabel, J. F. Taylor, M. Tokarska, G. Tosello, J. van der Plicht, A. van Loenen, J.-D. Vigne, O. Wooley, L. Orlando, R. Kowalczyk, B. Shapiro, A. Cooper, Early cave art and ancient DNA record the origin of European bison. Nat. Commun. 7, 13158 (2016).

37. N. Navani, P. K. Jain, S. Gupta, B. S. Sisodia, S. Kumar, A set of cattle microsatellite DNA markers for genome analysis of riverine buffalo (Bubalus bubalis). Anim. Genet. 33, 149–154 (2002).

38. A. Noce, S. Qanbari, R. González-Prendes, J. Brenmoehl, M. G. Luigi-Sierra, M. Theerkorn, M.-A. Fiege, H. Pilz, A. Bota, L. Vidu, C. Horwath, L. Haraszthy, P. Penchev, Y. Ilieva, T. Peeva, W. Lüpcke, R. Krawczynski, K. Wimmers, M. Thiele, A. Hoeflich, Genetic Diversity of Bubalus bubalis in Germany and Global Relations of Its Genetic Background. Front. Genet. 11, 610353 (2020).

39. R. Kysely, The palaeoeconomy of the Bohemian and Moravian Lengyel and Eneolithic periods from the perspective of archaeozoology. Památky Archeologické. 103, 5–70 (2012).

40. A. Németh, A. Bárány, G. Csorba, E. Magyari, P. Pazonyi, J. Pálfy, Holocene mammal extinctions in the Carpathian Basin: a review. Mammal Review. 47, 38–52 (2017).

41. S. F. Altschul, T. L. Madden, A. A. Schäffer, J. Zhang, Z. Zhang, W. Miller, D. J. Lipman, Gapped BLAST and PSI-BLAST: a new generation of protein database search programs. Nucleic Acids Res. 25, 3389–3402 (1997).

42. A. Kirilov, N. Georgieva, I. Stoycheva, Determination of composition and palatability of certain weeds. International Journal of Agricultural Science and Food Technology. 2, 41–43 (2016).

43. J. J. McGhee, N. Rawson, B. A. Bailey, A. Fernandez-Guerra, L. Sisk-Hackworth, S. T. Kelley, Meta-SourceTracker: application of Bayesian source tracking to shotgun metagenomics. PeerJ. 8, e8783 (2020).

44. G. Pietramellara, J. Ascher, F. Borgogni, M. T. Ceccherini, G. Guerri, P. Nannipieri, Extracellular DNA in soil and sediment: fate and ecological relevance. Biol. Fertil. Soils. 45, 219–235 (2009).

45. A. Wolińska, Z. Stępniewska, Dehydrogenase activity in the soil environment. Dehydrogenases. 10, 183–210 (2012).

46. E. Wnuk, A. Waśko, A. Walkiewicz, P. Bartmiński, R. Bejger, L. Mielnik, A. Bieganowski, The effects of humic substances on DNA isolation from soils. PeerJ. 8, e9378 (2020).

47. S. Köchl, H. Niederstätter, W. Parson, DNA extraction and quantitation of forensic samples using the phenol-chloroform method and real-time PCR. Methods Mol. Biol. 297, 13–30 (2005).

48. T. Demeke, G. R. Jenkins, Influence of DNA extraction methods, PCR inhibitors and quantification methods on real-time PCR assay of biotechnology-derived traits. Anal. Bioanal. Chem. 396, 1977–1990 (2010).

49. J. Haile, R. Holdaway, K. Oliver, M. Bunce, M. T. P. Gilbert, R. Nielsen, K. Munch, S. Y. W. Ho, B. Shapiro, E. Willerslev, Ancient DNA chronology within sediment deposits: are paleobiological reconstructions possible and is DNA leaching a factor? Mol. Biol. Evol. 24, 982–989 (2007).

50. K. Andersen, K. L. Bird, M. Rasmussen, J. Haile, H. Breuning-Madsen, K. H. Kjaer, L. Orlando, M. T. P. Gilbert, E. Willerslev, Meta-barcoding of “dirt” DNA from soil reflects vertebrate biodiversity. Mol. Ecol. 21, 1966–1979 (2012).

51. C. L. Freeman, L. Dieudonné, O. B. A. Agbaje, M. Zure, J. Q. Sanz, M. Collins, K. K. Sand, Survival of environmental DNA in sediments: Mineralogic control on DNA taphonomy. https://www.biorxiv.org/content/10.1101/2020.01.28.922997v2 (2023), doi:10.1101/2020.01.28.922997.

52. 52. V. Ložek, “Late Bronze Age environmental collapse in the sandstone areas of northern Bohemia” in Mensch und Umwelt in der Bronzezeit Europas, B. Hansel, Ed. (Oetker-Voges Verlag, Kiel, 1998), pp. 57–60.

53. A. Achilli, S. Bonfiglio, A. Olivieri, A. Malusà, M. Pala, B. Hooshiar Kashani, U. A. Perego, P. Ajmone-Marsan, L. Liotta, O. Semino, H.-J. Bandelt, L. Ferretti, A. Torroni, The multifaceted origin of taurine cattle reflected by the mitochondrial genome. PLoS One. 4, e5753 (2009).

54. A. Olivieri, F. Gandini, A. Achilli, A. Fichera, E. Rizzi, S. Bonfiglio, V. Battaglia, S. Brandini, A. De Gaetano, A. El-Beltagi, H. Lancioni, S. Agha, O. Semino, L. Ferretti, A. Torroni, Mitogenomes from Egyptian Cattle Breeds: New Clues on the Origin of Haplogroup Q and the Early Spread of Bos taurus from the Near East. PLoS One. 10, e0141170 (2015).

55. R. Kyselý, M. Hájek, MtDNA haplotype identification of aurochs remains originating from the Czech Republic (Central Europe). Environ. Archaeol. 17, 118–125 (2012).

56. 56. E. Nikulina, U. Schmölcke, “The first genetic evidence for the origin of central European sheep (Ovis ammon f. aries) populations from two different routes of Neolithisation and contributions to the history of woolly sheep” in The competition of fibres : Early textile production in western Asia, south-east and central Europe (10,000-500BCE), S. Pollock, W. & Schier, Eds. (Oxbow Books, Oxford, England, 2020), pp. 203–210.

57. M. Tapio, N. Marzanov, M. Ozerov, M. Cinkulov, G. Gonzarenko, T. Kiselyova, M. Murawski, H. Viinalass, J. Kantanen, Sheep mitochondrial DNA variation in European, Caucasian, and Central Asian areas. Mol. Biol. Evol. 23, 1776–1783 (2006).

58. J. R. S. Meadows, I. Cemal, O. Karaca, E. Gootwine, J. W. Kijas, Five ovine mitochondrial lineages identified from sheep breeds of the near East. Genetics. 175, 1371–1379 (2007).

59. J. R. S. Meadows, S. Hiendleder, J. W. Kijas, Haplogroup relationships between domestic and wild sheep resolved using a mitogenome panel. Heredity. 106, 700–706 (2011).

60. A. Achilli, A. Olivieri, M. Pellecchia, C. Uboldi, L. Colli, N. Al-Zahery, M. Accetturo, M. Pala, B. Hooshiar Kashani, U. A. Perego, V. Battaglia, S. Fornarino, J. Kalamati, M. Houshmand, R. Negrini, O. Semino, M. Richards, V. Macaulay, L. Ferretti, H.-J. Bandelt, P. Ajmone-Marsan, A. Torroni, Mitochondrial genomes of extinct aurochs survive in domestic cattle. Curr. Biol. 18, R157–8 (2008).

61. A. Götherström, C. Anderung, L. Hellborg, R. Elburg, C. Smith, D. G. Bradley, H. Ellegren, Cattle domestication in the Near East was followed by hybridization with aurochs bulls in Europe. Proc. Biol. Sci. 272, 2345–2350 (2005).

62. J. Schibler, J. Elsner, A. Schlumbaum, Incorporation of aurochs into a cattle herd in Neolithic Europe: single event or breeding? Scientific Reports. 4, 1–6 (2014).

63. 63. R. Kyselý, The size of domestic cattle, sheep, goats and pigs in the Czech Neolithic and Eneolithic Periods: Temporal variations and their causes (2016), (available at https://revistas.uam.es/archaeofauna/article/download/6358/6831/12813).

64. C. J. Edwards, R. Bollongino, A. Scheu, A. Chamberlain, A. Tresset, J.-D. Vigne, J. F. Baird, G. Larson, S. Y. W. Ho, T. H. Heupink, B. Shapiro, A. R. Freeman, M. G. Thomas, R.-M. Arbogast, B. Arndt, L. Bartosiewicz, N. Benecke, M. Budja, L. Chaix, A. M. Choyke, E. Coqueugniot, H.-J. Döhle, H. Göldner, S. Hartz, D. Helmer, B. Herzig, H. Hongo, M. Mashkour, M. Ozdogan, E. Pucher, G. Roth, S. Schade-Lindig, U. Schmölcke, R. J. Schulting, E. Stephan, H.-P. Uerpmann, I. Vörös, B. Voytek, D. G. Bradley, J. Burger, Mitochondrial DNA analysis shows a Near Eastern Neolithic origin for domestic cattle and no indication of domestication of European aurochs. Proc. Biol. Sci. 274, 1377–1385 (2007).

65. A. Beja-Pereira, D. Caramelli, C. Lalueza-Fox, C. Vernesi, N. Ferrand, A. Casoli, F. Goyache, L. J. Royo, S. Conti, M. Lari, A. Martini, L. Ouragh, A. Magid, A. Atash, A. Zsolnai, P. Boscato, C. Triantaphylidis, K. Ploumi, L. Sineo, F. Mallegni, P. Taberlet, G. Erhardt, L. Sampietro, J. Bertranpetit, G. Barbujani, G. Luikart, G. Bertorelle, The origin of European cattle: evidence from modern and ancient DNA. Proc. Natl. Acad. Sci. U. S. A. 103, 8113–8118 (2006).

66. S. Bonfiglio, A. Achilli, A. Olivieri, R. Negrini, L. Colli, L. Liotta, P. Ajmone-Marsan, A. Torroni, L. Ferretti, The enigmatic origin of bovine mtDNA haplogroup R: sporadic interbreeding or an independent event of Bos primigenius domestication in Italy? PLoS One. 5, e15760 (2010).

67. L. Kovačikova, S. Bréhard, R. Šumberová, M. Balasse, A. Tresset, The new insights into the subsistence and early farming from Neolithic settlements in Central Europe: the archaeozoological evidence from the Czech Republic. Archaeofauna. 21 (2012) (available at https://www.researchgate.net/profile/Anne-Tresset/publication/235898345_The_new_insights_into_the_subsistence_and_early_farming_from_neolithic_settlements_in_Central_Europe_The_archaeozoological_evidence_from_the_Czech_Republic/links/09e41513f7194e14ac000000/The-new-insights-into-the-subsistence-and-early-farming-from-neolithic-settlements-in-Central-Europe-The-archaeozoological-evidence-from-the-Czech-Republic.pdf).

68. T. G. Papachristou, L. E. Dziba, F. D. Provenza, Foraging ecology of goats and sheep on wooded rangelands. Small Rumin. Res. 59, 141–156 (2005).

69. P. Hejcmanová, M. Stejskalová, M. Hejcman, Forage quality of leaf-fodder from the main broad-leaved woody species and its possible consequences for the Holocene development of forest vegetation in Central Europe. Vegetation history and archaeobotany. 23, 607–613 (2014).

70. M. Hejcman, P. Hejcmanová, M. Stejskalová, V. Pavlů, Nutritive value of winter-collected annual twigs of main European woody species, mistletoe and ivy and its possible consequences for winter foddering of livestock in prehistory. The Holocene. 24, 659–667 (2014).

71. M. S. DeFelice, Henbit and the deadnettles, Lamium spp.—archangels or demons? Weed Technol. 19, 768–774 (2005).

72. G. Larson, U. Albarella, K. Dobney, P. Rowley-Conwy, J. Schibler, A. Tresset, J.-D. Vigne, C. J. Edwards, A. Schlumbaum, A. Dinu, A. Balaçsescu, G. Dolman, A. Tagliacozzo, N. Manaseryan, P. Miracle, L. Van Wijngaarden-Bakker, M. Masseti, D. G. Bradley, A. Cooper, Ancient DNA, pig domestication, and the spread of the Neolithic into Europe. Proceedings of the National Academy of Sciences. 104, 15276–15281 (2007).

73. M. N. Barrios-Garcia, S. A. Ballari, Impact of wild boar (Sus scrofa) in its introduced and native range: a review. Biol. Invasions. 14, 2283–2300 (2012).

74. A. J. Carpio, M. García, L. Hillström, M. Lönn, J. Carvalho, P. Acevedo, C. G. Bueno, Wild Boar Effects on Fungal Abundance and Guilds from Sporocarp Sampling in a Boreal Forest Ecosystem. Animals (Basel). 12 (2022), doi:10.3390/ani12192521.

75. P. Pokorný, P. Kuneš, Holocene acidifi cation process recorded in three pollen profi les from Czech sandstone and river terrace environments, (available at https://epic.awi.de/36253/10/Ferrantia_44-107.pdf).

76. E. Ellis, M. Maslin, N. Boivin, A. Bauer, Involve social scientists in defining the Anthropocene. Nature Publishing Group UK (2016), doi:10.1038/540192a.

77. Y. Wang, M. W. Pedersen, I. G. Alsos, B. De Sanctis, F. Racimo, A. Prohaska, E. Coissac, H. L. Owens, M. K. F. Merkel, A. Fernandez-Guerra, A. Rouillard, Y. Lammers, A. Alberti, F. Denoeud, D. Money, A. H. Ruter, H. McColl, N. K. Larsen, A. A. Cherezova, M. E. Edwards, G. B. Fedorov, J. Haile, L. Orlando, L. Vinner, T. S. Korneliussen, D. W. Beilman, A. A. Bjørk, J. Cao, C. Dockter, J. Esdale, G. Gusarova, K. K. Kjeldsen, J. Mangerud, J. T. Rasic, B. Skadhauge, J. I. Svendsen, A. Tikhonov, P. Wincker, Y. Xing, Y. Zhang, D. G. Froese, C. Rahbek, D. N. Bravo, P. B. Holden, N. R. Edwards, R. Durbin, D. J. Meltzer, K. H. Kjær, P. Möller, E. Willerslev, Late Quaternary dynamics of Arctic biota from ancient environmental genomics. Nature. 600, 86–92 (2021).

78. M. Meyer, M. Kircher, Illumina sequencing library preparation for highly multiplexed target capture and sequencing. Cold Spring Harb. Protoc. 2010, db.prot5448 (2010).

79. M. Schubert, S. Lindgreen, L. Orlando, AdapterRemoval v2: rapid adapter trimming, identification, and read merging. BMC research notes. 9, 1–7 (2016).

80. W. T. T. Taylor, M. Pruvost, C. Posth, W. Rendu, M. T. Krajcarz, A. Abdykanova, G. Brancaleoni, R. Spengler, T. Hermes, S. Schiavinato, G. Hodgins, R. Stahl, J. Min, S. Alisher Kyzy, S. Fedorowicz, L. Orlando, K. Douka, A. Krivoshapkin, C. Jeong, C. Warinner, S. Shnaider, Evidence for early dispersal of domestic sheep into Central Asia. Nat Hum Behav. 5, 1169–1179 (2021).

81. H. Mannen, T. Yonezawa, K. Murata, A. Noda, F. Kawaguchi, S. Sasazaki, A. Olivieri, A. Achilli, A. Torroni, Cattle mitogenome variation reveals a post-glacial expansion of haplogroup P and an early incorporation into northeast Asian domestic herds. Sci. Rep. 10, 20842 (2020).

82. D. H. Parks, M. Chuvochina, C. Rinke, A. J. Mussig, P.-A. Chaumeil, P. Hugenholtz, GTDB: an ongoing census of bacterial and archaeal diversity through a phylogenetically consistent, rank normalized and complete genome-based taxonomy. Nucleic acids research. 50, D785–D794 (2022).

