## Supplementary Information for "Early Pastoralism in Central European Forests: Insights from Ancient Environmental Genomics"

**This PDF file includes:**

Supporting text
Table S1
Figures S1 to S15
Legends for Datasets S1 to S7
SI References

**Other supporting materials for this manuscript include the following:**

Datasets S1 to S7 (pdf format)

### SI1 Chronometric Data and Age Modeling of DNA Samples

All dates were calibrated in Oxcal v 4.4.4(1) using the IntCal20(2). Samples were taken during the initial excavation of the test trench at Velký Mamuťák in sectors B and D; exceptions include Beta-473738, Poz-97105, Poz-97106, and taken in the microstratigraphy column in sector B profile (Table S1). The age-depth model (Figure S1) was constructed assuming a non-linear deposition (mediated by a Poisson process). The model was created using Oxcal v 4.4.4(1) with r:5 atmospheric data (IntCal20) from Reimer et al (2020)(2). The age-depth model indicates a relatively steady sedimentation rate and fluctuations have been noted to align with general changes in precipitation during the Holocene(3).

### SI2 Library quality assessment

In this paragraph, we assess the quality of our libraries by investigating the cycles of index PCR used for amplification and the proportion of duplicates detected by filtering the sequences with SGA (v. 0.10.15). As a comparison to the remainder of the dataset, we also display the fraction of reads that map to the superkingdom of Eukaryotes, which includes both animals and plants (Figure S2).

We find the level of duplication for all samples to vary between 23%–95%, (Fig. S2b), in particular samples from the top and bottom layers, had high levels of duplicates (75 and 95% VM 2-11 and VM 22-28 respectively, Dataset S1). The high duplication rate seems to correlate with a relatively higher number of cycles needed to amplify the libraries sufficiently as found by quantitative PCR (Fig. S2a). For the reasons stated in the Discussion(4–8), this is commonly caused by a low abundance of DNA molecule templates within the extracts. The libraries with cycles between 13 and 15 (VM 14–19; Fig. S2c–d) had low levels of duplication <30% and showed increased complexity. We also evaluated our sequencing effort (x-axis) and the variety of distinct taxa categorized at the genus level (y-axis) for each library using the function "rarefy" (R package vegan v. 2.5-7). This demonstrates how, after accounting for variations in genus richness (Fig. S2e), the lowest-ranking samples (VM22-28) show a lower diversity (50-100 taxa) when compared to the other samples.

### SI3 MetaDMG for animals and plants

#### Setting the threshold for filtering ancient authentic reads

We initially ran metaDMG in LCA mode using likelihood estimation of damage patterns for all LCA ranks: --simscorelow 0.95, --simscorehigh 1.0, --maxposition 15, --weight-type 1, --lca-rank " ", --bayesian: false. We then visualized the damage profiles and plots with the metaDMG interactive dashboard, setting the x and y axes to likelihood ratio ($\lambda$LR) and degree of damage ($D$fit) respectively. We also filtered the data by read number ($\geq 20$) and taxa rank at the genus level.

Using the range of the negative controls as a reference, a threshold considering the level of damage ($D\text{fit} \geq 0.05$) and uncertainty ($\lambda\text{LR} \geq 10$) was set with the purpose of targeting only authentic ancient reads.

Given the high abundance of taxa with a low number of reads (ranging from 20 to 150), a high standard deviation ($\geq$10), and a phi value ($\geq$100), metaDMG was run again using a fully Bayesian model (--bayesian: true), aiming at a better resolution for the fitting model of those taxa (Fig. S3 a-c). We then set the x and y axes to the significance for the fitting model ($Z_{fit}$) and level of damage ($D_{fit}$) respectively, keeping only reads with a degree of damage greater than 5%. Moreover, we considered additional parameters: the concentration for the beta-binomial distribution (Bayesian_phi) and the standard deviation of $D$fit (Bayesian_D_max_std).

The final settings we considered to be optimal for our dataset are the following: N_reads ≥ 100, $D$fi ≥ 0.05, Zfit ≥ 2, Bayesian_phi ≥ 100, Bayesian_D_max_std ≤ 0.10. Besides, to get a more accurate resolution of our filtered taxa, we selected only the reads classified at the genus level.

Based on the range of controls and the damage fit result (Fig. S4 a-b), the value of $D$fit at 0.05 and a significance ($Z_{fit}$) greater than 2 (the latter also recommended by Michelsen et al. 2022)(9) have been confirmed as a reliable threshold for our dataset, allowing us to eliminate false positive and modern contamination (Fig. S4). For example, in the discarded reads for animals (Fig. S4a), human DNA (*Homo*) is found in samples VM-17, VM-22, VM-28, and VM-26, and pig DNA (*Sus*) is found in sample VM-3. Because VM-3 is one of the most recent layers, it is reasonable to detect such a low level of damage (2%), but for Mesolithic/Late Neolithic layers, Homo reads with a $D_{fit}$ of 0-0.01 are likely modern contamination (Dataset S2).

Fragment length distribution by depth (Fig. S8, S9) and mean length (Fig. S5) of the most abundant taxa found, independently confirm the authenticity of our parsed reads. The metaDMG output for the animal and plant reads can be found in Dataset S2.

**Damage profile through time**

Using the entire dataset and visualizing the degree of damage by depth, it was possible to confirm changes in ancient molecules' detection over time. The profile for animals and plants by depth (Fig. S7) shows an overall increase in the degree of damage over time, until layer 196. (VM-24). The samples at the bottom (VM-26, VM-28), which correspond to Mesolithic strata (222, 236), do not contain reads with a degree of damage greater than 10%, and only a few taxa were found (see section 2 about complexity). While no reads above 5% of damage were detected in one sample (VM22) at a depth of 185 cm (Fig. S7a). The low abundance of molecules detected in these samples is most likely the reason for the lack of significant damage.

After applying our threshold (above paragraph), the profile for the same reads includes only layers 64,107,116,134,151,196 (VM-11, VM-14, VM-15, VM-17, VM-19, VM-24) and reports a level of damage ranging from 5 to 16% for animals and 5 to 14% for plants (Fig. S7b). The $D$fit, and mean length (with standard deviation) of the most abundant animals and plants detected also support this trend (Figures S8, S9).

**SI4 Phylogenetic placement**

We sought to phylogenetically place the mammalian mitochondrial DNA by following the method described in Kjær et al. (2022)(10, 11) that investigates biallelic transversions and transitions of the aligned reads to the reference mitochondrial genomes (see Methods

section). The summary statistics of the aligned reads to the consensus sequence for *Ovis* and *Bos* for PATHPHYNDER analysis can be found in Figs. S 11-12.
We then ran metaDMG with the "global" damage-mode setting to get an estimation of the damage for all the mapped reads for each sample. For both *Ovis* and *Bos* reads, we estimated a level of damage between 6 and 17% (Dataset S5).
Regarding the reads aligned to *Ovis* mitogenomes, we report a mean read depth of 0.363 for sample VM-17 covering 25.3 percent of the mitochondrial genome (4248 bp in total) with 110 reads. The support for the placement in *Ovis aries* haplogroup B has a read depth of nine for a total of 14 SNPs analyzed.
On the other hand, sample VM-19 counts for a lower number of reads mapping (14) and a mean read depth of 0.038 covering 3.56 percent of the consensus mitogenome (598 bp in total). The support for the placement in *Ovis aries* haplogroup B/*Ovis aries musimon* has a read-depth of two for a total of 14 SNPs analyzed.
We report a mean read depth of 0.464 for sample VM-17 when mapping the reads to the consensus sequence of Bos mitogenomes for PATHPHYNDER analysis, covering 34.9 percent of the mitochondrial genome (5,707 bp in total) with 172 reads. With 113 reads, we recover a mean read depth of 0.364 covering 29.4 percent of the consensus mitogenome (4803 bp in total). The mapping statistics for VM-14 and VM-19 are very similar: the mean read depth is 0.147 and 0.145, respectively, covering 13.3 and 13.5 percent of the mitogenome (2277 bp and 2177 bp) with 58 and 50 reads.
The average read length for all the samples ranges between 41 and 55 bp and all reads have a mean mapping quality above 30 (Dataset S5).
PATHPHYNDER algorithm failed the analysis of transition and transversion for sample VM-19, for this reason, we excluded this sample for further haplotype identification.
With the purpose of further investigating the quality of the placement produced by PATHPHYNDER, we manually visualized the transitions and their position in the mapped sequences by using SEAVIEW software (v. 5.0.5)(12) with the purpose of double-checking the presence of deamination at the beginning of the reads for informative markers. We only identified one deaminated base at the beginning of the read for A> G transition in sample VM-17 in a SNP supporting *Bos* reads placement basal to haplogroup P, Q, and T (Dataset S5). This does not change our previous interpretation, since only one read was excluded from the supporting SNPs.

**SI5 Sourcetracker2 analyses**
**MetaDMG for the microbial community**

The dataset was filtered for DNA damage ($D_{fit}$) above 5% and significance ($Z_{fit}$) ≥ 2 (see Supplementary section 3). As seen for the animal and plant datasets, the bottom-most layers (185-236 cm) present a lower degree of damage (1-2%) and a wider range of mean length

(Fig. S14). These are consistent with the low complexity found for the same layers in the animal and plant datasets (Fig. 2).

### Sourcetracker2 with applied metaDMG

A total of 7 gut metagenomes and 15 environments were run through Sourcetracker2 (Dataset S6). The output was grouped by source categories (Dataset S7). Multiple sources were utilized to cover diverse environments, and then categorized under "Other soils". These included metagenomes from the desert, freshwater, permafrost, melting permafrost, agricultural soil, tropical forests, and biocrust.
Overall, the taxonomic profile with reads parsed for DNA damage replicates the taxonomic profile shown in Fig. 3, but with fewer taxa detected (Fig. S15; Dataset S7).

**Table S1. Chronometric Data of DNA Samples.** All dates were calibrated in Oxcal v 4.4.4(1) using the IntCal20(2). Strata are identified by mechanical excavation layers and the nearest associated context number. Bolded laboratory codes are new dates from Velký Mamuťák published with this study.

| **Laboratory Code** | **Depth (-m)** | **Sector** | **Excavation Layer** | **Material** | **Taxon** | **Date (14C years BP)** | **cal BP 95.4% confidence level** |
|---|---|---|---|---|---|---|---|
| Poz-104297 | 0.36 | B | 30-35 (26) | seed | *Centaurea cyanus* | 870±30 | 903-689 |
| Beta - 473738 | 0.39 | B | F30-40/2 | needle | *Abies alba* | 2200±30 | 2317-2122 |
| Poz-104298 | 0.41 | B | 38,5-44 (30) | twig | *Viscum album* | 2200±30 | 2317-2122 |
| Poz-99530 | 0.49 | D | 40-45 (55) | animal dropping | | 2575±30 | 2759-2519 |
| Poz-104374 | 0.66 | B | 55-60 (63) | panicle (5 spikes) | *Panicum* | 3010±30 | 3335-3076 |
| **UGAMS - 59900** | 0.89 | B | 75-80 (72) | nutshell (charred) | *Corylus avellana* | 3510±30 | 3871-3695 |
| Poz-97105 | 1.16 | B | F100-110/2 | charcoal | | 3750±35 | 4235-3985 |
| Poz-104309 | 1.19 | B | 105-110 (91) | animal dropping | | 4150±35 | 4828-4535 |
| Poz-97106 | 1.64 | B | F150-160/1 | charcoal | | 5930±40 | 6881-6663 |
| Poz-104296 | 1.75 | B | 160-165 (125) | animal dropping | | 5900±40 | 6844-6635 |

| | | | | | | | |
|---|---|---|---|---|---|---|---|
| **UGAMS - 59903** | 1.98 | B | 185-190 (132) | twig (charred) | | 6130±30 | 7159-6907 |
| Poz-104310 | 2.18 | B | 205-210 (134a) | nutshell (charred) | *Corylus avellana* | 8540±50 | 9597-9445 |
| **UGAMS - 59904** | 2.39 | B | 225-230 (134b) | nutshell (charred) | *Corylus avellana* | 8810±30 | 10119-9689 |
| Beta-473739 | 2.47 | B | 235-240 (147) | needle) | *Pinus sylvestris* | 9450±30 | 10987-10578 |

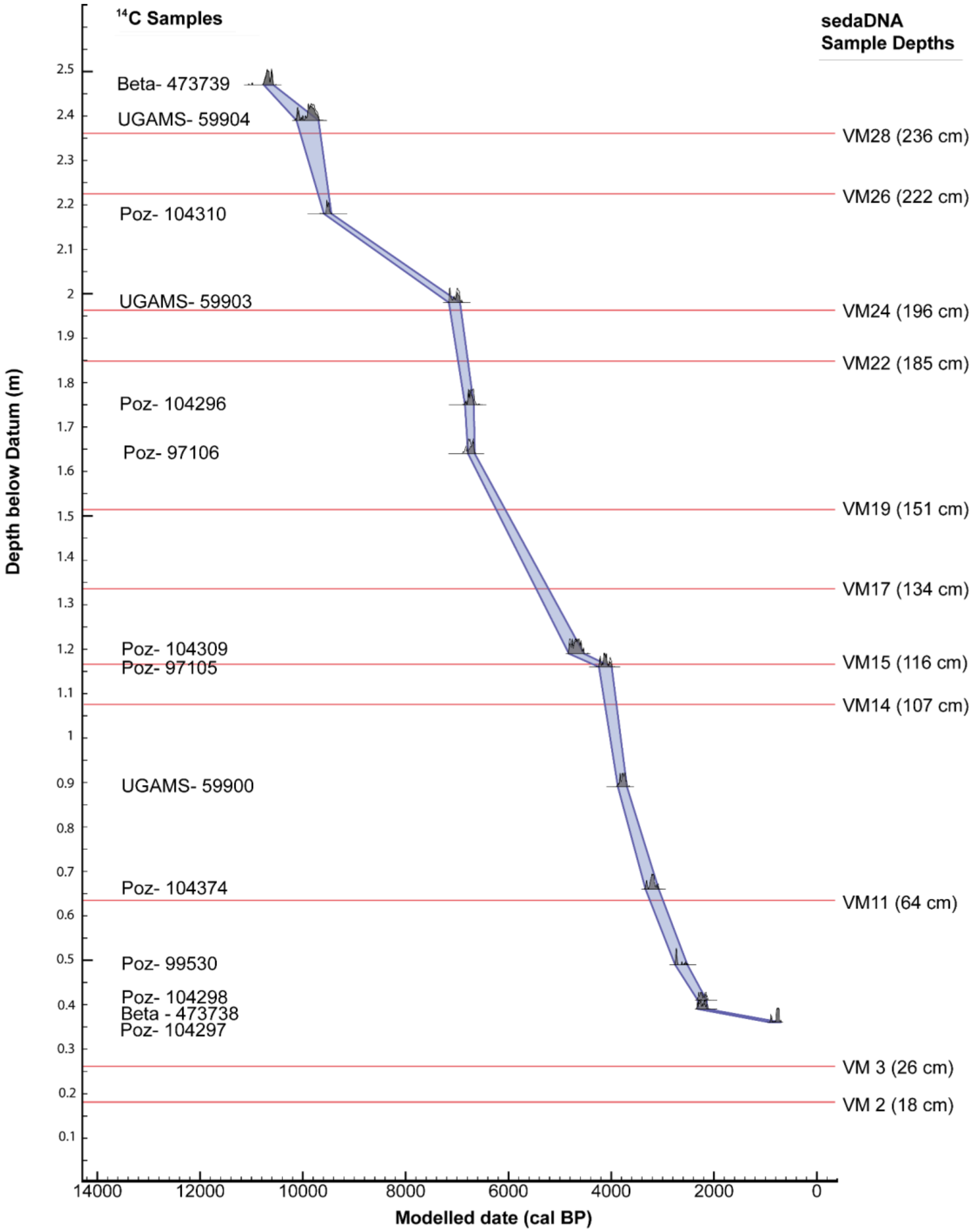


**Fig. S1. Age depth model for Velký Mamuťák.** The model was created using Oxcal v 4.4.4(1) with r:5 atmospheric data (IntCal20) from Reimer et al (2020)(2). The blue band indicates the 95.4% confidence interval and the depths of each *seda*DNA sample are plotted along the right column.

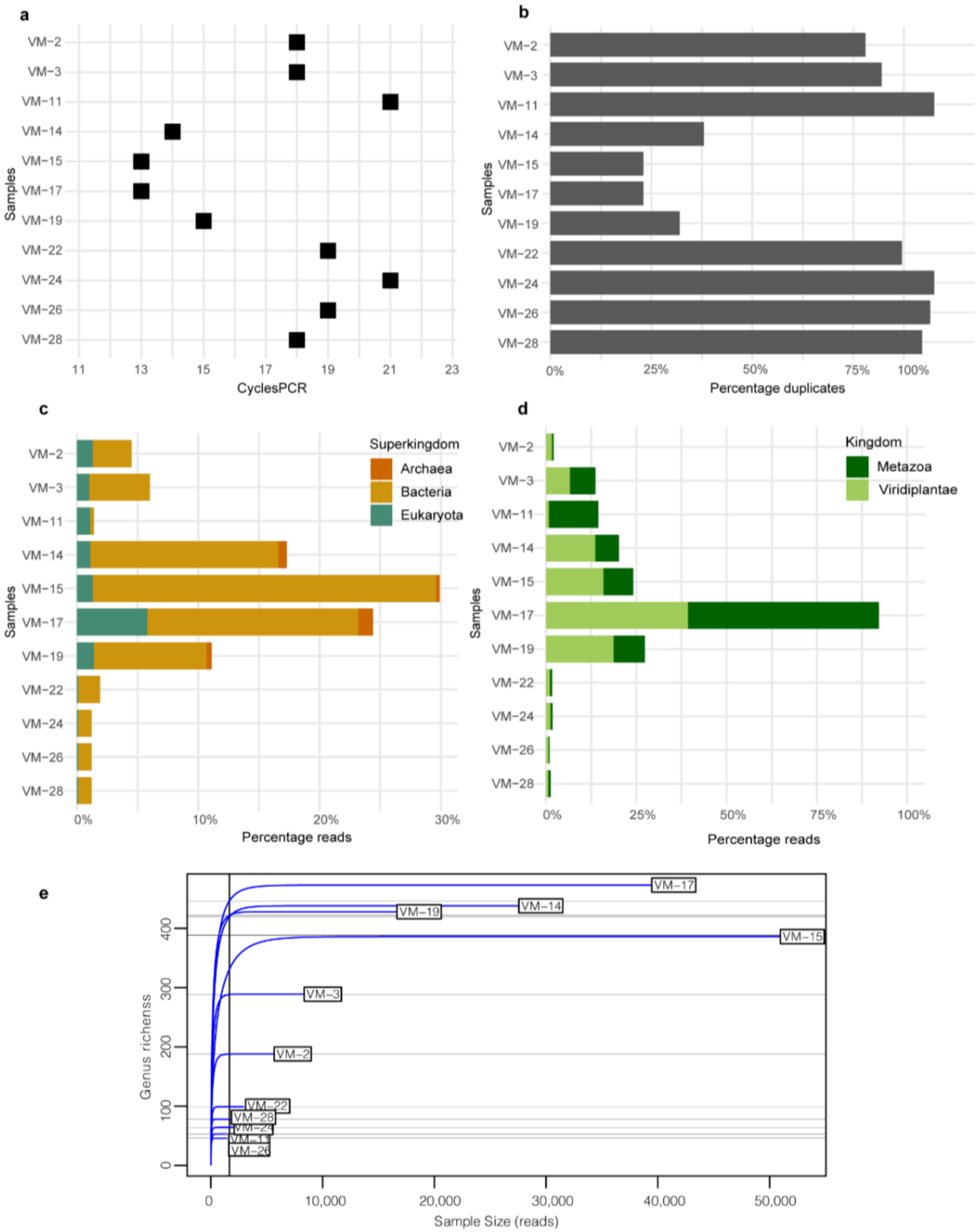


**Fig. S2. Library quality assessment. a.** PCR amplification cycles used per each sample based on qPCR ct-values -2. **b.** Percentage of duplicates for each library after filtering for low complexity reads. **c**. The proportion of Archaea, Bacteria, and Eukaryotes in the aligned reads after duplicate removal against the total number of reads. **d.** The proportion of animal and plant DNA in the aligned reads after duplicate removal against the total number of reads**. e.** Rarefaction curve showing the number of different genus frequencies (y-axis) and the sequencing depth calculated in the number of reads (x-axis).

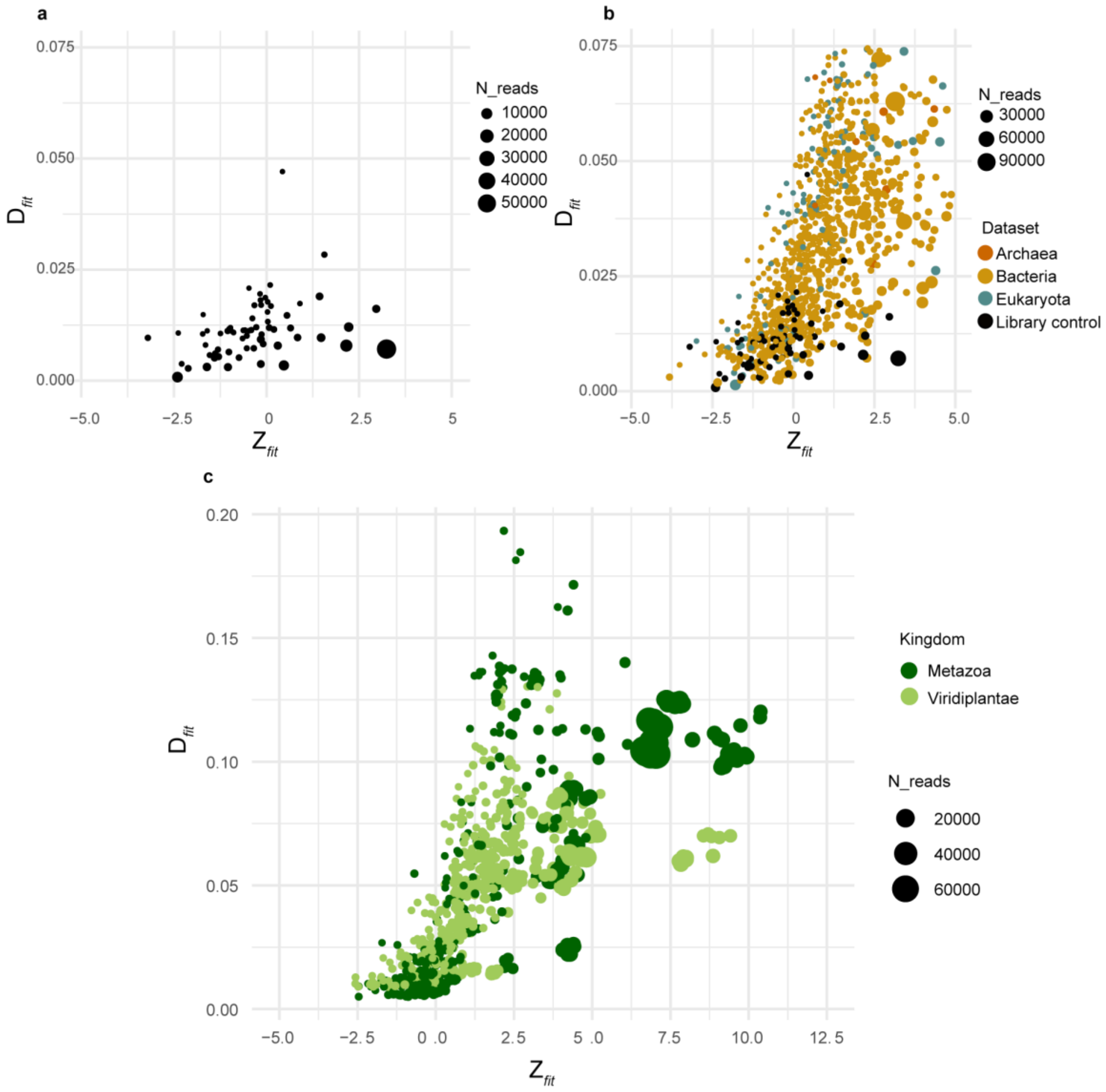


**Fig. S3. Level of damage for the whole dataset with the Bayesian fitting model.**
**a.** Degree of damage (*D*fit) and significance (Zfit) for the library controls. Number of reads ≥ 100. **b.** Level of damage (*D*fit) and significance (Zfit) for the aligned reads divided into superkingdoms (Archaea, Bacteria, Eukaryota) together with the library controls (black). **c.** Degree of damage (*D*fit) and significance (Zfit) for animals and plants (reads ≥ 100).

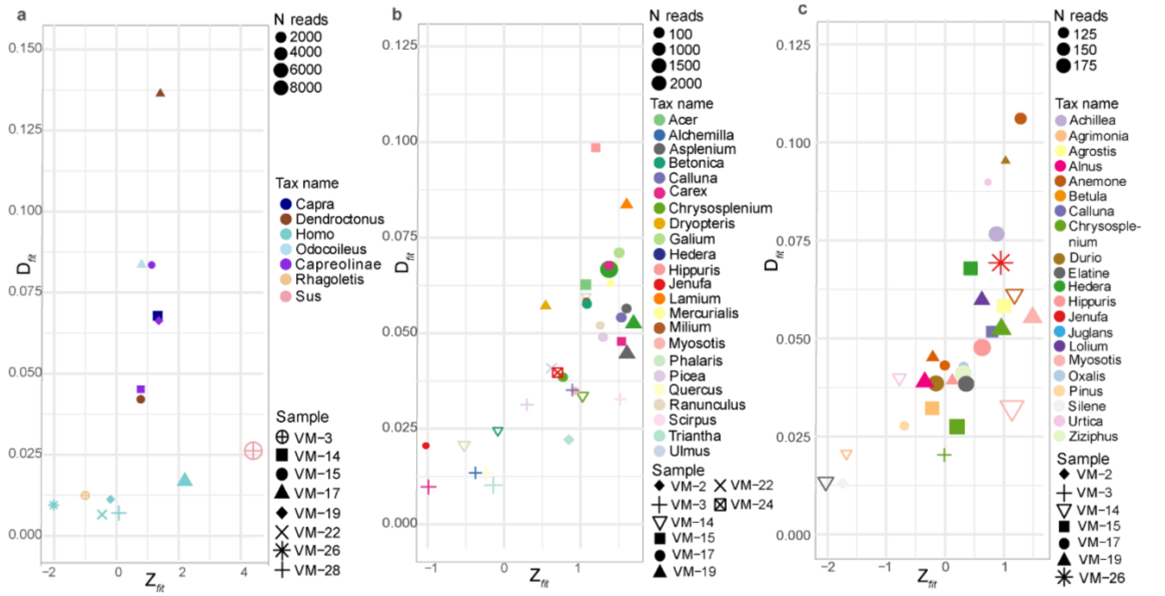


**Fig. S4. Discarded reads for animals and plants using the Bayesian fitting model at the genus level.**

**a.** Discarded reads for animals with significance ($Z_{fit}$) < 2 and a significance ($Z_{fit}$) > 2 but with a degree of damage (*D*fit) < 0.05. Number of reads > 100. **b-c.** Discarded reads for plants with significance ($Z_{fit}$) <2. Number of reads > 200 (b), 100>200 (c).

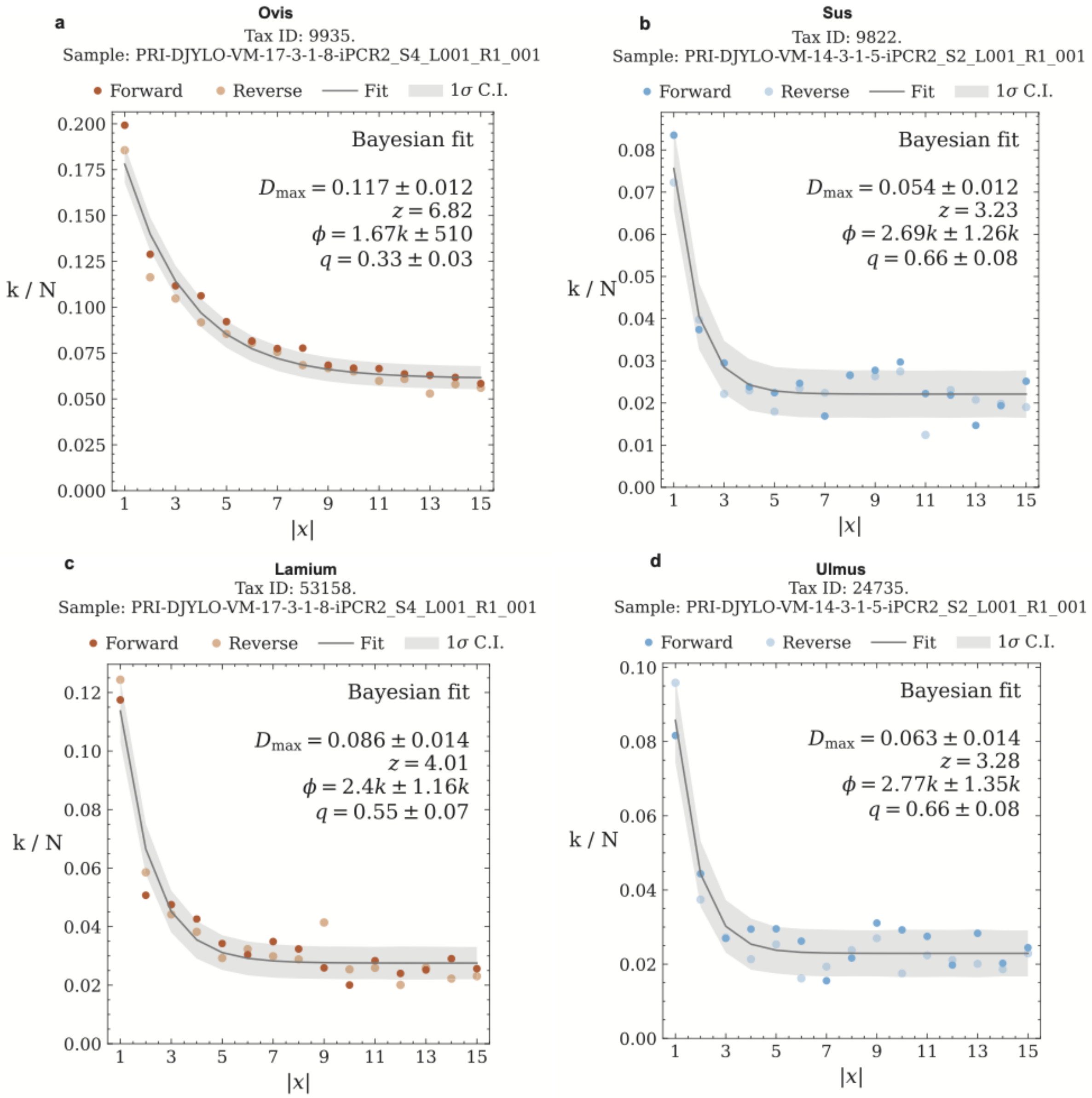


**Fig. S5. Position (x) of specific nucleotide mis-incorporations (k/N) due to DNA damage on the forward and reverse strand for most abundant animals and plants found (Bayesian_z ≥ 2).**
Dark-colored dots are C > T transitions, light-colored dots are G > A transitions, gray areas indicate the Bayesian fit. a. Sheep (*Ovis*), sample VM-17, *D*fit: 0.117 ± 0.012, Zfit: 6.82, nr reads: 53.7k, meanL: 55.4 ± 17.1. b. Pig (*Sus*), sample VM-14, *D*fit: 0.054 ± 0.012, Zfit: 3.23, nr reads: 4.32k, meanL: 42.1 ± 11.7. c. Dead-nettle (*Lamium*), sample VM-17, *D*fit: 0.086 ± 0.014, Zfit: 4.01, nr reads: 6.94k, meanL: 45.6 ± 14.0. d. Elm (*Ulmus*), sample VM-14, *D*fit: 0.063 ± 0.014, Zfit: 3.28, nr reads: 3.18k, meanL: 44.0 ± 11.0.

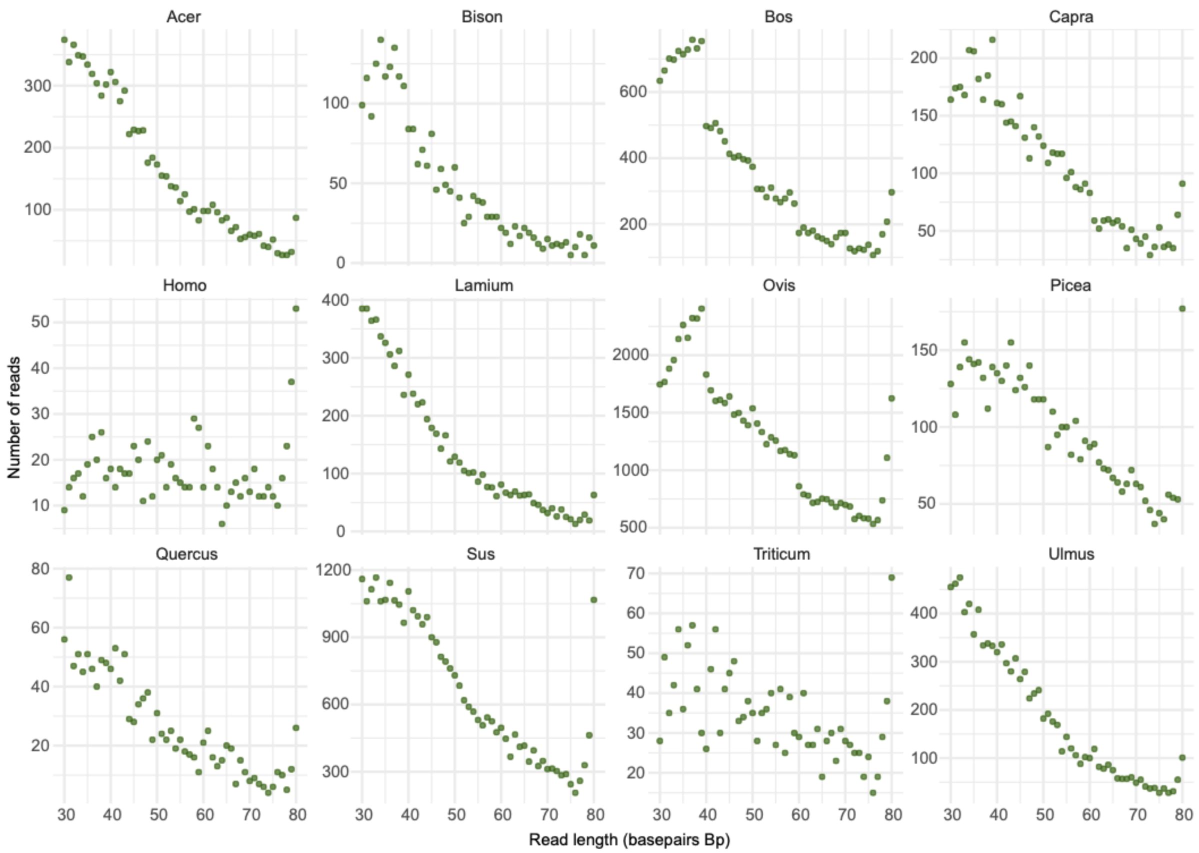


**Fig. S6. Read length distribution of all animal and plant reads across all the samples assigned to the same taxon.**

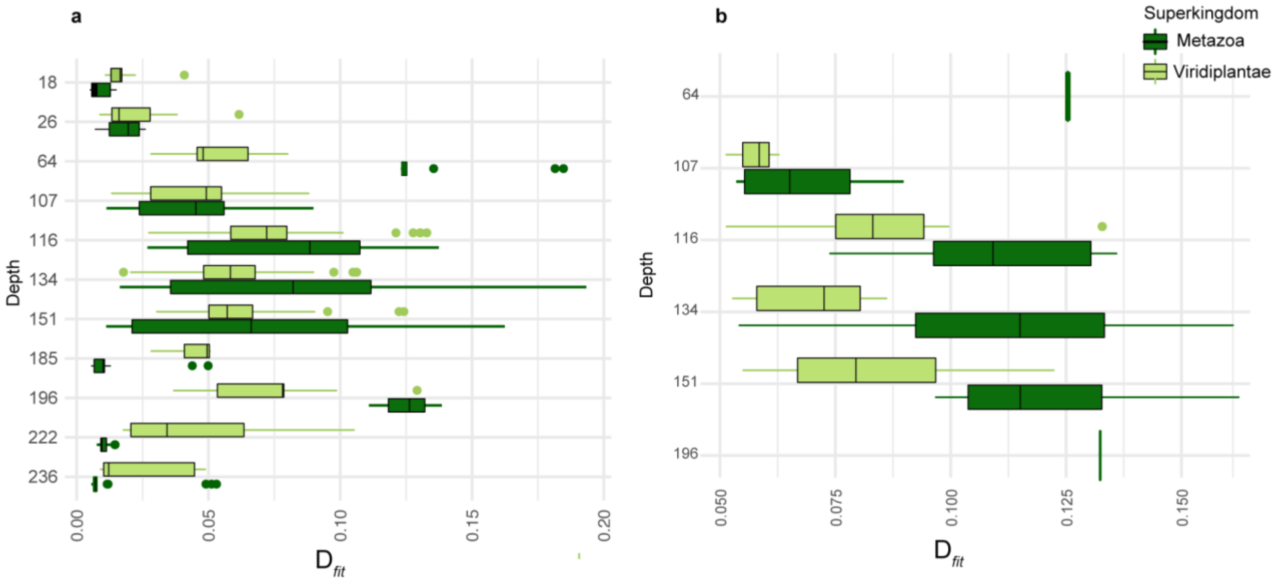


**Fig. S7. Damage profile by depth.**

Degree of damage (*D*fit) of plants and animals for each layer before (a) and after (b) applying the threshold for filtering ancient reads (Number of reads ≥ 100).

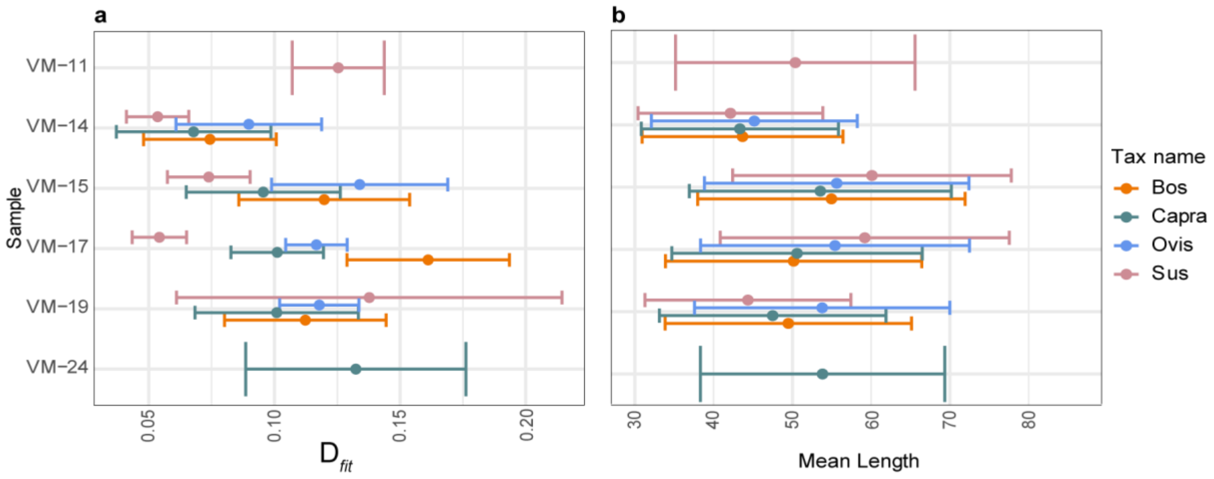


**Fig. S8. Degree of damage (a) and mean read length (b) for the most abundant animal taxa and their standard deviation.**

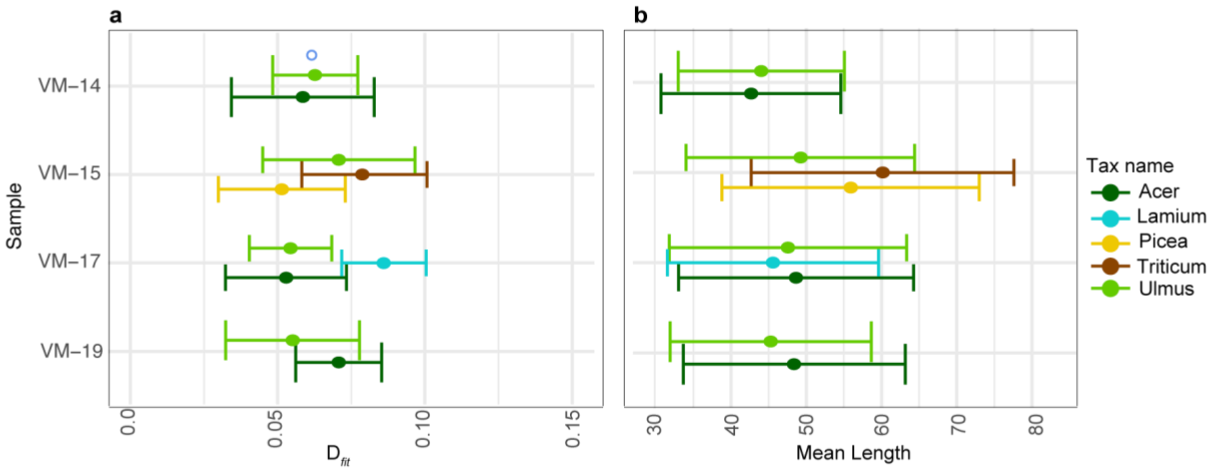


**Fig. S9. Degree of damage (a) and mean read length (b) for the most abundant plant taxa and their standard deviation.**

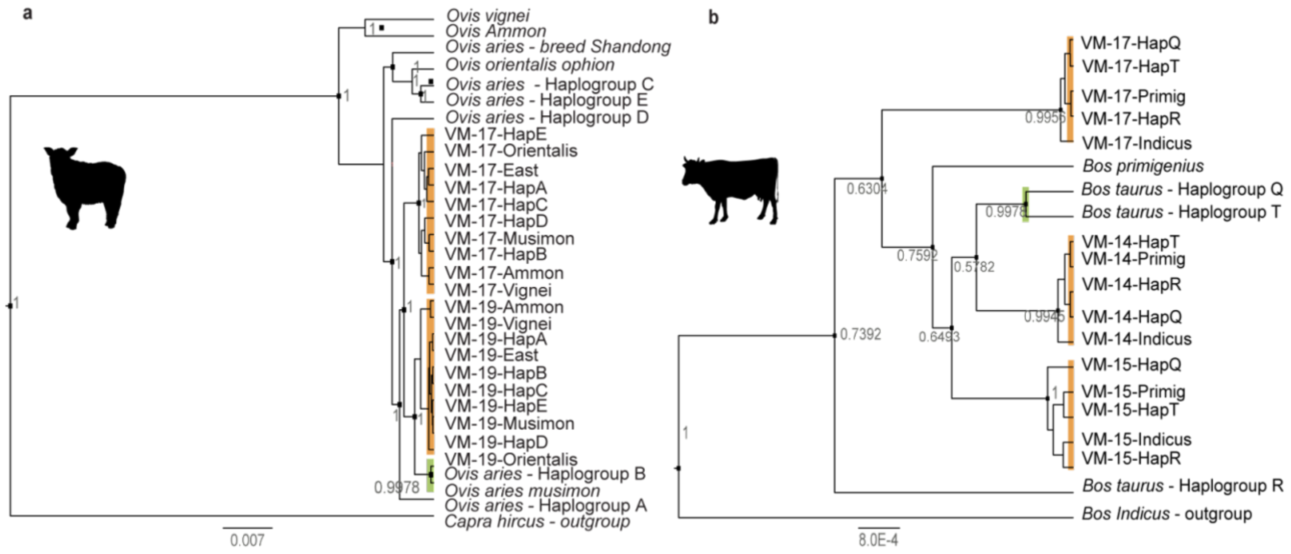


**Fig. S10. Phylogenetic placement of consensus mitochondrial genomes.** Phylogenetic relationships of reads assigned to *Ovis (*a*)* and *Bos* (c) inferred using Bayesian analysis. Colors of the branches differentiate the samples (orange) from the placement (green). **a**. Phylogenetic placement of *Ovis* reads for samples VM-17 and VM-19 (Late Neolithic) versus modern mitochondrial genomes of the domestic sheep (see Methods). **b**. Phylogenetic placement of *Bos* reads for samples VM-14, VM-15, VM-17 (Late Neolithic-Early Bronze Age) against modern mitochondrial genomes of the domestic cattle and the aurochs (see Methods).

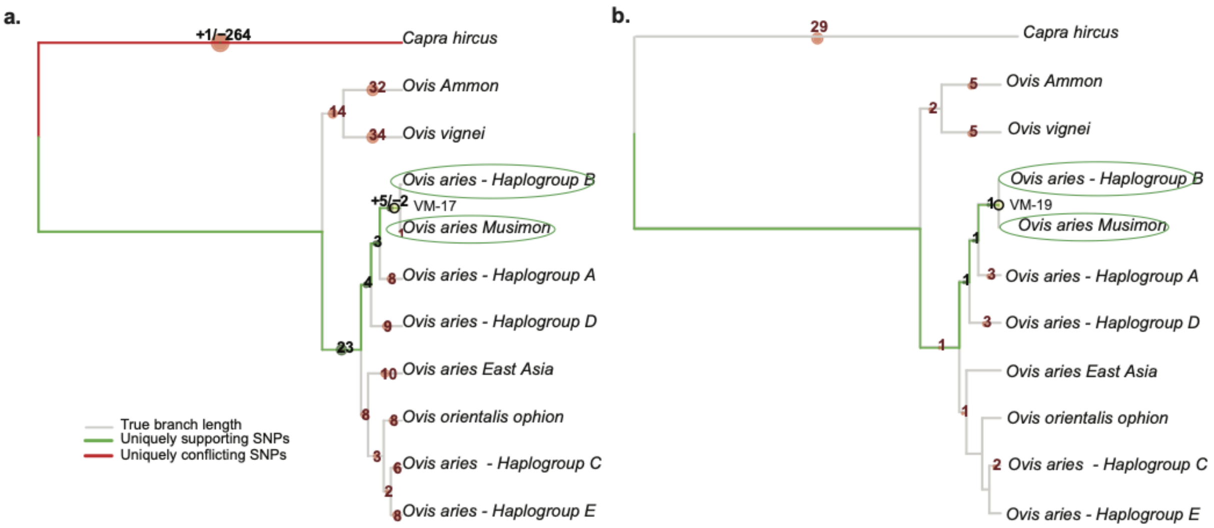


**Fig. S11. Pathphynder placement for *Ovis* reads.** The trees show the biallelic transition and transversion SNPs in VM-17 (a) and VM-19 (b) partitioned by read mapping. Green lines indicate shared supporting SNPs; red lines indicate uniquely conflicting SNPs. **a.** The aligned reads have a total of 23 supporting SNPs for the domesticated species *Ovis aries* with +5/-2 SNPs supporting the *Haplogroup B/Ovis aries musimon*. **b.** A single supporting SNPs leading to the branch *Ovis aries Haplogroup B/Ovis aries musimon*.

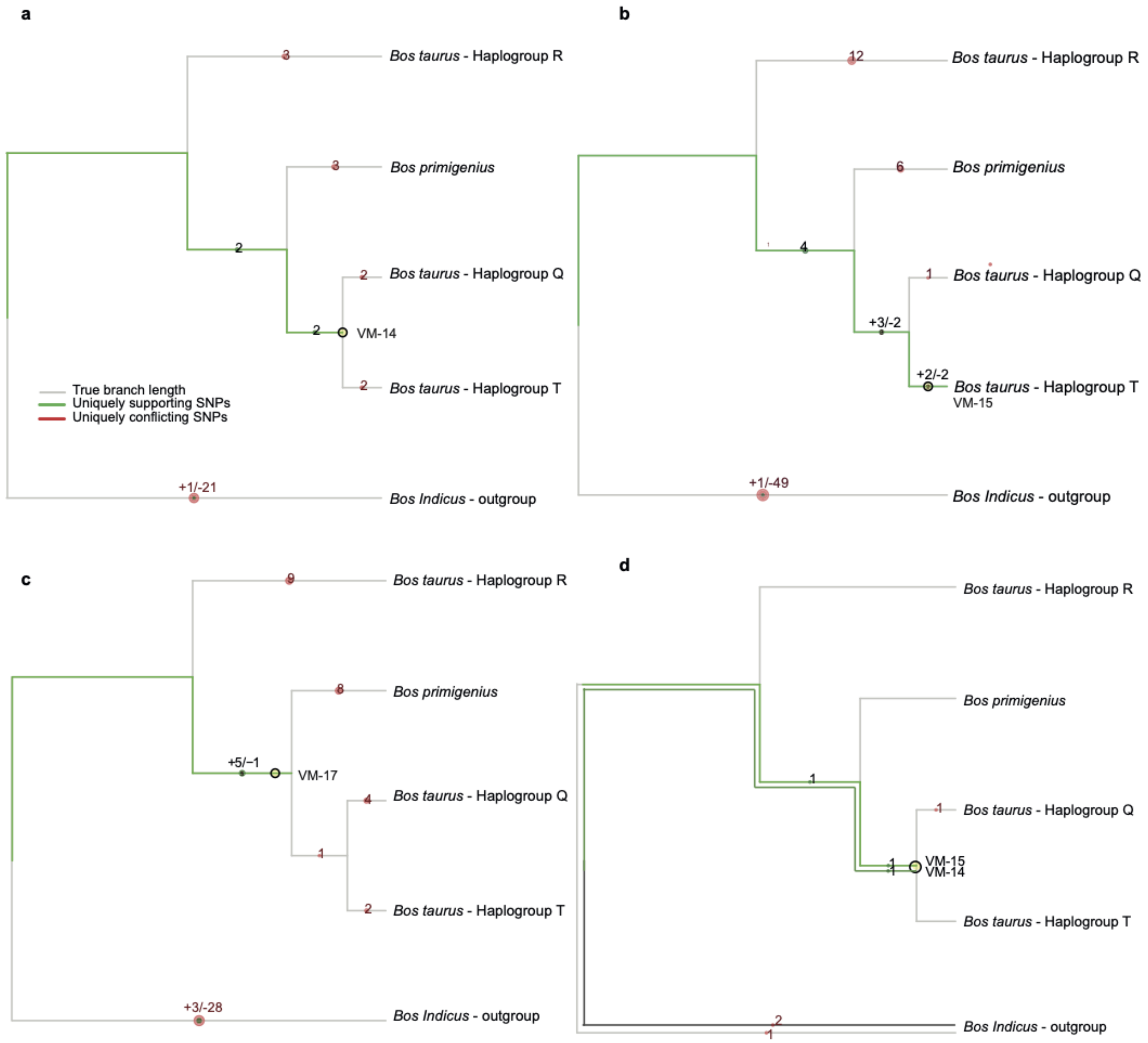


**Fig. S12 Pathphynder placement for *Bos* reads**.

Biallelic transition and transversion SNPs for samples VM-14 (a), VM-15 (b), VM-17 (c) and only transversions tree for samples VM-14 and VM-15 (d). Green lines indicate shared supporting SNPs; red lines represent uniquely conflicting SNPs. a. Aligned reads of sample VM-14 with 2 supporting SNPs dropping at the basal node of *Bos taurus* Haplogroup Q and T. b. Aligned reads of sample VM-15 with +3/-2 supporting SNP leading to the basal node of domesticated species of *Bos taurus* (haplogroup Q and T) and +2/-2 mismatches for *Bos taurus* haplogroup T. c. Aligned reads for sample VM-17 fall within the basal node of *Bos primigenius* and *Bos taurus* (haplogroup Q and T) with a support of +5/-1 SNPs. d. Transversions only tree for samples VM-14 and VM-15, both reporting 1 supporting SNPs to the basal node of *Bos taurus* haplogroup Q and T.

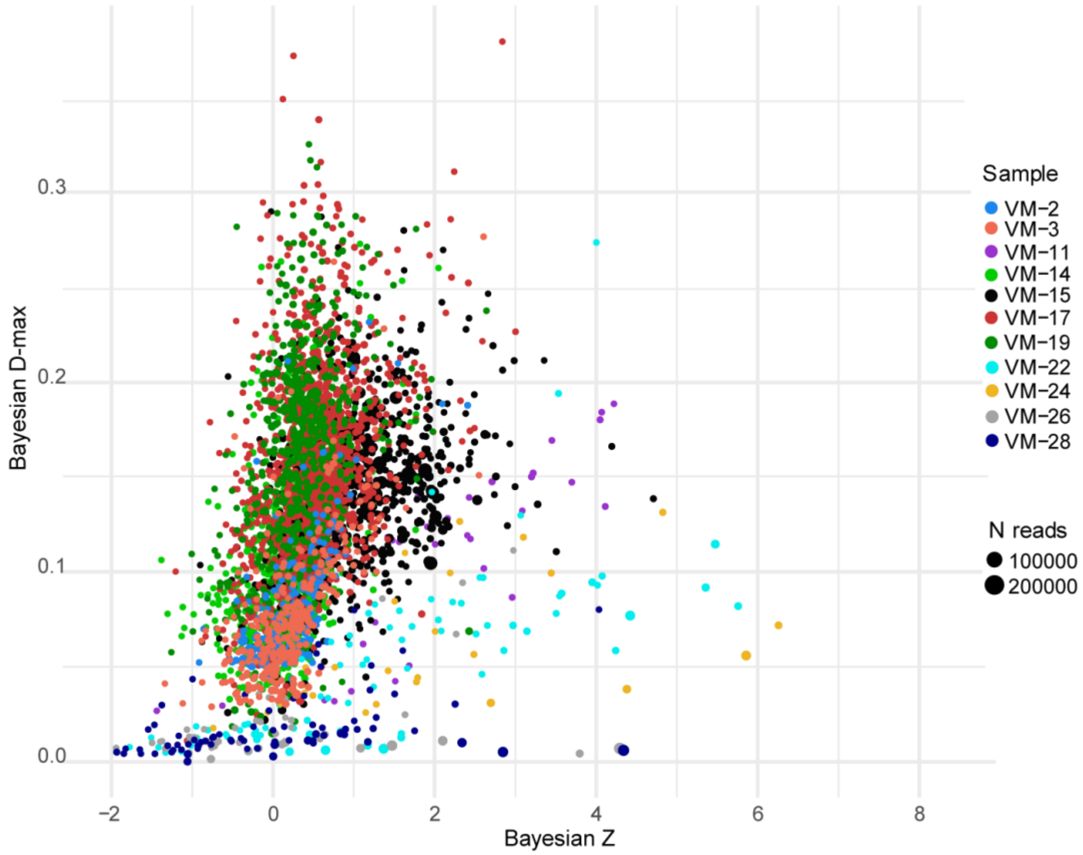


**Fig. S13. Degree of damage for the microbial dataset (at species level) with the Bayesian fitting model.**

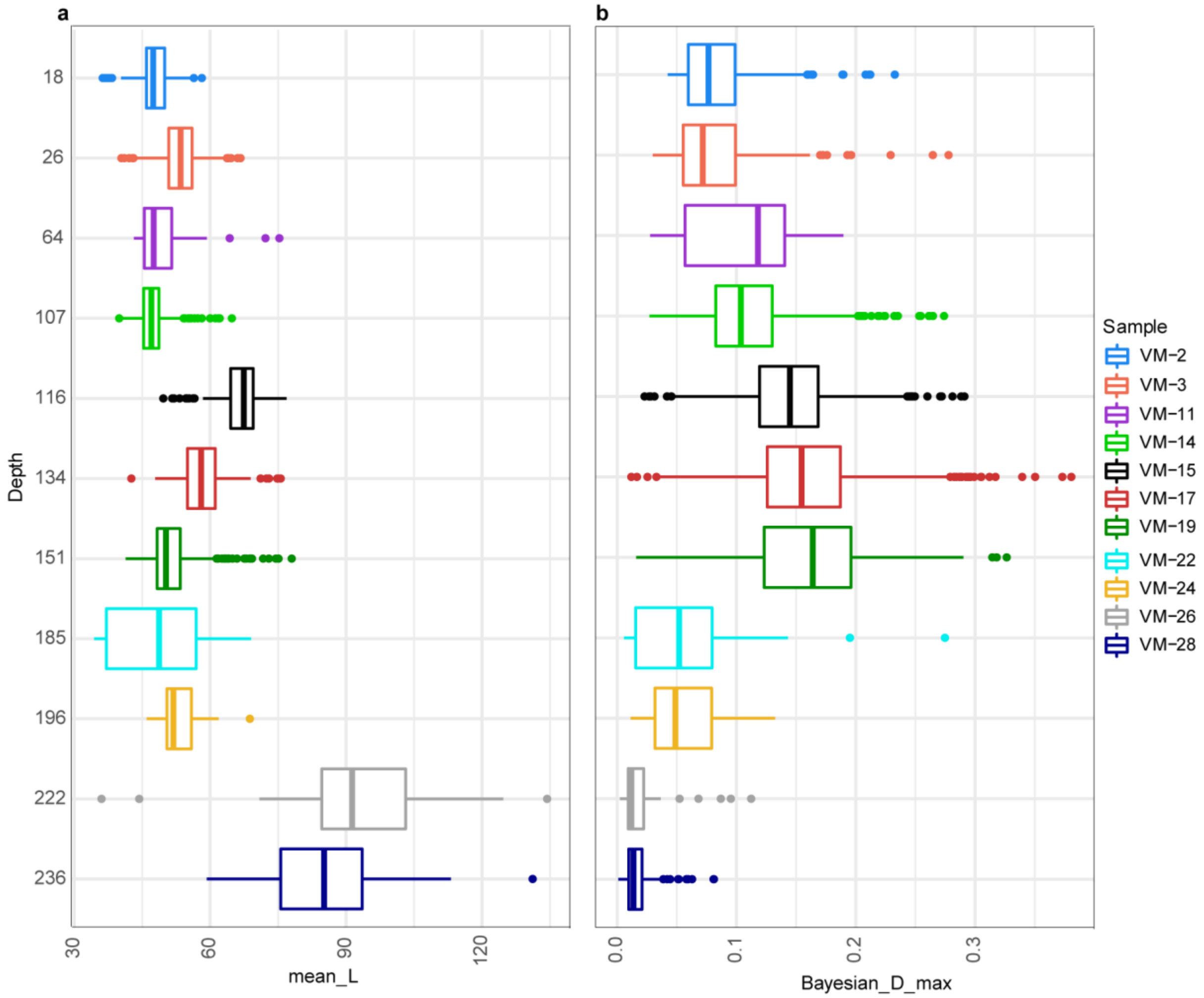


**Fig. S14. Mean read length (a) and degree of damage (b) for microbial DNA (at species level) by depth for the entire dataset.**

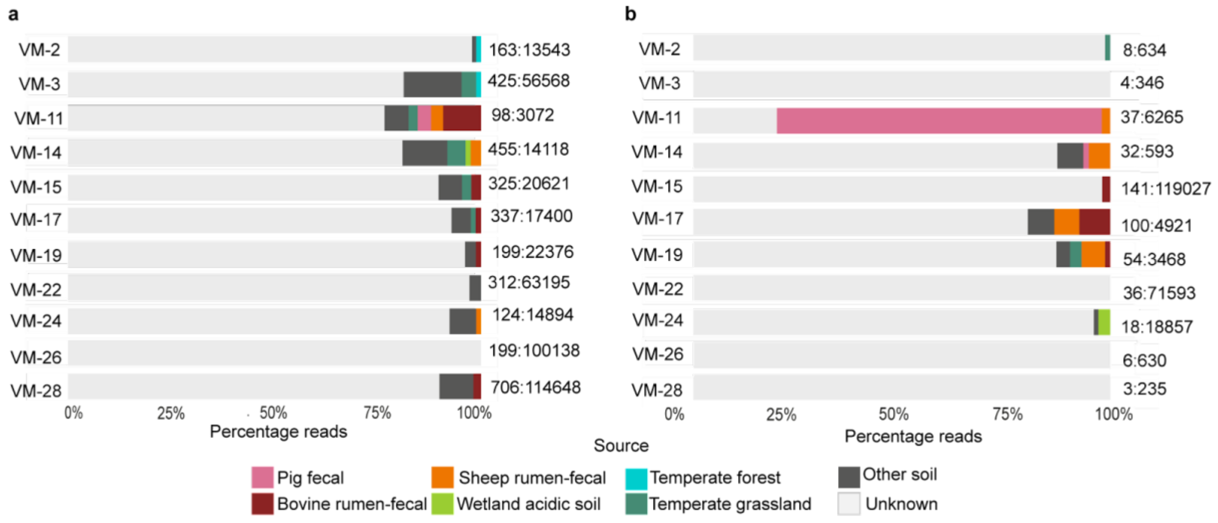


**Fig. S15. MetaDMG and Sourcetracker2 analyses for the microbial data.**
a-b. Microbial DNA parsed through Sourcetracker2 and metaDMG (at species level) with taxon count: reads count. $D_{fit} < 5$ (a); $D_{fit} \geq 5$, $Z_{fit} \geq 2$ (b).

**Dataset S1 (separate file). Overview of the laboratory procedures for sequenced (sheet 1) and not sequenced libraries (sheet 2).**

**Dataset S2 (separate file). MetaDMG output showing animal and plant reads included (sheets 1, 3) and discarded in the analyses (sheets 2, 4).**

**Dataset S3 (separate file). The taxa used for phylogenetic analyses (sheet 1) and the taxa excluded due to low mapping coverage (sheet 2).**

**Dataset S4 (separate file). List of references used for *Ovis* and *Bos* consensus genomes of the different haplogroups.**

**Dataset S5 (separate file). MetaDMG output for *Bos* and *Ovis* consensus genomes (sheet 1) and pathPhynder SNPs count (sheet 2).**

**Dataset S6 (separate file). List of metagenomic sources used for Sourcetracker2 analysis.**

**Dataset S7 (separate file). Sourcetracker2 outputs with the proportion for each reference source parsed through metaDMG: combined results (sheet 1), filtered dataset (sheet 2), reads below the thresholds (sheet 3).**

### SI References


1. C. B. Ramsey, Bayesian Analysis of Radiocarbon Dates. *Radiocarbon* **51**, 337–360 (2009).

2. P. J. Reimer, *et al.*, The IntCal20 Northern Hemisphere Radiocarbon Age Calibration Curve (0–55 cal kBP). *Radiocarbon* **62**, 725–757 (2020).

3. P. Šída, P. Pokorný, *Mezolit Severních Čech III: Vývoj Pravěké Krajiny Českého Ráje: Vegetace, Fauna, Lidé* (Archeologický ústav AV ČR, 2020).

4. G. Pietramellara, *et al.*, Extracellular DNA in soil and sediment: fate and ecological relevance. *Biol. Fertil. Soils* **45**, 219–235 (2009).

5. A. Wolińska, Z. Stępniewska, Dehydrogenase activity in the soil environment. *Dehydrogenases* **10**, 183–210 (2012).

6. T. Demeke, G. R. Jenkins, Influence of DNA extraction methods, PCR inhibitors and quantification methods on real-time PCR assay of biotechnology-derived traits. *Anal. Bioanal. Chem.* **396**, 1977–1990 (2010).

7. E. Wnuk, *et al.*, The effects of humic substances on DNA isolation from soils. *PeerJ* **8**, e9378 (2020).

8. S. Köchl, H. Niederstätter, W. Parson, DNA extraction and quantitation of forensic samples using the phenol-chloroform method and real-time PCR. *Methods Mol. Biol.* **297**, 13–30 (2005).

9. C. Michelsen, *et al.*, metaDMG – A Fast and Accurate Ancient DNA Damage Toolkit for Metagenomic Data. *bioRxiv*, 2022.12.06.519264 (2022).

10. K. H. Kjær, *et al.*, A 2-million-year-old ecosystem in Greenland uncovered by environmental DNA. *Nature* **612**, 283–291 (2022).

11. R. Martiniano, B. De Sanctis, P. Hallast, R. Durbin, Placing Ancient DNA Sequences into Reference Phylogenies. *Mol. Biol. Evol.* **39** (2022).

12. M. Gouy, E. Tannier, N. Comte, D. P. Parsons, Seaview Version 5: A Multiplatform Software for Multiple Sequence Alignment, Molecular Phylogenetic Analyses, and Tree Reconciliation. *Multiple Sequence Alignment: Methods and Protocols*, 241–260 (2021).
