## Supplementary material for "Early Pastoralism in Central European Forests: Insights from Ancient Environmental Genomics": Dataset S1

| SampleID | Volume (g) | Zymo Clean-up | MinElute Clean-up | Qubit | Volume DNA (ul) | qPCR-Ct | Index PCR Cycles | Qubit | Index Used | Bioanalyser Molarity(pm/l) | Library Molarity | Raw reads | Filtered reads | Duplicates reads | Duplicates (%) |
|---|---|---|---|---|---|---|---|---|---|---|---|---|---|---|---|
| **VM-2** | 0.25 | 1 | 0 | 2.86 | 20 | 20.03 | 18 | 11.8 | UDI0030 | 12687.2113 | 74.9 | 41389695 | 25970277 | 20146409 | 78 |
| **VM-3** | 0.25 | 1 | 0 | 3.54 | 20 | 20.19 | 18 | 2.06 | UDI0031 | 5553.5914 | 5.6 | 67498965 | 54813814 | 44681144 | 82 |
| **VM-11** | 0.25 | 1 | 0 | 13.1 | 20 | 23.25 | 21 | 5.2 | UDI0033 | 9397.5758 | 24.4 | 34994307 | 25208392 | 24059177 | 95 |
| **VM-14** | 0.25 | 1 | 0 | Too Low | 20 | 16.25 | 14 | 1.23 | UDI0034 | 3537.8 | 3.5 | 56716548 | 41537739 | 15741155 | 38 |
| **VM-15** | 0.25 | 1 | 0 | 1.73 | 20 | 15.11 | 13 | 0.85 | UDI0037 | 2713.9999 | 2.7 | 30001708 | 28037599 | 6317335 | 23 |
| **VM-17** | 0.25 | 1 | 0 | 3.38 | 20 | 15.36 | 13 | 11.4 | UDI0039 | 5957.1541 | 34.0 | 84386458 | 74052027 | 16790792 | 23 |
| **VM-19** | 0.25 | 1 | 0 | 1.9 | 20 | 17.05 | 15 | 10.2 | UDI0040 | 7286.8692 | 37.2 | 56144636 | 45483823 | 14377853 | 32 |
| **VM-22** | 0.25 | 1 | 0 | 0.712 | 20 | 21 | 19 | 5.94 | UDI0042 | 9107.3487 | 27.0 | 37914131 | 20816199 | 18029818 | 87 |
| **VM-24** | 0.25 | 1 | 0 | 1.68 | 20 | 23.19 | 21 | 9.78 | UDI0043 | 10480.2508 | 51.2 | 54675567 | 40668078 | 38706373 | 95 |
| **VM-26** | 0.25 | 1 | 0 | 0.846 | 20 | 21.24 | 19 | 2.8 | UDI0045 | 8989.3165 | 12.6 | 45987295 | 34422855 | 32435510 | 94 |
| **VM-28** | 0.25 | 1 | 0 | 0.718 | 20 | 20.25 | 18 | 5.52 | UDI0047 | 7241.6254 | 20.0 | 73635745 | 53146920 | 48685483 | 92 |
| **CTRL-2** | - | 1 | 0 | Too Low | 20 | 20.31 | 21 | 20.4 | UDI0048 | 8579.2038 | 87.5 | 20535075 | 6849175 | 13685900 | 66 |
| **LIB-CTRL-1** | . | - | - | - | 20 | 22.42 | 21 | 0.178 | UDI0050 | 87.5098 | 0.1 | 900421 | 180807 | 719614 | 80 |

**Supplementary Table 1 (sheet 1). Sequenced samples.**

| SampleID | Volume (g) | Zymo Clean-up | MinElute Clean-up | Qubit | Volume DNA(ul) | qPCR -Ct | Index PCRCycles | Qubit | Index Used | Bioanalyser Molarity(pm/l) | Library Molarity | Notes |
|---|---|---|---|---|---|---|---|---|---|---|---|---|
| **VM-7** | 0.25 | 1 | 0 | 2.74 | 20 | 21.18 | 19 | 0.77 | UDI0032 | 57.63 | 0.1 | failed |
| **VM-16** | 0.25 | 1 | 0 | 8.16 | 20 | 24.19 | 22 | 0.816 | UDI0038 | 967.61 | 1 | failed |
| **VM-20** | 0.25 | 1 | 0 | TooLow | 20 | 19.26 | 17 | 4.58 | UDI0041 | 7864.906 | 18 | adapter dimers |
| **VM-25** | 0.25 | 1 | 0 | 2 | 20 | 23.36 | 21 | 0.308 | UDI0044 | 58.69 | 0 | failed |
| **VM-1** | 0.25 | 1 | 1 | 3.50 | 10 | 23.30 | 22 | 0.30 | UDI0001 | 130.2 | 0 | failed |
| **VM-5** | 0.25 | 1 | 1 | 7.82 | 20 | 11.36 | 10 | 7.36 | UDI0003 | 8,934.70 | 35.738,8 | ok |
| **VM-9** | 0.25 | 1 | 1 | 13.5 | 5 | NoCt | 26 | 0.25 | UDI0004 | 0 | 0 | failed |
| **VM-18** | 0.25 | 1 | 0 | 1.88 | 20 | 10.96 | 10 | 2.74 | UDI0005 | 9,756.10 | 19.512,2 | adapter dimers |
| **VM-21** | 0.25 | 1 | 0 | 0.39 | 20 | 18.82 | 18 | 7.42 | UDI0006 | 13,100.00 | 52.400,00 | adaper dimers |
| **VM-23** | 0.25 | 1 | 1 | 1.03 | 20 | 18.56 | 18 | 0.27 | UDI0007 | 64 | 0 | failed |
| **VM-27** | 0.25 | 1 | 0 | 0.72 | 10 | 20.08 | 19 | 5.24 | UDI0008 | 6,722.00 | 26.888,00 | adapter-dimers |

**Supplementary Table 1 (sheet 2). Samples not sequenced**.
