## Supplementary material for "Early Pastoralism in Central European Forests: Insights from Ancient Environmental Genomics": Dataset S2

| tax id | tax name | tax rank | sample | N_reads | N_alignments | Bayesian z | Bayesian D_max | Bayesian D_max_std | mean_L | std_L |
|---|---|---|---|---|---|---|---|---|---|---|
| 9822 | Sus | genus | VM-11 | 19107 | 579432 | 7.376597 | 0.12539655 | 0.018326212 | 50.3773 | 15.20254 |
| 9822 | Sus | genus | VM-14 | 4319 | 411690 | 3.234768 | 0.053543355 | 0.012386373 | 42.13939 | 11.71935 |
| 9900 | Bison | genus | VM-14 | 620 | 11038 | 2.1509 | 0.055930864 | 0.028148236 | 42.09193 | 11.0488 |
| 9903 | Bos | genus | VM-14 | 1158 | 17711 | 2.278015 | 0.07434724 | 0.026416106 | 43.67358 | 12.7568 |
| 9935 | Ovis | genus | VM-14 | 879 | 34824 | 2.903726 | 0.089806005 | 0.028953828 | 45.16724 | 13.07526 |
| 9605 | Homo | genus | VM-15 | 1127 | 11640 | 3.170284 | 0.13593441 | 0.031629723 | 60.92635 | 16.97385 |
| 9822 | Sus | genus | VM-15 | 6536 | 444481 | 3.414994 | 0.07387795 | 0.016445031 | 60.10205 | 17.69219 |
| 9900 | Bison | genus | VM-15 | 504 | 8027 | 2.337582 | 0.09852042 | 0.038240712 | 49.61508 | 15.41217 |
| 9903 | Bos | genus | VM-15 | 948 | 15375 | 2.568237 | 0.11980395 | 0.03400362 | 54.96097 | 16.9672 |
| 9922 | Capra | genus | VM-15 | 801 | 82992 | 3.346514 | 0.09556448 | 0.030669581 | 53.54682 | 16.6221 |
| 9935 | Ovis | genus | VM-15 | 715 | 44325 | 3.995386 | 0.13392974 | 0.03503174 | 55.62937 | 16.78439 |
| 7106 | Spodoptera | genus | VM-17 | 160 | 213 | 2.089201 | 0.13296323 | 0.0783851 | 65.9625 | 18.01258 |
| 9822 | Sus | genus | VM-17 | 8793 | 479379 | 4.522457 | 0.05422004 | 0.010848547 | 59.18833 | 18.34578 |
| 9869 | Rangifer | genus | VM-17 | 2319 | 4337 | 4.607052 | 0.06632892 | 0.016985565 | 50.6326 | 16.68363 |
| 9900 | Bison | genus | VM-17 | 674 | 9432 | 4.045144 | 0.1133476 | 0.0350166 | 47.23294 | 15.08294 |
| 9903 | Bos | genus | VM-17 | 1266 | 14009 | 4.212742 | 0.16116342 | 0.03232278 | 50.15877 | 16.25794 |

| | | | | | | | | | | |
|---|---|---|---|---|---|---|---|---|---|---|
| 9918 | Bubalus | genus | VM-17 | 346 | 756 | 2.825442 | 0.13437642 | 0.052783545 | 48.05203 | 14.78719 |
| 9922 | Capra | genus | VM-17 | 3763 | 92452 | 5.200697 | 0.10111978 | 0.01841629 | 50.59527 | 15.89242 |
| 9935 | Ovis | genus | VM-17 | 53698 | 2463482 | 6.818388 | 0.11670803 | 0.012212124 | 55.41134 | 17.06771 |
| 9822 | Sus | genus | VM-19 | 157 | 4857 | 2.203991 | 0.13774426 | 0.07669815 | 44.35669 | 13.06084 |
| 9900 | Bison | genus | VM-19 | 790 | 10862 | 3.758634 | 0.09677571 | 0.03126048 | 44.51266 | 13.03797 |
| 9903 | Bos | genus | VM-19 | 1258 | 11723 | 3.874253 | 0.11230852 | 0.03214293 | 49.48331 | 15.6312 |
| 9918 | Bubalus | genus | VM-19 | 151 | 312 | 3.895037 | 0.1623977 | 0.08222423 | 48.01987 | 14.27665 |
| 9922 | Capra | genus | VM-19 | 791 | 33077 | 3.376575 | 0.10092918 | 0.032522332 | 47.49684 | 14.38026 |
| 9935 | Ovis | genus | VM-19 | 7122 | 336249 | 10.36418 | 0.11785975 | 0.01576554 | 53.78377 | 16.19847 |
| 9922 | Capra | genus | VM-24 | 568 | 66317 | 2.105745 | 0.13240187 | 0.043794904 | 53.8257 | 15.51172 |

**Supplementary Table 2 (sheet 1)**. metaDMG output for animal reads (Metazoa). Due to space limitations only the main statistics are shown.

| Tax id | Tax name | Tax rank | sample | N_reads | N_alignments | Bayesian z | Bayesian D_max | Bayesian D_max_std | mean_L | std_L |
|---|---|---|---|---|---|---|---|---|---|---|
| 9869 | Rangifer | genus | VM-14 | 168 | 270 | 0.7879365 | 0.04522432 | 0.04858463 | 43.023811 | 13.1703759 |
| 9922 | Capra | genus | VM-14 | 730 | 111098 | 1.3281344 | 0.06782559 | 0.03074879 | 43.335617 | 12.5138101 |
| 9869 | Rangifer | genus | VM-15 | 125 | 185 | 1.1341809 | 0.08351998 | 0.07393324 | 53.335999 | 16.5549327 |
| 77156 | Dendroctonus | genus | VM-15 | 294 | 340 | 0.7830976 | 0.042092 | 0.03430296 | 68.363945 | 15.451155 |
| 9605 | Homo | genus | VM-17 | 2321 | 34072 | 2.1985846 | 0.01678161 | 0.00940506 | 69.828522 | 14.9865749 |

| | | | | | | | | | | |
|---|---|---|---|---|---|---|---|---|---|---|
| 9871 | Odocoileus | genus | VM-17 | 125 | 140 | 0.81171286 | 0.08361646 | 0.07157334 | 46.368 | 13.874554 |
| 77156 | Dendroctonus | genus | VM-17 | 120 | 145 | 1.4118043 | 0.13633937 | 0.09051972 | 62.450001 | 17.7907256 |
| 9605 | Homo | genus | VM-19 | 392 | 8558 | -0.1889077 | 0.0112201 | 0.01807768 | 64.553574 | 16.8799165 |
| 9869 | Rangifer | genus | VM-19 | 438 | 572 | 1.3644423 | 0.06635018 | 0.04386491 | 45.226028 | 13.2802579 |
| 9605 | Homo | genus | VM-22 | 439 | 5206 | -0.4575679 | 0.00655198 | 0.01419727 | 62.006832 | 16.7390055 |
| 9822 | Sus | genus | VM-3 | 8610 | 1116669 | 4.399277 | 0.02620799 | 0.00638303 | 42.87294 | 11.5788946 |
| 28609 | Rhagoletis | genus | VM-3 | 186 | 201 | -1.0005367 | 0.01240317 | 0.02469642 | 51.037633 | 15.7018155 |
| 9605 | Homo | genus | VM-26 | 115 | 983 | -2.0146244 | 0.00948942 | 0.02710047 | 90.930435 | 41.2115366 |
| 9605 | Homo | genus | VM-28 | 764 | 13408 | 0.08075919 | 0.00701261 | 0.01055238 | 77.316757 | 43.2350752 |

**Supplementary Table 2 (sheet 2)**. metaDMG output for discarded animal reads (Metazoa). Due to space limitations only the main statistics are shown.

| tax id | tax name | tax rank | sample | N_reads | N_alignments | Bayesian z | Bayesian D_max | Bayesian_D_max_std | mean_L | std_L |
|---|---|---|---|---|---|---|---|---|---|---|
| 4022 | Acer | genus | VM-14 | 1262 | 12296 | 2.1940873 | 0.05854126 | 0.02422669 | 42.649764 | 11.94 |
| 13384 | Calluna | genus | VM-14 | 315 | 3559 | 2.1756625 | 0.05138103 | 0.03301592 | 44.387302 | 11.54 |
| 24735 | Ulmus | genus | VM-14 | 3175 | 70960 | 3.2755458 | 0.0627775 | 0.01446115 | 44.033072 | 11.03 |
| 3287 | Dryopteris | genus | VM-15 | 216 | 1134 | 2.4267113 | 0.09957606 | 0.05370223 | 53.893519 | 16.64 |
| 3328 | Picea | genus | VM-15 | 1369 | 1951 | 2.5751944 | 0.05142983 | 0.0215426 | 55.88824 | 17.08 |
| 3445 | Ranunculus | genus | VM-15 | 188 | 1559 | 3.2121863 | 0.13284001 | 0.065914 | 56.053192 | 16.86 |
| 3511 | Quercus | genus | VM-15 | 662 | 16493 | 2.7592194 | 0.08312589 | 0.03303182 | 50.543806 | 16.01 |
| 4564 | Triticum | genus | VM-15 | 1822 | 12890 | 3.08302 | 0.07946138 | 0.02123224 | 60.141054 | 17.46 |

| | | | | | | | | | | |
|---|---|---|---|---|---|---|---|---|---|---|
| 21024 | Fagus | genus | VM-15 | 470 | 918 | 4.2539935 | 0.09421362 | 0.03460744 | 56.274469 | 16.77 |
| 24735 | Ulmus | genus | VM-15 | 1251 | 21365 | 2.4682846 | 0.07080207 | 0.0258441 | 49.217427 | 15.18 |
| 72171 | Ziziphus | genus | VM-15 | 260 | 359 | 2.373453 | 0.09174149 | 0.04389537 | 68.957695 | 15.93 |
| 3287 | Dryopteris | genus | VM-17 | 628 | 3447 | 2.2405403 | 0.07627627 | 0.03386837 | 47.113058 | 15.10 |
| 3853 | Lathyrus | genus | VM-17 | 350 | 6134 | 2.301905 | 0.06891531 | 0.04506043 | 46.671428 | 14.11 |
| 4022 | Acer | genus | VM-17 | 1903 | 18229 | 2.4083128 | 0.05282589 | 0.02052292 | 48.63689 | 15.61 |
| 24735 | Ulmus | genus | VM-17 | 4206 | 75716 | 3.6909297 | 0.05437997 | 0.01401585 | 47.564433 | 15.77 |
| 38871 | Fraxinus | genus | VM-17 | 389 | 2167 | 2.1873164 | 0.08179359 | 0.04610297 | 45.632389 | 14.49 |
| 53158 | Lamium | genus | VM-17 | 6938 | 154478 | 4.0131145 | 0.08609603 | 0.01433044 | 45.578986 | 14.02 |
| 3511 | Quercus | genus | VM-19 | 136 | 7345 | 2.0931482 | 0.12232672 | 0.07861342 | 45.676471 | 12.93 |
| 4022 | Acer | genus | VM-19 | 5669 | 48693 | 8.697742 | 0.07074951 | 0.01460256 | 48.411362 | 14.74 |
| 24735 | Ulmus | genus | VM-19 | 1228 | 23902 | 2.4643123 | 0.05507202 | 0.02270807 | 45.274429 | 13.35 |
| 38871 | Fraxinus | genus | VM-19 | 217 | 1084 | 2.2421446 | 0.08823524 | 0.05892783 | 46.281105 | 13.21 |

**Supplementary Table 2 (sheet 3)**. metaDMG output for plant reads (Viridiplantae). Due to space limitations only the main statistics are shown.

| Tax id | Tax name | Tax rank | sample | N_reads | N_alignments | Bayesian z | Bayesian D_max | Bayesian D_max_std | mean_L | std_L |
|---|---|---|---|---|---|---|---|---|---|---|
| 3445 | Ranunculus | genus | VM-14 | 278 | 3224 | -0.5214261 | 0.02079102 | 0.03261297 | 42.564747 | 11.2837648 |

| | | | | | | | | | | |
|---|---|---|---|---|---|---|---|---|---|---|
| 3500 | Urtica | genus | VM-14 | 108 | 1322 | -0.7781616 | 0.03987466 | 0.09241644 | 48.277779 | 15.0438715 |
| 15740 | Phalaris | genus | VM-14 | 202 | 8004 | 1.072083 | 0.05966582 | 0.0476965 | 43.217823 | 11.3717318 |
| 22868 | Anemone | genus | VM-14 | 129 | 3645 | 1.180413 | 0.06120521 | 0.05902736 | 43.16279 | 10.8508143 |
| 23137 | Agrimonia | genus | VM-14 | 102 | 3482 | -1.680038 | 0.02061185 | 0.06939499 | 42.803921 | 10.7229658 |
| 23242 | Chrysosplenium | genus | VM-14 | 252 | 707 | 1.0257076 | 0.03366343 | 0.02607768 | 41.876983 | 11.645597 |
| 125026 | Elatine | genus | VM-14 | 119 | 139 | -2.0306985 | 0.01339231 | 0.03330917 | 46.394958 | 14.791098 |
| 192333 | Myosotis | genus | VM-14 | 199 | 232 | 1.1374393 | 0.0323092 | 0.0311262 | 46.839198 | 14.4509124 |
| 694344 | Betonica | genus | VM-14 | 203 | 2134 | -0.0810782 | 0.02454943 | 0.03771403 | 43.147785 | 12.0834439 |
| 4022 | Acer | genus | VM-15 | 455 | 3947 | 1.0676551 | 0.06270309 | 0.04080681 | 51.956045 | 16.6662678 |
| 4051 | Hedera | genus | VM-15 | 143 | 3264 | 0.43401107 | 0.06781886 | 0.07371197 | 55.370628 | 16.3318561 |
| 13384 | Calluna | genus | VM-15 | 121 | 1133 | 0.8012103 | 0.05169896 | 0.05582635 | 48.05785 | 15.8720792 |
| 13398 | Carex | genus | VM-15 | 274 | 476 | 1.5343285 | 0.04787089 | 0.03616064 | 58.532846 | 15.9216659 |
| 15297 | Agrostis | genus | VM-15 | 142 | 182 | 1.0034207 | 0.05825968 | 0.04846759 | 50.169014 | 15.4856937 |

| | | | | | | | | | | |
|---|---|---|---|---|---|---|---|---|---|---|
| 16718 | Juglans | genus | VM-15 | 142 | 3438 | -0.2219879 | 0.03215439 | 0.05159994 | 51.823944 | 16.6485081 |
| 23242 | Chrysosplenium | genus | VM-15 | 162 | 425 | 0.20281959 | 0.02744941 | 0.0356737 | 59.84568 | 18.2562383 |
| 39200 | Hippuris | genus | VM-15 | 289 | 501 | 1.1987033 | 0.09854195 | 0.04712093 | 65.68512 | 17.2133142 |
| 3328 | Picea | genus | VM-17 | 328 | 457 | 1.2933365 | 0.04896877 | 0.044335 | 43.021341 | 12.3448297 |
| 3337 | Pinus | genus | VM-17 | 106 | 233 | -0.6939515 | 0.02767875 | 0.04839742 | 52.320756 | 17.2606912 |
| 3445 | Ranunculus | genus | VM-17 | 244 | 2381 | 1.2571186 | 0.05204562 | 0.04684252 | 47.643443 | 15.0870981 |
| 3500 | Urtica | genus | VM-17 | 101 | 1355 | 0.7309036 | 0.08982276 | 0.1061109 | 44.752477 | 14.3278955 |
| 3504 | Betula | genus | VM-17 | 109 | 209 | -0.0055121 | 0.04308143 | 0.06949379 | 48.587155 | 16.8675351 |
| 3511 | Quercus | genus | VM-17 | 414 | 23793 | 1.4386088 | 0.0677105 | 0.04290636 | 47.649759 | 15.852215 |
| 3984 | Mercurialis | genus | VM-17 | 207 | 1844 | 1.393858 | 0.0630658 | 0.05214481 | 49.734299 | 14.5485978 |
| 4034 | Oxalis | genus | VM-17 | 108 | 1971 | 0.3154346 | 0.04276439 | 0.06120668 | 46.203705 | 14.6357872 |
| 4051 | Hedera | genus | VM-17 | 2446 | 51568 | 1.3729725 | 0.06671246 | 0.02337899 | 54.271465 | 16.910616 |
| 13328 | Achillea | genus | VM-17 | 165 | 1054 | 0.87543696 | 0.07660942 | 0.07702182 | 47.969696 | 15.5724663 |

| | | | | | | | | | | |
|---|---|---|---|---|---|---|---|---|---|---|
| 13384 | Calluna | genus | VM-17 | 419 | 4520 | 1.5333593 | 0.0541065 6 | 0.0321990 3 | 46.25537 1 | 15.0981 136 |
| 13398 | Carex | genus | VM-17 | 317 | 2485 | 1.3697034 | 0.0677100 6 | 0.0409616 1 | 52.74763 4 | 15.1928 63 |
| 22868 | Anemone | genus | VM-17 | 122 | 2131 | 1.2852796 | 0.1060289 3 | 0.0936671 1 | 48.22950 9 | 14.9543 708 |
| 23242 | Chrysospleni um | genus | VM-17 | 358 | 1036 | 0.770098 45 | 0.0384934 4 | 0.0308945 9 | 49.29888 4 | 17.0437 41 |
| 25168 | Galium | genus | VM-17 | 476 | 2887 | 1.5055004 | 0.0710463 9 | 0.0394153 6 | 46.19958 1 | 13.8201 892 |
| 32071 | Asplenium | genus | VM-17 | 359 | 2417 | 1.6010982 | 0.0564528 | 0.0353078 9 | 46.62674 1 | 14.5531 686 |
| 39200 | Hippuris | genus | VM-17 | 192 | 236 | 0.628837 76 | 0.0476500 5 | 0.0394598 1 | 53.75520 7 | 17.3249 511 |
| 66655 | Durio | genus | VM-17 | 172 | 534 | -0.1554813 | 0.0385042 2 | 0.0502878 1 | 45.27325 4 | 14.6269 149 |
| 72171 | Ziziphus | genus | VM-17 | 176 | 261 | 0.3089143 | 0.0410518 2 | 0.0462298 8 | 62.06818 | 17.5628 614 |
| 12502 6 | Elatine | genus | VM-17 | 171 | 203 | 0.357492 | 0.0383925 1 | 0.0396785 1 | 52.38011 6 | 17.4763 434 |
| 19233 3 | Myosotis | genus | VM-17 | 220 | 264 | 0.924110 83 | 0.0349531 5 | 0.0315327 5 | 53.50454 7 | 18.4248 036 |
| 36834 0 | Milium | genus | VM-17 | 221 | 3453 | 1.0782028 | 0.0583235 7 | 0.0486186 7 | 49.48416 1 | 15.3144 376 |
| 69434 4 | Betonica | genus | VM-17 | 338 | 3273 | 1.0845356 | 0.0576294 | 0.0468983 8 | 46.21301 7 | 14.9045 276 |

| | | | | | | | | | | |
|---|---|---|---|---|---|---|---|---|---|---|
| 1072081 | Jenufa | genus | VM-17 | 201 | 442 | -1.0216042 | 0.02058155 | 0.04204299 | 62.000001 | 17.7842316 |
| 3287 | Dryopteris | genus | VM-19 | 261 | 1075 | 0.5434875 | 0.0570695 | 0.05038835 | 47.149426 | 14.5213662 |
| 3504 | Betula | genus | VM-19 | 108 | 126 | -0.2098737 | 0.04515034 | 0.07501492 | 46.75 | 14.7283185 |
| 3515 | Alnus | genus | VM-19 | 143 | 259 | -0.3462719 | 0.03887081 | 0.05905734 | 45.58042 | 13.0409062 |
| 4051 | Hedera | genus | VM-19 | 802 | 18773 | 1.6926988 | 0.05253967 | 0.03273097 | 48.790524 | 13.9189653 |
| 4520 | Lolium | genus | VM-19 | 127 | 1117 | 0.6251078 | 0.05973695 | 0.06022552 | 51.385826 | 15.1049399 |
| 23242 | Chrysosplenium | genus | VM-19 | 158 | 431 | 0.9567373 | 0.05235249 | 0.04704672 | 44.955698 | 13.8374138 |
| 32071 | Asplenium | genus | VM-19 | 779 | 3994 | 1.6067332 | 0.04467059 | 0.02939702 | 52.243901 | 16.517393 |
| 39200 | Hippuris | genus | VM-19 | 107 | 153 | 0.12106786 | 0.03921264 | 0.05026181 | 49.2243 | 16.061551 |
| 53158 | Lamium | genus | VM-19 | 336 | 10141 | 1.6028959 | 0.0835991 | 0.05045437 | 42.092261 | 12.3112425 |
| 66655 | Durio | genus | VM-19 | 101 | 223 | 1.0293341 | 0.09524754 | 0.08932153 | 45.009899 | 13.6903787 |
| 192333 | Myosotis | genus | VM-19 | 151 | 157 | 1.4894193 | 0.05535323 | 0.04228227 | 48.218544 | 16.3937304 |
| 24735 | Ulmus | genus | VM-22 | 327 | 6927 | 0.61970145 | 0.04084806 | 0.03700938 | 44.14373 | 13.1105226 |

| | | | | | | | | | | |
|---|---|---|---|---|---|---|---|---|---|---|
| 1072081 | Jenufa | genus | VM-24 | 353 | 776 | 0.7005385 | 0.03971831 | 0.04454596 | 55.841359 | 17.0594182 |
| 3573 | Silene | genus | VM-2 | 116 | 122 | -1.745035 | 0.01302676 | 0.03457943 | 48.043105 | 12.0414681 |
| 284690 | Triantha | genus | VM-2 | 509 | 1575 | 0.8463814 | 0.02215645 | 0.01881635 | 47.966602 | 12.8990438 |
| 3328 | Picea | genus | VM-3 | 241 | 378 | 0.29735297 | 0.03132053 | 0.03167525 | 45.80498 | 11.7941458 |
| 3511 | Quercus | genus | VM-3 | 389 | 111498 | -0.2342472 | 0.01337964 | 0.02252503 | 47.473009 | 12.1200733 |
| 13384 | Calluna | genus | VM-3 | 245 | 5382 | 0.89327735 | 0.03512275 | 0.0327859 | 46.893879 | 13.4996507 |
| 13398 | Carex | genus | VM-3 | 425 | 721 | -0.9887556 | 0.00979382 | 0.01813254 | 51.764707 | 15.8764442 |
| 23138 | Alchemilla | genus | VM-3 | 238 | 254 | -0.3747561 | 0.01350953 | 0.02318729 | 51.596638 | 16.6284969 |
| 23242 | Chrysosplenium | genus | VM-3 | 105 | 247 | -0.0128806 | 0.02027861 | 0.03193466 | 44.304762 | 13.0408056 |
| 46334 | Scirpus | genus | VM-3 | 219 | 340 | 1.5189608 | 0.03275859 | 0.02729484 | 57.744293 | 15.5839235 |
| 284690 | Triantha | genus | VM-3 | 691 | 1729 | -0.1411192 | 0.0102599 | 0.01340884 | 63.32272 | 15.5316543 |
| 1072081 | Jenufa | genus | VM-26 | 180 | 392 | 0.9478649 | 0.06927855 | 0.07441808 | 46.122221 | 13.7295586 |

**Supplementary Table 2 (sheet 4)**. metaDMG output for discarded plant reads (Viridiplantae). Due to space limitations only the main statistics are shown.
