## Supplementary material for "Early Pastoralism in Central European Forests: Insights from Ancient Environmental Genomics": Dataset S3

| #rname | startpos | endpos | numreads | covbases | coverage | meandepth | meanbaseq | meanmapq | Sample_ID |
|---|---|---|---|---|---|---|---|---|---|
| Bos_taurus_haplogroup_T | 1 | 16341 | 99 | 4363 | 26.70 | 0.322196 | 39 | 41 | VM_15 |
| Bos_taurus_haplogroup_Q | 1 | 16340 | 99 | 4406 | 26.96 | 0.322277 | 39 | 41 | VM_15 |
| Bos_primigenius | 1 | 16339 | 95 | 4290 | 26.26 | 0.309688 | 39 | 41 | VM_15 |
| Bos_taurus_haplogroup_R | 1 | 16342 | 87 | 4050 | 24.78 | 0.286685 | 39 | 41 | VM_15 |
| Bos_Indicus_I_outgroup | 1 | 16341 | 76 | 3534 | 21.63 | 0.245028 | 39 | 41 | VM_15 |
| Bos_taurus_haplogroup_T | 1 | 16341 | 52 | 2125 | 13.00 | 0.140261 | 39 | 41 | VM_17 |
| Bos_taurus_haplogroup_Q | 1 | 16340 | 51 | 2082 | 12.74 | 0.137638 | 39 | 41 | VM_17 |
| Bos_primigenius | 1 | 16339 | 49 | 1954 | 11.96 | 0.129812 | 39 | 41 | VM_17 |
| Bos_taurus_haplogroup_R | 1 | 16342 | 48 | 1978 | 12.10 | 0.129972 | 39 | 41 | VM_17 |
| Bos_taurus_haplogroup_T | 1 | 16341 | 42 | 1655 | 10.13 | 0.10544 | 39 | 41 | VM_14 |
| Bos_taurus_haplogroup_Q | 1 | 16340 | 42 | 1655 | 10.13 | 0.105447 | 39 | 41 | VM_14 |
| Bos_Indicus_I_outgroup | 1 | 16341 | 40 | 1623 | 9.93 | 0.10593 | 39 | 41 | VM_17 |
| Bos_primigenius | 1 | 16339 | 40 | 1558 | 9.54 | 0.0995165 | 39 | 41 | VM_14 |
| Bos_taurus_haplogroup_R | 1 | 16342 | 37 | 1469 | 8.99 | 0.0932566 | 39 | 41 | VM_14 |
| Bos_Indicus_I_outgroup | 1 | 16341 | 32 | 1292 | 7.91 | 0.0824307 | 39 | 41 | VM_14 |
| Bos_taurus_haplogroup_Q | 1 | 16340 | 26 | 1201 | 7.35 | 0.0758874 | 38 | 41 | VM_19 |
| Bos_primigenius | 1 | 16339 | 25 | 1148 | 7.03 | 0.0720974 | 38 | 41 | VM_19 |
| Bos_taurus_haplogroup_T | 1 | 16341 | 25 | 1124 | 6.88 | 0.0711707 | 38 | 41 | VM_19 |
| Bos_taurus_haplogroup_R | 1 | 16342 | 24 | 1081 | 6.61 | 0.0679843 | 38 | 40 | VM_19 |
| Bos_Indicus_I_outgroup | 1 | 16341 | 22 | 968 | 5.92 | 0.0610734 | 38 | 40 | VM_19 |
| Ovis_aries_Haplogroup_B | 1 | 16620 | 84 | 3269 | 19.67 | 0.277798 | 39 | 41 | VM_17 |

| | | | | | | | | | |
|---|---|---|---|---|---|---|---|---|---|
| Ovis_aries_musimon | 1 | 16616 | 83 | 3246 | 19.54 | 0.274615 | 39 | 41 | VM_17 |
| Ovis_aries_Haplogroup_A | 1 | 16622 | 75 | 3138 | 18.88 | 0.250451 | 39 | 41 | VM_17 |
| Ovis_aries_Haplogroup_D | 1 | 16616 | 69 | 2944 | 17.72 | 0.235255 | 39 | 41 | VM_17 |
| KP981380.1_Ovis_aries_East_Asia | 1 | 16613 | 67 | 2843 | 17.11 | 0.222777 | 39 | 41 | VM_17 |
| KF312238.2_Ovis_orientalis_ophion | 1 | 16620 | 64 | 2847 | 17.13 | 0.214621 | 39 | 41 | VM_17 |
| Ovis_aries_Haplogroup_E | 1 | 16620 | 61 | 2776 | 16.70 | 0.20373 | 39 | 41 | VM_17 |
| Ovis_aries_Haplogroup_C | 1 | 16624 | 60 | 2720 | 16.36 | 0.204343 | 39 | 41 | VM_17 |
| Ovis_vignei | 1 | 16697 | 31 | 1378 | 8.25 | 0.0975025 | 39 | 40 | VM_17 |
| Ovis_ammon | 1 | 16615 | 30 | 1261 | 7.59 | 0.0840807 | 39 | 40 | VM_17 |
| Ovis_aries_Haplogroup_B | 1 | 16620 | 25 | 1090 | 6.56 | 0.0774368 | 40 | 41 | VM_19 |
| Ovis_aries_musimon | 1 | 16616 | 24 | 1041 | 6.27 | 0.0745065 | 40 | 41 | VM_19 |
| Ovis_aries_Haplogroup_A | 1 | 16622 | 21 | 987 | 5.94 | 0.0676814 | 40 | 40 | VM_19 |
| Ovis_aries_Haplogroup_D | 1 | 16616 | 18 | 817 | 4.92 | 0.0527203 | 40 | 41 | VM_19 |
| KP981380.1_Ovis_aries_East_Asia | 1 | 16613 | 17 | 825 | 4.97 | 0.0564618 | 40 | 41 | VM_19 |
| Ovis_aries_Haplogroup_E | 1 | 16620 | 16 | 725 | 4.36 | 0.0471721 | 40 | 41 | VM_19 |
| KF312238.2_Ovis_orientalis_ophion | 1 | 16620 | 14 | 671 | 4.04 | 0.0425391 | 40 | 41 | VM_19 |
| Ovis_aries_Haplogroup_C | 1 | 16624 | 13 | 574 | 3.45 | 0.0366939 | 40 | 41 | VM_19 |
| Ovis_ammon | 1 | 16615 | 9 | 294 | 1.77 | 0.0198616 | 40 | 40 | VM_19 |
| Ovis_vignei | 1 | 16697 | 7 | 436 | 2.61 | 0.0282686 | 40 | 41 | VM_19 |
| KP981380.1_Ovis_aries_East_Asia | 1 | 16613 | 3 | 81 | 0.49 | 0.0048757 | 40 | 40 | VM_15 |
| Ovis_aries_Haplogroup_B | 1 | 16620 | 1 | 81 | 0.49 | 0.00487365 | 40 | 40 | VM_15 |
| Ovis_aries_Haplogroup_A | 1 | 16622 | 1 | 81 | 0.49 | 0.00487306 | 40 | 40 | VM_15 |

| | | | | | | | | | |
|---|---|---|---|---|---|---|---|---|---|
| KR059146.1_Capra_hircus_outgroup | 1 | 16642 | 0 | 44 | 0.26 | 0.00264391 | 40 | 0 | VM_17 |
| Ovis_aries_Haplogroup_D | 1 | 16616 | 0 | 81 | 0.49 | 0.00487482 | 40 | 0 | VM_15 |
| Ovis_aries_musimon | 1 | 16616 | 0 | 81 | 0.49 | 0.00487482 | 40 | 0 | VM_15 |

**Supplementary Table 3 (sheet 1)**. Mapping statistics of the taxa used for phylogenetic analyses.

| #rname | startpos | endpos | numreads | covbases | coverage | meandepth | meanbaseq | meanmapq | Sample_ID |
|---|---|---|---|---|---|---|---|---|---|
| NC_012920.1_Homo_sapiens_mitochondrion_complete_genome | 1 | 16569 | 12 | 678 | 4.09 | 0.0409198 | 39 | 40 | VM-15 |
| Capra_aegagrus_Consensus | 1 | 16642 | 32 | 1197 | 7.19 | 0.082442 | 40 | 41 | VM-17 |
| Capra_Hircus_HaplogroupA_Consensus | 1 | 16647 | 32 | 1197 | 7.19 | 0.0824173 | 40 | 41 | VM-17 |
| NC_020683.1_Capra_caucasica_isolate_MA4105_mitochondrion_complete_genome | 1 | 16624 | 28 | 1050 | 6.32 | 0.0720645 | 39 | 40 | VM-17 |
| Capra_aegagrus_Consensus | 1 | 16642 | 14 | 671 | 4.03 | 0.0495133 | 39 | 41 | VM-15 |
| Capra_Hircus_HaplogroupA_Consensus | 1 | 16647 | 14 | 668 | 4.01 | 0.0480567 | 39 | 41 | VM-15 |
| NC_020683.1_Capra_caucasica_isolate_MA4105_mitochondrion_complete_genome | 1 | 16624 | 9 | 398 | 2.39 | 0.0318816 | 39 | 41 | VM-15 |
| Capra_aegagrus_Consensus | 1 | 16642 | 4 | 150 | 0.90 | 0.00901334 | 38 | 41 | VM-19 |
| NC_020683.1_Capra_caucasica_isolate_MA4105_mitochondrion_complete_genome | 1 | 16624 | 4 | 150 | 0.90 | 0.0090231 | 38 | 41 | VM-19 |
| Capra_Hircus_HaplogroupA_Consensus | 1 | 16647 | 4 | 150 | 0.90 | 0.00901063 | 38 | 41 | VM-19 |
| Capra_aegagrus_Consensus | 1 | 16642 | 0 | 0 | 0 | 0 | 0 | 0 | VM-24 |
| NC_020683.1_Capra_caucasica_isolate_MA4105_ | 1 | 16624 | 0 | 0 | 0 | 0 | 0 | 0 | VM-24 |

| | | | | | | | | | |
|---|---|---|---|---|---|---|---|---|---|
| mitochondrion_complete_genome | | | | | | | | | |
| Capra_Hircus_HaplogroupA_Consensus | 1 | 16647 | 0 | 0 | 0 | 0 | 0 | 0 | VM-24 |
| NC_020623.1_Capra_ibex | 1 | 16716 | 0 | 0 | 0 | 0 | 0 | 0 | VM-15 |
| NC_020623.1_Capra_ibex | 1 | 16716 | 0 | 0 | 0 | 0 | 0 | 0 | VM-17 |
| NC_020623.1_Capra_ibex | 1 | 16716 | 0 | 0 | 0 | 0 | 0 | 0 | VM-19 |
| NC_020623.1_Capra_ibex | 1 | 16716 | 0 | 0 | 0 | 0 | 0 | 0 | VM-24 |
| Sus_scrofa_European_Wild_Boar | 1 | 18108 | 13 | 378 | 2.09 | 0.0319748 | 38 | 41 | VM-11 |
| NC_012095.1_Sus_domesticus | 1 | 17851 | 8 | 261 | 1.46 | 0.0172539 | 37 | 41 | VM-11 |
| Sus_scrofa_European_Wild_Boar | 1 | 18108 | 8 | 416 | 2.30 | 0.0229733 | 38 | 41 | VM-15 |
| NC_012095.1_Sus_domesticus | 1 | 17851 | 8 | 416 | 2.33 | 0.023304 | 38 | 40 | VM-15 |
| KY911729.1_Sus_celebensis_isolate_dINc_14028_D_loop_partial_sequence_mitochondrial | 1 | 1010 | 5 | 92 | 9.11 | 0.223762 | 36 | 41 | VM-11 |
| MW538280.1_Porcula_salvania_isolate_PH41_mitochondrion_complete_genome | 1 | 16741 | 4 | 90 | 0.53 | 0.010752 | 40 | 41 | VM-11 |
| Sus_scrofa_European_Wild_Boar | 1 | 18108 | 4 | 109 | 0.60 | 0.00773139 | 40 | 40 | VM-17 |
| NC_012095.1_Sus_domesticus | 1 | 17851 | 4 | 109 | 0.61 | 0.0078427 | 40 | 40 | VM-17 |
| MW538280.1_Porcula_salvania_isolate_PH41_mitochondrion_complete_genome | 1 | 16741 | 4 | 107 | 0.64 | 0.00824324 | 40 | 40 | VM-17 |
| Sus_scrofa_European_Wild_Boar | 1 | 18108 | 3 | 102 | 0.56 | 0.00563287 | 38 | 42 | VM-14 |
| NC_012095.1_Sus_domesticus | 1 | 17851 | 2 | 72 | 0.40 | 0.00403339 | 37 | 42 | VM-14 |
| MW538280.1_Porcula_salvania_isolate_PH41_mitochondrion_complete_genome | 1 | 16741 | 1 | 32 | 0.19 | 0.00191147 | 34 | 42 | VM-14 |
| MW538280.1_Porcula_salvania_isolate_PH41_mitochondrion_complete_genome | 1 | 16741 | 1 | 66 | 0.39 | 0.00394242 | 36 | 40 | VM-15 |

| | | | | | | | | | |
|---|---|---|---|---|---|---|---|---|---|
| Sus_scrofa_European_Wild_Boar | 1 | 18108 | 1 | 38 | 0.21 | 0.00209852 | 39 | 40 | VM-19 |
| NC_012095.1_Sus_domesticus | 1 | 17851 | 1 | 38 | 0.21 | 0.00212873 | 39 | 40 | VM-19 |
| MW538280.1_Porcula_salvania_isolate_PH41_mitochondrion_complete_genome | 1 | 16741 | 0 | 0 | 0 | 0 | 0 | 0 | VM-19 |
| KX898017.1_Bison_bonasus | 1 | 16325 | 12 | 471 | 2.88515 | 0.0288515 | 38.4 | 41 | VM-14 |
| KX898017.1_Bison_bonasus | 1 | 16325 | 36 | 1540 | 9.43338 | 0.104502 | 39.5 | 41.2 | VM-15 |
| KX898017.1_Bison_bonasus | 1 | 16325 | 19 | 705 | 4.31853 | 0.045023 | 39.5 | 40.7 | VM-17 |
| KX898017.1_Bison_bonasus | 1 | 16325 | 6 | 234 | 1.43338 | 0.0161715 | 38.5 | 40.7 | VM-19 |
| KM593920.1_Bison_priscus | 1 | 16318 | 11 | 401 | 2.45741 | 0.0245741 | 38.3 | 40.9 | VM-14 |
| KM593920.1_Bison_priscus | 1 | 16318 | 32 | 1254 | 7.68477 | 0.0910038 | 39.6 | 41.4 | VM-15 |
| KM593920.1_Bison_priscus | 1 | 16318 | 21 | 810 | 4.96384 | 0.0540507 | 39.4 | 40.7 | VM-17 |
| KM593920.1_Bison_priscus | 1 | 16318 | 8 | 362 | 2.21841 | 0.0240226 | 38.6 | 40.5 | VM-19 |
| MK995024.1_Bubalus_quarlesi | 1 | 16354 | 15 | 527 | 3.22245 | 0.0374832 | 39.6 | 40.9 | VM-17 |
| MK995024.1_Bubalus_quarlesi | 1 | 16354 | 2 | 89 | 0.544209 | 0.00544209 | 40.1 | 40 | VM-19 |
| MT237632.1_Bubalus_bubalis | 1 | 16358 | 3 | 119 | 0.727473 | 0.00727473 | 38.2 | 40 | VM-19 |
| MT237632.1_Bubalus_bubalis | 1 | 16358 | 12 | 463 | 2.83042 | 0.0298325 | 39 | 41 | VM-17 |

**Supplementary Table 3 (sheet 2)**. Mapping statistics of the taxa discarded for phylogenetic analyses.
