## Supplementary material for "Early Pastoralism in Central European Forests: Insights from Ancient Environmental Genomics": Dataset S4

| Group | Accession number | Sample location | Common name | Reference |
|---|---|---|---|---|
| Haplogroup A - Ovis aries | MF004242 | Iraqi Kurdistan Region | Hamdani mixed with Karadi | Mustafa e al. 2018 |
| | HM236175 | Australia | Merino | Meadows et al. 2011 |
| | HM236174 | Australia | Merino | Meadows et al. 2011 |
| | KF302445 | Italy | Comisana | Lancioni et al. 2013 |
| | KF302444 | Italy | Comisana | Lancioni et al. 2013 |
| | KF938325 | China | Qinhai Tibetan | Lv et al. 2015 |
| | KF938319 | Mongolia, China | Ujimqin | Lv et al. 2015 |
| | KU681220 | China | Suffolk | Niu et al. 2017 |
| | KU681207 | China | Suffolk | Niu et al. 2017 |
| | KU681209 | China | Suffolk | Niu et al. 2017 |
| | KU681181 | Garze, Tibetan Autonomous Prefecture | Garze Tibetan | Niu et al. 2017 |
| | KU681194 | Ngawa, Tibetan Qiang Autonomous Prefecture | Ngawa Tibetan | Niu et al. 2017 |
| | KF938336 | China | Tan | Lv et al. 2015 |
| | KF977847 | NA | Small-tailed han | Brahi et al. 2015 |
| Haplogroup B - Ovis aries | HM236176 | Turkey | Karakas | Meadows et al. 2011 |
| | KP702285 | NA | Small-tail hulunbuir | NA |
| | KU681196 | Ngawa, Tibetan Qiang Autonomous Prefecture | Ngawa Tibetan | Niu et al. 2017 |

| | | | | |
|---|---|---|---|---|
| | KF938335 | China | Lanzhou Large-tailed | Lv et al. 2015 |
| | KF938348 | Southeast Europe, Moldova | Karakul | Lv et al. 2015 |
| | KU681204 | China | Suffolk | Niu et al. 2017 |
| | KU681223 | China | Suffolk | Niu et al. 2017 |
| | KU681175 | Garze, Tibetan Autonomous Prefecture | Garze Tibetan | Niu et al. 2017 |
| | KU681176 | Garze, Tibetan Autonomous Prefecture | Garze Tibetan | Niu et al. 2017 |
| | KF938339 | Buryatia, Russia | Baidarak | Lv et al. 2015 |
| | KF938328 | China | Jingzhong | Lv et al. 2015 |
| | KU681212 | China | Suffolk | Niu et al. 2017 |
| | KF938351 | North Caucasian, Russia | Karachai | Lv et al. 2015 |
| | MF004245 | Iraqi Kurdistan Region | Karadi mixed with Awassi | Mustafa e al. 2018 |
| | KF938346 | Volga Region, Russia | Kuibyshev | Lv et al. 2015 |
| | KF302460 | Italy | Lacaune | Lancioni et al. 2013 |
| | KF302457 | Italy | Gentile di Puglia | Lancioni et al. 2013 |
| | KU681210 | China | Suffolk | Niu et al. 2017 |
| Haplogroup C - Ovis aries | HM236178 | Turkey | Karakas | Meadows et al. 2011 |
| | HM236179 | Turkey | Morkaraman | Meadows et al. 2011 |
| | KF938318 | China | Minxian Black Fur | Lv et al. 2015 |
| | KF938327 | Inner Mongolia, China | Hulun Buir | Lv et al. 2015 |

| | | | | |
|---|---|---|---|---|
| | KP998471 | Tibet | Huoba | Liu et al. 2016 |
| | KT148968 | Tibet | Oula | NA |
| | KU681187 | Garze, Tibetan Autonomous Prefecture | Garze Tibetan | Niu et al. 2017 |
| | KU681188 | Garze, Tibetan Autonomous Prefecture | Garze Tibetan | Niu et al. 2017 |
| | KU681189 | Garze, Tibetan Autonomous Prefecture | Garze Tibetan | Niu et al. 2017 |
| | KU681191 | Garze, Tibetan Autonomous Prefecture | Garze Tibetan | Niu et al. 2017 |
| | KU681201 | Ngawa, Tibetan Qiang Autonomous Prefecture | Ngawa Tibetan | Niu et al. 2017 |
| Haplogroup D - Ovis aries | HM236181 | Turkey | Morkaraman | Meadows et al. 2011 |
| | HM236180 | Turkey | Morkaraman | Meadows et al. 2011 |
| Haplogroup E - Ovis aries | HM236183 | Turkey | Tuj | Meadows et al. 2011 |
| | HM236182 | Israel | Awassi | Meadows et al. 2011 |
| Ovis orientalis ophion | KF312238 | Cyprus | Cyprus Mouflon | Sanna et al. 2015 |
| Ovis vignei | HM236186 | Kazakhstan | Urial | Meadows et al. 2011 |
| | HM236187 | Kazakhstan | Urial | Meadows et al. 2011 |
| | HM236189 | Kazakhstan | Urial | Meadows et al. 2011 |
| Ovis East Breed Shadong | KP981380 | China | Shadong | Fan et al. 2016 |
| Capra hircus (outgroup) | KR059146 | Albania | Mati | Colli et al. 2015 |
| Ovis Ammon | HM236188 | Kazakhstan | Argali | Meadows et al. 2011 |

| | | | | |
|---|---|---|---|---|
| Ovis Musimon | HM236184 | Germany | Mouflon | Meadows et al. 2011 |
| | HM236185 | Germany | Mouflon | Meadows et al. 2011 |
| Haplogroup T - Bos taurus | JN817321.1 | Egypt | Domiaty | Bonfiglio et al., 2012 |
| | JN817322.1 | Egypt | Domiaty | Bonfiglio et al., 2012 |
| | JN817323.1 | Egypt | Domiaty | Bonfiglio et al., 2012 |
| | JN817324.1 | Egypt | Domiaty | Bonfiglio et al., 2012 |
| | JN817325.1 | Egypt | Menofi | Bonfiglio et al., 2012 |
| | JN817326.1 | Egypt | Menofi | Bonfiglio et al., 2012 |
| | JN817329.1 | Egypt | Menofi | Bonfiglio et al., 2012 |
| | KT184462.1 | Egypt | Domiaty | Olivieri et al., 2015 |
| | KT184463.1 | Egypt | Domiaty | Olivieri et al., 2015 |
| | KT184456.1 | Egypt | Menofi | Olivieri et al., 2015 |
| | KT184457.1 | Egypt | Domiaty | Olivieri et al., 2015 |
| | KT184451.1 | Egypt | Menofi | Olivieri et al., 2015 |
| | KT184452.1 | Egypt | Domiaty | Olivieri et al., 2015 |
| | V00654.1 | NA | NA | Anderson et al., 1982 |
| | EU177862.1 | Italy | Valdostana | Achilli et al., 2008 |

| | | | | |
|---|---|---|---|---|
| Haplogroup R - Bos taurus | FJ971084.1 | Italy | Agerolese | Achilli et al., 2009 |
| | FJ971085.1 | Italy | Cinisara | Achilli et al., 2009 |
| | HQ184040.1 | Italy | Romagnola | Bonfiglio et al., 2010 |
| | FJ971087.1 | Italy | Romagnola | Achilli et al., 2009 |
| Haplogroup Q - Bos taurus | FJ971082.1 | Italy | Italian Red Pied | Achilli et al., 2009 |
| | FJ971083.1 | Italy | Romagnola | Achilli et al., 2009 |
| | HQ184036.1 | Italy | Grey Alpine | Bonfiglio et al., 2010 |
| | HQ184039.1 | Italy | Chianina | Bonfiglio et al., 2010 |
| | KT184471.1 | Egypt | Domiaty | Olivieri et al., 2015 |
| | KT184472.1 | Egypt | Domiaty | Olivieri et al., 2015 |
| | FJ971081.1 | Italy | Chianina | Achilli et al., 2009 |
| | HQ184030.1 | Italy | Chianina | Bonfiglio et al., 2010 |
| Haplogroup P - Bos primigenius | GU985279.1 | England - (6.7 kyr BP) | European aurochs | Edwards et al. 2010 |
| | JQ437479.1 | Poland | White Park Cattle | Ludwig et al. 2013 |
| | LC537309.1 | North-East Asia | Japanese short - horn | Mannen et al. 2020 |
| | LC537310.1 | North-East Asia | Japanese short - horn | Mannen et al. 2020 |
| | LC537317.1 | North-East Asia | Japanese short - horn | Mannen et al. 2020 |

| | | | | |
|---|---|---|---|---|
| Bos indicus - outgroup | NC_005971.1 (AY126697) | NA | Nelore | NCBI curated |
| | EU177869.1 | Iraq | Iraqi | Achilli et al. 2008 |

**Supplementary Table 4. Accessions, origin and references for the modern mitochondrial genomes of Bos and Ovis used in the phylogenetic analyses.**

**Reference**


1. Achilli, A., Bonfiglio, S., Olivieri, A., Malusa, A., Pala, M., Kashani, B. H., Perego, U. A., Ajmone-Marsan, P., Liotta, L. and Semino, O. 2009. The multifaceted origin of taurine cattle reflected by the mitochondrial genome. PLoS One 4(6), e5753.

2. Bonfiglio, S., Achilli, A., Olivieri, A., Negrini, R., Colli, L., Liotta, L., Ajmone-Marsan, P., Torroni, A. and Ferretti, L. 2010. The enigmatic origin of bovine mtDNA haplogroup R: Sporadic interbreeding or an independent event of Bos primigenius domestication in Italy? PLoS One 5(12), e15760.

3. Olivieri, A., Gandini, F., Achilli, A., Fichera, A., Rizzi, E., Bonfiglio, S., Battaglia, V., Brandini, S., De Gaetano, A., El-Beltagi, A. and Lancioni, H., 2015. Mitogenomes from Egyptian cattle breeds: new clues on the origin of haplogroup Q and the early spread of Bos taurus from the Near East. PLoS One, 10(10), p.e0141170.

4. Achilli, A., Olivieri, A., Pellecchia, M., Uboldi, C., Colli, L., Al-Zahery, N., Accetturo, M., Pala, M., Kashani, B.H., Perego, U.A. and Battaglia, V., 2008. Mitochondrial genomes of extinct aurochs survive in domestic cattle. Current Biology, 18(4), pp.R157-R158.

5. Mannen, Hideyuki, Takahiro Yonezawa, Kako Murata, Aoi Noda, Fuki Kawaguchi, Shinji Sasazaki, Anna Olivieri, Alessandro Achilli, and Antonio Torroni. 2020. "Cattle Mitogenome Variation Reveals a Post-Glacial Expansion of Haplogroup P and an Early Incorporation into Northeast Asian Domestic Herds." Scientific Reports 10 (1): 20842.

6.Edwards, C.J., Magee, D.A., Park, S.D., McGettigan, P.A., Lohan, A.J., Murphy, A., Finlay, E.K., Shapiro, B., Chamberlain, A.T., Richards, M.B. and Bradley, D.G., 2010. A complete mitochondrial genome sequence from a mesolithic wild aurochs (Bos primigenius). PloS one, 5(2), p.e9255.

7. Ludwig, A., Alderson, L., Fandrey, E., Lieckfeldt, D., Soederlund, T.K. and Froelich, K., 2013. Tracing the genetic roots of the indigenous White Park Cattle. Animal Genetics, 44(4), pp.383-386.

8. Taylor, William T. T., Mélanie Pruvost, Cosimo Posth, William Rendu, Maciej T. Krajcarz, Aida Abdykanova, Greta Brancaleoni, et al. 2021. "Evidence for Early Dispersal of Domestic Sheep into Central Asia." Nature Human Behaviour 5 (9): 1169–79.

9. Meadows, J., Hiendleder, S. & Kijas, J., 2011. Haplogroup relationships between domestic and wild sheep resolved using a mitogenome panel. Heredity 106, 700–706.

10. Niu, L., Chen, X., Xiao, P., Zhao, Q., Zhou, J., Hu, J., Sun, H., Guo, J., Li, L., Wang, L. and Zhang, H., 2017. Detecting signatures of selection within the Tibetan sheep mitochondrial genome. Mitochondrial DNA Part A, 28(6), pp.801-809.

11. Sanna, D., Barbato, M., Hadjisterkotis, E., Cossu, P., Decandia, L., Trova, S., Pirastru, M., Leoni, G.G., Naitana, S., Francalacci, P. and Masala, B., 2015. The first mitogenome of the Cyprus mouflon (Ovis gmelini ophion): new insights into the phylogeny of the genus Ovis. PloS one, 10(12), p.e0144257.

12. Lancioni, H., Di Lorenzo, P., Ceccobelli, S., Perego, U.A., Miglio, A., Landi, V., Antognoni, M.T., Sarti, F.M., Lasagna, E. and Achilli, A., 2013. Phylogenetic relationships of three Italian merino-derived sheep breeds evaluated through a complete mitogenome analysis. PloS one, 8(9), p.e73712

13. Lv, F.H., Peng, W.F., Yang, J., Zhao, Y.X., Li, W.R., Liu, M.J., Ma, Y.H., Zhao, Q.J., Yang, G.L., Wang, F. and Li, J.Q., 2015. Mitogenomic meta-analysis identifies two phases of migration in the history of eastern Eurasian sheep. Molecular biology and evolution, 32(10), pp.2515-2533.

14. Brahi, O.H.D., Xiang, H., Chen, X., Farougou, S. and Zhao, X., 2015. Mitogenome revealed multiple postdomestication genetic mixtures of West African sheep. Journal of Animal Breeding and Genetics, 132(5), pp.399-405.

15. Mustafa, S.I., Schwarzacher, T. and Heslop-Harrison, J.S., 2018. Complete mitogenomes from Kurdistani sheep: abundant centromeric nuclear copies representing diverse ancestors. Mitochondrial DNA Part A, 29(8), pp.1180-1193.

16. Liu, J.B., Ding, X.Z., Guo, T.T., Yue, Y.J., Zeng, Y.F., Guo, X., Chu, M., Han, J.L., Feng, R.L., Sun, X.P. and Niu, C.E., 2016. The complete mitochondrial genome sequence of the wild Huoba

Tibetan sheep of the Qinghai-Tibetan Plateau in China. Mitochondrial DNA Part A, 27(6), pp.4689-4690

17.Fan, H., Zhao, F., Zhu, C., Li, F., Liu, J., Zhang, L., Wei, C. and Du, L., 2016. Complete mitochondrial genome sequences of Chinese indigenous sheep with different tail types and an analysis of phylogenetic evolution in domestic sheep. Asian-Australasian Journal of Animal Sciences, 29(5), p.631.

18.Colli, L., Lancioni, H., Cardinali, I., Olivieri, A., Capodiferro, M.R., Pellecchia, M., Rzepus, M., Zamani, W., Naderi, S., Gandini, F. and Vahidi, S.M.F., 2015. Whole mitochondrial genomes unveil the impact of domestication on goat matrilineal variability. BMC genomics, 16(1), pp.1-12.
