## Supplementary material for "Early Pastoralism in Central European Forests: Insights from Ancient Environmental Genomics": Dataset S5

| #rname | startpos | endpos | n reads | cov bases | coverage | mean depth | mean baseq | mean mapq | Mean_length | D_max | lambdaLR | Sample ID |
|---|---|---|---|---|---|---|---|---|---|---|---|---|
| Bos_consensus | 1 | 16347 | 172 | 5707 | 34.9116 | 0.464183 | 39.7 | 34.6 | 44.13 | 0.12138755 | 26,257 | VM-17 |
| Bos_consensus | 1 | 16347 | 113 | 4803 | 29.3815 | 0.36496 | 39.6 | 36.7 | 52.79 | 0.092010096 | 56,453 | VM-15 |
| Bos_consensus | 1 | 16347 | 58 | 2177 | 13.3174 | 0.147428 | 39.5 | 36.8 | 41.58 | 0.08079707 | 17,284 | VM-14 |
| Bos_consensus | 1 | 16347 | 50 | 2221 | 13.5866 | 0.144675 | 39.3 | 36.3 | 47.29 | 0.17095686 | 11,150 | VM-19 |
| Ovis_consensus | 1 | 16772 | 110 | 4248 | 25.3279 | 0.363284 | 39.6 | 36.9 | 55.39 | 0.117461324 | 55.391 | VM-17 |
| Ovis_consensus | 1 | 16772 | 14 | 598 | 3.56547 | 0.038457 | 40.2 | 36.1 | 46.09 | 0.06694059 | 0.952 | VM-19 |

**Supplementary Table 5 (sheet 1). metaDMG output of the global damage estimates of the reads mapping to the consensus sequences used for constructing the phylogenetic tree.**

| pathphynder SNPs - Ovis | | | | | | | | | |
|---|---|---|---|---|---|---|---|---|---|
| Type of SNPs | Branch | Node | Position | Mutation | snp_count | Reads | Type of mutation | Extremity of read | Sample_ID |
| Supporting snps | 14 | 20 | 7446 | T->C | 14 | 2 | Transition | no | VM-19 |

| | | | | | | | | | |
|---|---|---|---|---|---|---|---|---|---|
| Conflicting snps | 2 | 14 | 16181 | G>A | 2 | 1 | Transition | no | VM-19 |
| Supporting snps | 14 | 20 | 13590 | C->T | 14 | 2 | Transition | no | VM-17 |
| | 14 | 20 | 6278 | C->T | | 1 | Transition | no | VM-17 |
| | 14 | 20 | 10130 | A->G | | 1 | Transition | no | VM-17 |
| | 14 | 20 | 11848 | C->T | | 4 | Transition | no | VM-17 |
| | 14 | 20 | 3673 | A->G | | 1 | Transition | no | VM-17 |
| Conflicting snps | 14 | 20 | 8132 | C->T | 14 | 1 | Transition | no | VM-17 |
| | 14 | 20 | 11329 | A->G | | 1 | Transition | no | VM-17 |
| Conflicting (Musimon snp) | 15 | 7 | 15459 | T->C | 15 | 1 | Transition | no | VM-17 |
| **pathphynder SNPs - Bos** | | | | | | | | | |
| Type of SNPs | Branch | Node | Position | Mutation | snp_count | Reads | Type of mutation | Extremity of read | Sample_ID |
| Supporting snps - origin node Q-T | 4 | 9 | 3558 | A->G | 13 | 1 | Transition | no | VM-14 |
| | 4 | 9 | 10699 | G->C | | 1 | Transversion | no | VM-14 |
| Conflicting snps - | 6 - Q | 3 | 8326 | T->C | 9 | 1 | Transition | no | VM-14 |

| | | | | | | | | | |
|---|---|---|---|---|---|---|---|---|---|
| origin node Q-T | 6 - Q | 3 | 3423 | A->T | | 1 | Transversion | no | VM-14 |
| | 5 - T | 2 | 5509 | T->C | 14 | 1 | Transition | no | VM-14 |
| | 5 - T | 2 | 16121 | T->C | | 1 | Transition | no | VM-14 |
| Supporting snps | 5 | 2 | 2566 | A > G | 14 | 1 | Transition | no | VM-15 |
| | 5 | 2 | 15142 | T > C | | 1 | Transition | no | VM-15 |
| Conflicting snps | 5 | 2 | 5509 | T > C | 14 | 1 | Transition | no | VM-15 |
| | 5 | 2 | 16121 | T > C | | 1 | Transition | no | VM-15 |
| Origin Node (Taurus Q conflict) | 6 | 3 | 3423 | A->T | 9 | 2 | Transversion | no | VM-15 |
| Supporting snps(base P) | 3 | 8 | 12809 | A->G | 35 | 1 | Transition | no | VM-17 |
| | 3 | 8 | 15625 | A->G | | 1 | Transition | no | VM-17 |
| | 3 | 8 | 16264 | T->C | | 2 | Transition | no | VM-17 |
| | 3 | 8 | 3447 | G->A | | 2 | Transition | no | VM-17 |
| | 3 | 8 | 5622 | A->G | | 1 | Transition | 1st from end | VM-17 |

| | | | | | | | | | |
|---|---|---|---|---|---|---|---|---|---|
| Conflicting snps (base P) | 3 | 8 | 3608 | T->C | 35 | 1 | Transition | no | VM-17 |
| Conflicting Bos P | 7 | 4 | 10134 | C->T | 36 | 1 | Transition | no | VM-17 |
| | 7 | 4 | 11148 | A->G | | 1 | Transition | no | VM-17 |
| | 7 | 4 | 12024 | T->C | | 1 | Transition | no | VM-17 |
| | 7 | 4 | 4260 | T->C | | 2 | Transition | no | VM-17 |
| | 7 | 4 | 4301 | C->T | | 1 | Transition | no | VM-17 |
| | 7 | 4 | 4684 | A->G | | 1 | Transition | no | VM-17 |
| | 7 | 4 | 5907 | A->G | | 1 | Transition | no | VM-17 |
| | 7 | 4 | 16273 | A->G | | 2 | Transition | no | VM-17 |
| Conflicting Bos Q | 6 | 3 | 3246 | T->C | 9 | 1 | Transition | no | VM-17 |
| | 6 | 3 | 3423 | A->T | | 1 | Transversion | no | VM-17 |
| | 6 | 3 | 7926 | C->T | | 1 | Transition | no | VM-17 |
| | 6 | 3 | 10935 | C->T | | 2 | Transition | no | VM-17 |
| Conflicting Bos T | 5 | 2 | 15635 | G->A | 14 | 1 | Transition | no | VM-17 |
| | 5 | 2 | 2566 | A->G | | 1 | Transition | no | VM-17 |

**Supplementary Table 5 (sheet 2). Pathphynder SNPs count.**
