## Supplementary material for "Early Pastoralism in Central European Forests: Insights from Ancient Environmental Genomics": Dataset S6

| Run access | Assay Type | BioProject | BioSample | Type of source |
|---|---|---|---|---|
| SRR6192418 | WGS | PRJNA366548 | SAMN06268883 | salt_marsh_metagenome |
| ERR257713 | WGS | PRJEB1760 | SAMEA2062903 | salt_marsh_metagenome |
| ERR257714 | WGS | PRJEB1760 | SAMEA2062904 | salt_marsh_metagenome |
| ERR257715 | WGS | PRJEB1760 | SAMEA2062906 | salt_marsh_metagenome |
| ERR257716 | WGS | PRJEB1760 | SAMEA2062905 | salt_marsh_metagenome |
| ERR1725642 | WGS | PRJEB14421 | SAMEA4537330 | freshwater_sediment_lake |
| ERR1725798 | WGS | PRJEB14421 | SAMEA4542491 | freshwater_sediment_lake |
| ERR1725848 | WGS | PRJEB14421 | SAMEA4544563 | freshwater_sediment_lake |
| ERR1725849 | WGS | PRJEB14421 | SAMEA4544564 | freshwater_sediment_lake |
| ERR1725853 | WGS | PRJEB14421 | SAMEA4544646 | freshwater_sediment_lake |
| ERR1960500 | WGS | PRJEB20765 | SAMEA104055711 | permafrost |
| ERR1960501 | WGS | PRJEB20765 | SAMEA104055712 | permafrost |
| ERR1960503 | WGS | PRJEB20765 | SAMEA104055714 | permafrost |

| | | | | |
|---|---|---|---|---|
| ERR1960504 | WGS | PRJEB20765 | SAMEA104055715 | permafrost |
| ERR1960505 | WGS | PRJEB20765 | SAMEA104055716 | permafrost |
| ERR1017187 | WGS | PRJEB10725 | SAMEA3539301 | melting_permafrost_tundra |
| ERR1019366 | WGS | PRJEB10725 | SAMEA3539303 | melting_permafrost_tundra |
| ERR1022686 | WGS | PRJEB10725 | SAMEA3539305 | melting_permafrost_tundra |
| ERR1022687 | WGS | PRJEB10725 | SAMEA3539304 | melting_permafrost_tundra |
| ERR1022692 | WGS | PRJEB10725 | SAMEA3539306 | melting_permafrost_tundra |
| ERR2486618 | WGS | PRJEB22376 | SAMEA104335343 | agricultural_soil |
| ERR2486619 | WGS | PRJEB22376 | SAMEA104335344 | agricultural_soil |
| ERR2486620 | WGS | PRJEB22376 | SAMEA104335345 | agricultural_soil |
| ERR2486621 | WGS | PRJEB22376 | SAMEA104335346 | agricultural_soil |
| ERR2486622 | WGS | PRJEB22376 | SAMEA104335347 | agricultural_soil |
| ERR1700674 | WGS | PRJEB8420 | SAMEA4521324 | temperate_forest_soil_mineral_layer |

| | | | | |
|---|---|---|---|---|
| ERR1700677 | WGS | PRJEB8420 | SAMEA4521327 | temperate_forest_soil_organic_layer |
| ERR1700680 | WGS | PRJEB8420 | SAMEA4521330 | temperate_forest_soil_mineral_layer |
| ERR1700683 | WGS | PRJEB8420 | SAMEA4521333 | temperate_forest_soil_organic_layer |
| ERR1700686 | WGS | PRJEB8420 | SAMEA4521336 | temperate_forest_soil_mineral_layer |
| ERR476938 | WGS | PRJEB5872 | SAMEA2464834 | wetland_acidic_soil |
| ERR476939 | WGS | PRJEB5872 | SAMEA2464835 | wetland_acidic_soil |
| ERR476940 | WGS | PRJEB5872 | SAMEA2464835 | wetland_acidic_soil |
| ERR476941 | WGS | PRJEB5872 | SAMEA2464836 | wetland_acidic_soil |
| ERR476942 | WGS | PRJEB5872 | SAMEA2464837 | wetland_acidic_soil |
| ERR1041384 | WGS | PRJEB10725 | SAMEA3539312 | temperate_prairie_grassland |
| ERR1041385 | WGS | PRJEB10725 | SAMEA3539313 | temperate_prairie_grassland |

| | | | | |
|---|---|---|---|---|
| ERR1043165 | WGS | PRJEB10725 | SAMEA3539314 | temperate_prairie_grassland |
| ERR1043166 | WGS | PRJEB10725 | SAMEA3539315 | temperate_prairie_grassland |
| ERR1044071 | WGS | PRJEB10725 | SAMEA3539317 | temperate_prairie_grassland |
| ERR249373 | WGS | PRJEB1725 | SAMEA2060399 | tropical_forest_soil |
| ERR249374 | WGS | PRJEB1725 | SAMEA2060400 | tropical_forest_soil |
| ERR249375 | WGS | PRJEB1725 | SAMEA1965874 | tropical_forest_soil |
| ERR249376 | WGS | PRJEB1725 | SAMEA2060401 | tropical_forest_soil |
| ERR249377 | WGS | PRJEB1725 | SAMEA2060402 | tropical_forest_soil |
| ERR1357129 | WGS | PRJEB13455 | SAMEA3928102 | biocrust |
| ERR1357130 | WGS | PRJEB13455 | SAMEA3928103 | biocrust |
| ERR1358721 | WGS | PRJEB13455 | SAMEA3928104 | biocrust |
| ERR1358722 | WGS | PRJEB13455 | SAMEA3928105 | biocrust |
| ERR1358723 | WGS | PRJEB13455 | SAMEA3928105 | biocrust |
| ERR1337728 | WGS | PRJEB13142 | SAMEA3905791 | desert_varnish |

| | | | | |
|---|---|---|---|---|
| ERR1337812 | WGS | PRJEB13142 | SAMEA3905791 | desert_varnish |
| ERR1341893 | WGS | PRJEB13142 | SAMEA3905791 | desert_varnish |
| ERR1351806 | WGS | PRJEB13142 | SAMEA3905791 | desert_varnish |
| ERR1352472 | WGS | PRJEB13142 | SAMEA3905793 | desert_varnish |
| ERR1352473 | WGS | PRJEB13142 | SAMEA3905793 | desert_varnish |
| ERR2191879 | WGS | PRJEB23251 | SAMEA104369071 | desert_fairy_circles |
| ERR2191880 | WGS | PRJEB23251 | SAMEA104369072 | desert_fairy_circles |
| ERR2191881 | WGS | PRJEB23251 | SAMEA104369073 | desert_fairy_circles |
| ERR2191958 | WGS | PRJEB23251 | SAMEA104369074 | desert_fairy_circles |
| ERR2191959 | WGS | PRJEB23251 | SAMEA104369075 | desert_fairy_circles |
| ERR481108 | WGS | PRJEB6137 | SAMEA2468441 | river_lime_clay |
| ERR481109 | WGS | PRJEB6137 | SAMEA2468442 | river_lime_clay |
| ERR481110 | WGS | PRJEB6137 | SAMEA2468443 | river_lime_clay |
| ERR481111 | WGS | PRJEB6137 | SAMEA2468444 | river_lime_clay |
| ERR481112 | WGS | PRJEB6137 | SAMEA2468445 | river_lime_clay |
| ERR059346 | WGS | PRJEB2790 | SAMEA1569094 | Soil_low_PH |

| | | | | |
|---|---|---|---|---|
| ERR059347 | WGS | PRJEB2790 | SAMEA1569093 | Soil_low_PH |
| ERR059348 | WGS | PRJEB2790 | SAMEA1569087 | Soil_low_PH |
| ERR059349 | WGS | PRJEB2790 | SAMEA1569089 | Soil_low_PH |
| ERR1755753 | WGS | PRJEB18597 | SAMEA26292418 | soil_pH.4.5_agricultural |
| ERR1755754 | WGS | PRJEB18597 | SAMEA26293168 | soil_pH.4.5_agricultural |
| ERR1755755 | WGS | PRJEB18597 | SAMEA26293918 | soil_pH.5.0_agricultural |
| ERR1755756 | WGS | PRJEB18597 | SAMEA26294668 | soil_pH.5.0_agricultural |
| ERR2200466 | WGS | PRJEB23561 | SAMEA104395147 | Bovine_Rumen |
| ERR2200467 | WGS | PRJEB23561 | SAMEA104395148 | Bovine_Rumen |
| ERR2200468 | WGS | PRJEB23561 | SAMEA104395149 | Bovine_Rumen |
| ERR2200469 | WGS | PRJEB23561 | SAMEA104395150 | Bovine_Rumen |
| SRR094166 | WGS | PRJNA60251 | SAMN00149621 | Cow_Rumen |
| SRR094403 | WGS | PRJNA60251 | SAMN00149621 | Cow_Rumen |
| SRR094405 | WGS | PRJNA60251 | SAMN00149621 | Cow_Rumen |
| SRR094415 | WGS | PRJNA60251 | SAMN00149621 | Cow_Rumen |
| SRR094416 | WGS | PRJNA60251 | SAMN00149621 | Cow_Rumen |
| ERR571345 | WGS | PRJEB6883 | SAMEA2673752 | Bos_taurus_dung |

| | | | | |
|---|---|---|---|---|
| ERR571346 | WGS | PRJEB6883 | SAMEA2673753 | Bos_taurus_dung |
| ERR571347 | WGS | PRJEB6883 | SAMEA2673754 | Bos_taurus_dung |
| ERR571348 | WGS | PRJEB6883 | SAMEA2673755 | Bos_taurus_dung |
| ERR571349 | WGS | PRJEB6883 | SAMEA2673756 | Bos_taurus_dung |
| ERR1135178 | WGS | PRJEB11755 | SAMEA3663005 | pig_fecal |
| ERR1135179 | WGS | PRJEB11755 | SAMEA3663006 | pig_fecal |
| ERR1135180 | WGS | PRJEB11755 | SAMEA3663007 | pig_fecal |
| ERR1135181 | WGS | PRJEB11755 | SAMEA3663008 | pig_fecal |
| ERR1135182 | WGS | PRJEB11755 | SAMEA3663009 | pig_fecal |
| ERR1447687 | WGS | PRJEB14312 | SAMEA4028452 | Ovis_aries_fecal |
| ERR1448096 | WGS | PRJEB14312 | SAMEA4028453 | Ovis_aries_fecal |
| ERR1448097 | WGS | PRJEB14312 | SAMEA4028454 | Ovis_aries_fecal |
| ERR1448098 | WGS | PRJEB14312 | SAMEA4028455 | Ovis_aries_fecal |
| ERR1448099 | WGS | PRJEB14312 | SAMEA4028456 | Ovis_aries_fecal |
| SRR1206671 | WGS | PRJNA202380 | SAMN02144231 | Sheep rumen microbiome |
| SRR1222429 | WGS | PRJNA202380 | SAMN02144230 | Sheep rumen microbiome |

| | | | | |
|---|---|---|---|---|
| SRR1222431 | WGS | PRJNA202380 | SAMN02144231 | Sheep rumen microbiome |
| SRR1267595 | WGS | PRJNA202380 | SAMN02144230 | Sheep rumen microbiome |
| SRR873595 | WGS | PRJNA202380 | SAMN02144230 | Sheep rumen microbiome |
| ERR2806347 | WGS | PRJEB28701 | SAMEA4931674 | Homo_sapiens_fecal |
| ERR2806348 | WGS | PRJEB28701 | SAMEA4931675 | Homo_sapiens_fecal |
| ERR2806349 | WGS | PRJEB28701 | SAMEA4931676 | Homo_sapiens_fecal |
| ERR2806350 | WGS | PRJEB28701 | SAMEA4931677 | Homo_sapiens_fecal |
| ERR2806351 | WGS | PRJEB28701 | SAMEA4931678 | Homo_sapiens_fecal |

**Supplementary Table 6. Accessions for the microbial dataset used in Sourcetracker2 with indication of the type of source.**
