## Supplementary material for "Early Pastoralism in Central European Forests: Insights from Ancient Environmental Genomics": Dataset S7

| Sample ID | Bovine rumen | Sheep fecal | Pig fecal | Human fecal | Temperate forest | Wetland acidic_soil | River lime_clay | Temperate grassland | Other soil | Unknown | taxon count | read count |
|---|---|---|---|---|---|---|---|---|---|---|---|---|
| VM-2 | 0.001 | 0 | 0 | 0 | 0.001 | 0 | 0 | 0.012 | 0.009 | 0.977 | 8 | 634 |
| VM-3 | 0 | 0 | 0 | 0 | 0 | 0 | 0 | 0 | 0 | 1 | 4 | 346 |
| VM-11 | 0.003 | 0.02 | 0.768 | 0.001 | 0 | 0 | 0.001 | 0.001 | 0.008 | 0.198 | 37 | 6265 |
| VM-14 | 0.005 | 0.05 | 0.013 | 0.001 | 0.003 | 0.005 | 0 | 0.006 | 0.061 | 0.856 | 32 | 953 |
| VM-15 | 0.019 | 0.002 | 0 | 0.001 | 0 | 0 | 0 | 0.001 | 0.004 | 0.972 | 141 | 119027 |
| VM-17 | 0.072 | 0.058 | 0 | 0.001 | 0.001 | 0 | 0.003 | 0.001 | 0.062 | 0.78 | 100 | 4921 |
| VM-19 | 0.012 | 0.055 | 0.001 | 0 | 0.007 | 0.009 | 0.001 | 0.027 | 0.032 | 0.853 | 54 | 3468 |
| VM-22 | 0.002 | 0 | 0.002 | 0.001 | 0 | 0 | 0.002 | 0 | 0.003 | 0.984 | 36 | 71593 |
| VM-24 | 0 | 0.001 | 0 | 0.001 | 0 | 0.028 | 0 | 0.001 | 0.011 | 0.957 | 18 | 18857 |
| VM-26 | 0 | 0 | 0 | 0 | 0 | 0 | 0 | 0 | 0 | 1 | 6 | 630 |
| VM-28 | 0 | 0 | 0 | 0 | 0 | 0 | 0 | 0 | 0.001 | 0.999 | 3 | 235 |

**Supplementary Table 7 (sheet 1). Results of Sourcetracker2 for reads with a post-mortem DNA damage above 5%.**

| Sample ID | Bovine rumen | Sheep fecal | Pig fecal | Human fecal | Temperate forest | Wetland acidic_soil | River lime_clay | Temperate grassland | Other soil | Unknown | Taxon count | Read count |
|---|---|---|---|---|---|---|---|---|---|---|---|---|
| VM-2 | 0 | 0 | 0.002 | 0.001 | 0.011 | 0.002 | 0 | 0.009 | 0.01 | 0.965 | 163 | 13543 |
| VM-3 | 0.002 | 0 | 0.001 | 0 | 0.011 | 0.004 | 0 | 0.035 | 0.138 | 0.801 | 425 | 56568 |
| VM-11 | 0.091 | 0.029 | 0.032 | 0.001 | 0 | 0 | 0 | 0.022 | 0.058 | 0.763 | 98 | 3072 |
| VM-14 | 0.003 | 0.024 | 0.005 | 0.003 | 0.004 | 0.012 | 0.009 | 0.043 | 0.106 | 0.79 | 455 | 14118 |
| VM-15 | 0.023 | 0.009 | 0.001 | 0 | 0.001 | 0.001 | 0.001 | 0.022 | 0.056 | 0.885 | 325 | 20621 |
| VM-17 | 0.012 | 0.002 | 0.001 | 0 | 0.01 | 0.001 | 0 | 0.012 | 0.046 | 0.915 | 337 | 17400 |
| VM-19 | 0.012 | 0 | 0.001 | 0 | 0 | 0.007 | 0.001 | 0 | 0.026 | 0.95 | 199 | 22376 |

| | | | | | | | | | | | |
|---|---|---|---|---|---|---|---|---|---|---|---|
| VM-22 | 0.006 | 0.001 | 0 | 0.003 | 0.002 | 0.002 | 0.002 | 0.002 | 0.027 | 0.955 | 312 | 63195 |
| VM-24 | 0 | 0.011 | 0 | 0 | 0.002 | 0.005 | 0.001 | 0.001 | 0.064 | 0.916 | 124 | 14894 |
| VM-26 | 0.01 | 0.002 | 0 | 0.001 | 0.001 | 0 | 0 | 0 | 0.01 | 0.976 | 199 | 100138 |
| VM-28 | 0.018 | 0.002 | 0 | 0.001 | 0.001 | 0.002 | 0.002 | 0.003 | 0.081 | 0.888 | 706 | 114648 |

**Supplementary Table 7 (sheet 2). Results of Sourcetracker2 for reads with a post-mortem DNA damage below 5%.**
